# Atlas-Guided Cell-Specific Transcriptomics Identifies Pathway-Level Remodeling of Human Hypothalamic CRH Neurons in Opioid Use Disorder

**DOI:** 10.64898/2026.09.09.749970

**Authors:** Lorna Barrall, Phillip Urbanczyk, Bruno Kluwe-Schiavon, Laura Stertz, Sakuni Rankothgedera, Micah Castillo, Preethi Gunaratne, Consuelo Walss-Bass

## Abstract

Opioid use disorder (OUD) disrupts hypothalamic stress signaling, yet the molecular state of human paraventricular nucleus (PVN) corticotropin-releasing hormone (CRH) neurons remains poorly defined. Using single-nucleus RNA sequencing of 343,819 human hypothalamic nuclei, we applied complementary human HYPOMAP/MapMyCells and mouse PVN Atlas strategies that independently converged on the standardized Allen Brain Map subcluster Splat_410_843, providing cross-atlas validation of a rare CRH-enriched PVN population. Integrated GO, Reactome and KEGG analysis revealed OUD-associated enrichment of synaptic, junctional, cytoskeletal, receptor-signaling and ion-transport programs, with relative enrichment of RNA-processing, ER–Golgi/vesicular, endolysosomal and glycan-related functions in controls. Targeted neuropeptide analysis further identified nominal increases in OXT, GHR, GAL and PRLR and decreases in VIP, AGRP, NPY, CRHBP and CALCR. Together, these findings define a reproducible human PVN CRH population and identify coordinated pathway and neuropeptide remodeling in OUD.

## Introduction

Opioid use disorder (OUD) is a significant public health problem and deaths due to overdose remain high, underscoring the importance of understanding the mechanisms of opioid-induced brain alterations. Most studies in human brain in OUD have focused on the dorsolateral prefrontal cortex and the nucleus accumbens, identifying neuroinflammation and synaptic-remodeling signatures (Green et al. 2026 [preprint], Seney et al., 2021), while other important brain regions such as the hypothalamus have remained largely unexplored.

Chronic opioid use and OUD produce bidirectional dysregulation of the hypothalamic-pituitary-adrenal (HPA) axis ***<u>(Figure 1)</u>***, the primary neuroendocrine system that mediates the physiological stress response. The HPA axis operates through three core components: corticotropin-releasing hormone (CRH) neurons in the paraventricular nucleus (PVN) of the hypothalamus, corticotropes of the anterior pituitary, and the adrenal cortex. PVN CRH neurons initiate this cascade by releasing CRH and arginine vasopressin (AVP) into the hypophyseal portal system, driving adrenocorticotropic hormone (ACTH) secretion at the pituitary and glucocorticoid synthesis at the adrenals, which in turn exert negative feedback at both hypothalamic and pituitary levels (Herman et al., 2016; Chrousos, 1995). During active opioid exposure, mu-, delta-, and kappa-opioid receptors exert inhibitory effects on PVN CRH neurons, suppressing CRH and AVP secretion and reducing ACTH release, leading to hypocortisolism. Systematic review and meta-analysis data indicate that approximately 15–24% of chronic opioid users develop biochemical hypocortisolism, with cumulative opioid exposure the strongest predictor (de Vries et al., 2020), and opioid-induced adrenal insufficiency prevalence estimated at ∼9% in cross-sectional studies (Li et al., 2020). Evidence from human longitudinal studies suggests these perturbations may be reversible with sustained abstinence: in a 20-year heroin-dependence cohort, no opioid-free participant had hypocortisolism, versus 14.8% of those with continuing opioid use (Tremonti et al., 2026).

**Figure 1.**
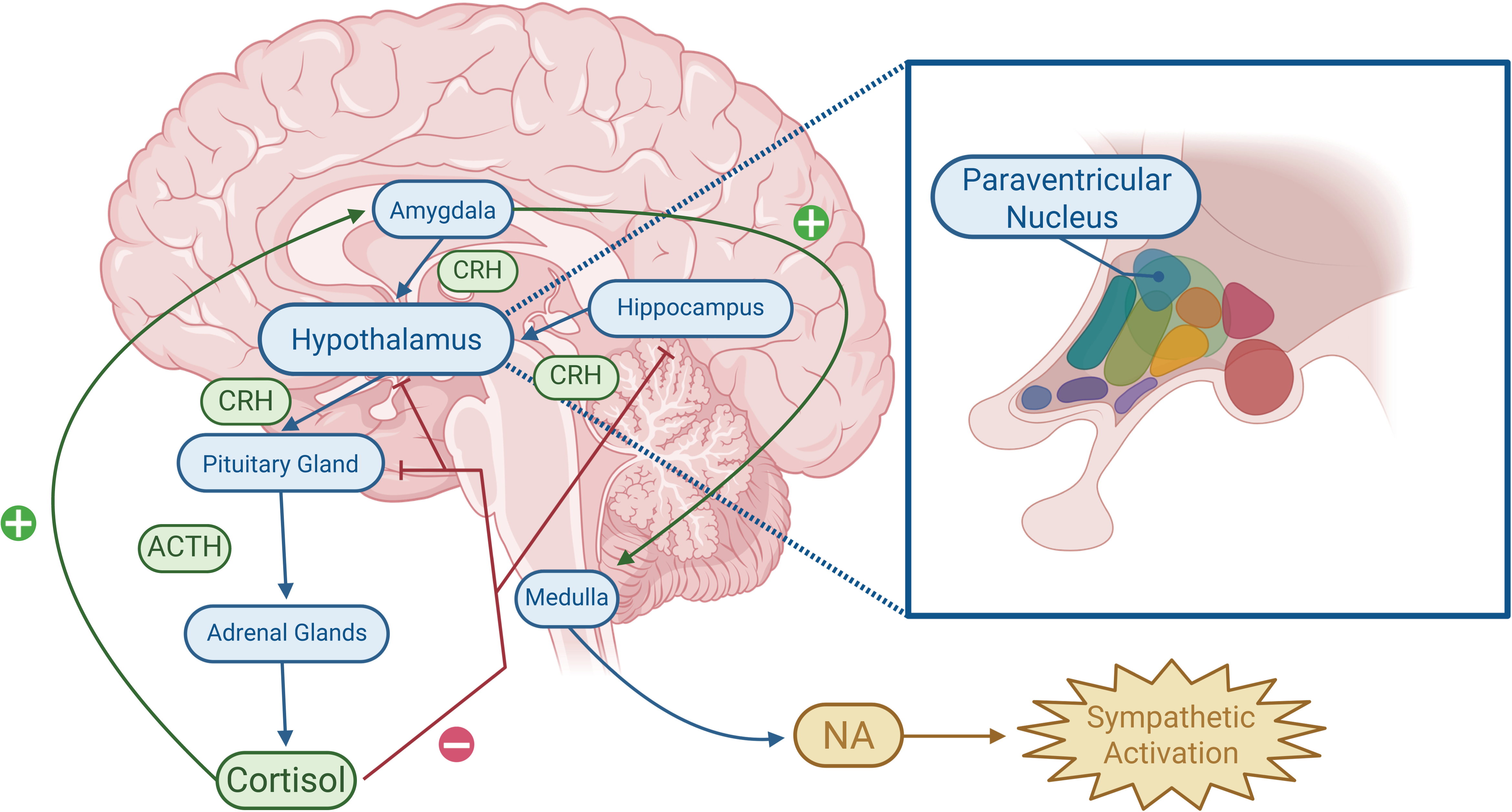
Proposed anti-reward opponent pathway in opioid dependence. Hypothalamic CRH activates the HPA axis, stimulating ACTH and cortisol release. Cortisol normally provides negative feedback to the hypothalamus (focusing in on the Paraventricular Nucleus), pituitary, and hippocampus. During opioid dependence, increased amygdala CRH signaling may overcome this feedback and activate noradrenergic stress pathways, contributing to sympathetic activation, stress, and negative affect. Adapted from Kakko et al. (2019).

PVN CRH neurons are conceptualized as central to the initial stages of drug use and the negative-reinforcement/withdrawal phase ***<u>(Figure 1),</u>*** while extrahypothalamic CRF populations in the CeA and BNST predominate in stress-induced relapse and protracted negative affect (Huang et al., 2024). However, direct human tissue data on PVN CRH neurons in OUD remain scarce. The most directly relevant human postmortem finding is a ∼20–27% reduction in total hypothalamic volume in male heroin addicts versus controls (Müller et al., 2018). The hypothalamus poses particular technical challenges: it is small and heterogeneous, requires rapid collection and specialized dissection, and CRH neurons constitute a minority of PVN cells. To our knowledge, no published study has directly examined CRH expression, neuron number, or transcriptomic profiles in the human PVN in the context of chronic opioid exposure or OUD.

This study characterized human hypothalamic CRH neurons in chronic opioid exposure using single-nucleus RNA sequencing (snRNA-seq) of postmortem tissue. Leveraging Allen MapMyCells (MMC), the human HYPOMAP atlas and the rodent PVN atlas as complementary references, this work provides the first direct cell-specific characterization of opioid-induced alteration of CRH neurons in the human PVN hypothalamus.

## Methods

### Human Subjects

Postmortem hypothalamic tissue was sourced from the University of Texas Health Science Center–Houston Brain Collection (UTHBC), in collaboration with the Harris County Institute of Forensic Sciences, with consent from next-of-kin (NOK) and approval by the Institutional Review Board. The cohort comprised 42 donors: 22 healthy controls (HC) and 20 donors with OUD ***(Supplementary Table 1),*** classified by consensus diagnosis from 3 independent clinicians, after review of available medical, autopsy, and toxicology reports, and the NOK interview using the UT Health Psychological Autopsy Interview Schedule (UTH-PAIS; Meyer et al., 2022). All OUD samples had confirmed chronic opioid use. Hypothalamic tissue was obtained from each donor via a 5-mm punch biopsy of flash-frozen coronal sections of the brain, guided by gross anatomical landmarks including third ventricle, thalamus and the mammillary bodies. Nuclei isolation, library preparation, and sequencing were performed at the University of Houston Bioinformatics Core (see Supplementary Methods).

### Sequencing, preprocessing, and quality control

Nuclei were processed using the 10x Genomics Chromium Fixed RNA Profiling (Flex) workflow and quantified with Cell Ranger 7.1.0, followed by mitochondrial filtering, targeted doublet removal, Harmony integration, and Louvain clustering. The final dataset comprised 343,819 nuclei. Full library-preparation details, read-alignment parameters, quality-control thresholds, doublet diagnostics, batch-mixing analyses, and clustering settings are provided in Supplementary Methods (“Library preparation and sequencing” through “Dimensionality reduction and clustering”).

### Cell type annotation

Allen Brain Institute MapMyCells (MMC). Cluster-to-cell-type assignment used the Allen Brain Institute MapMyCells (MMC) tool and its whole-brain reference taxonomy, providing supercluster, cluster, and subcluster identities with bootstrapping probability estimates. Superclusters with fewer than 100 cells were collapsed into "other" and were not included in further analysis. Thirty supercluster categories were represented, including glial/non-neuronal classes and a large set of neuronal categories ***<u>(Figure 2a)</u>*** among them a large, multi-region "Splatter" supercluster containing Splat_410, a hypothalamus-specific cluster (16,596 cells across 12 constituent subclusters) independently confirmed as anatomically hypothalamic via the reference taxonomy’s dissection-region metadata (66–98% hypothalamus per subcluster; Siletti et al., 2023). After excluding 7 donor-concentrated categories and 3 raw clusters (together 14,296 of 343,819 cells; 4.2%), 23 of 30 supercluster categories were retained. Broad neuronal/non-neuronal assignments and canonical and neuropeptide marker-panel validation are detailed in Supplementary Methods (“Neuronal and non-neuronal cell populations”).

**Figure 2.**
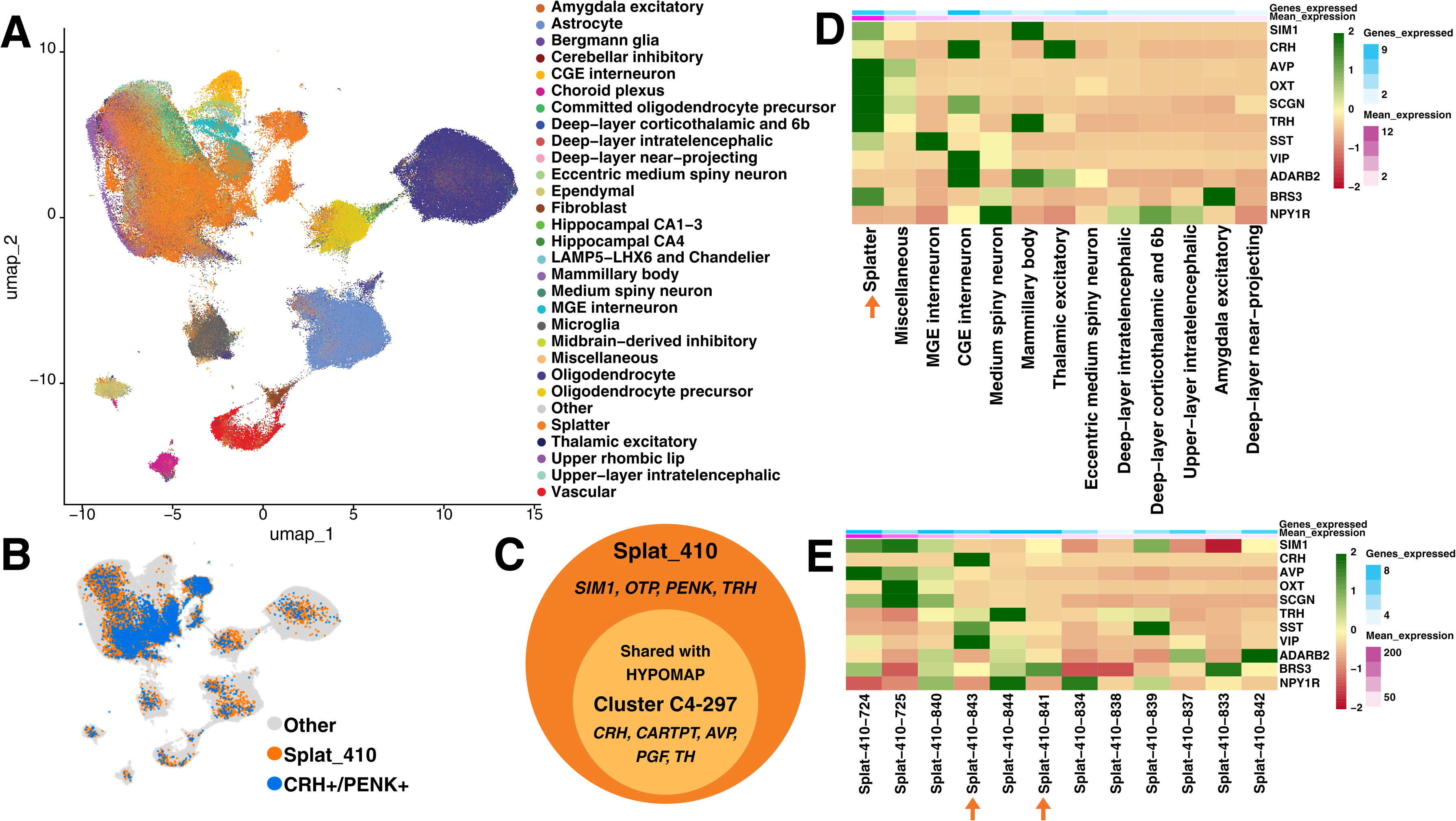
Atlas-guided identification and validation of CRH-enriched neurons in the human hypothalamus. **(A) Cell-type composition of the human hypothalamic snRNA-seq dataset.** UMAP visualization of 343,819 nuclei annotated using Allen Brain Institute MapMyCells, identifying 30 neuronal and non-neuronal supercluster categories across the hypothalamic dataset. **(B) Localization of the** CRH+/PENK+ **Splat_410 subpopulation.** UMAP highlighting Splat_410 within the full dataset and the CRH+/PENK+ subpopulation comprising Splat_410_841 and Splat_410_843. Other cell types are shown in grey, Splat_410 cells in orange, and CRH+/PENK+ cells in blue. **(C) Cross-atlas marker convergence between Splat_410 and HYPOMAP C4-297.** Nested-circle diagram showing that all five top C4-297 markers (CRH, CARTPT, AVP, PGF, and TH) are contained within the Splat_410 marker set, supporting cross-atlas concordance of the CRH-enriched population. **(D) PVN marker expression across neuronal cell types.** Heatmap showing scaled average expression of PVN/neuroendocrine marker genes across MMC neuronal superclusters. Red indicates lower, yellow average, and green higher relative expression for each gene. Top annotations indicate mean marker expression and the number of markers expressed per supercluster. Arrow indicated Splatter neurons. **(E) PVN marker expression across Splat_410 subclusters.** Heatmap showing scaled average expression of PVN-associated markers across Splat_410 subclusters. Splat_410_843 and Splat_410_841 showed the highest CRH expression.

### PVN CRH neuron identification

Two complementary atlas-guided strategies were used to identify PVN-resident CRH neurons: a human HYPOMAP/MapMyCells-guided definition of CRH+/PENK+ Splat_410 subclusters ***<u>(Figure 2)</u>*** and an independent mouse PVN Atlas-guided re-clustering of SIM1-positive nuclei ***<u>(Figure 3)</u>***. Both approaches converged on Splat_410_843, supporting the CRH-enriched identity used for downstream analysis. Detailed marker scoring, confidence criteria, atlas cross-validation, and species-specific comparisons are provided in Supplementary Methods (“CRH neuron identification”) and Supplementary Results (“Cross-atlas validation of the CRH-enriched population”).

**Figure 3.**
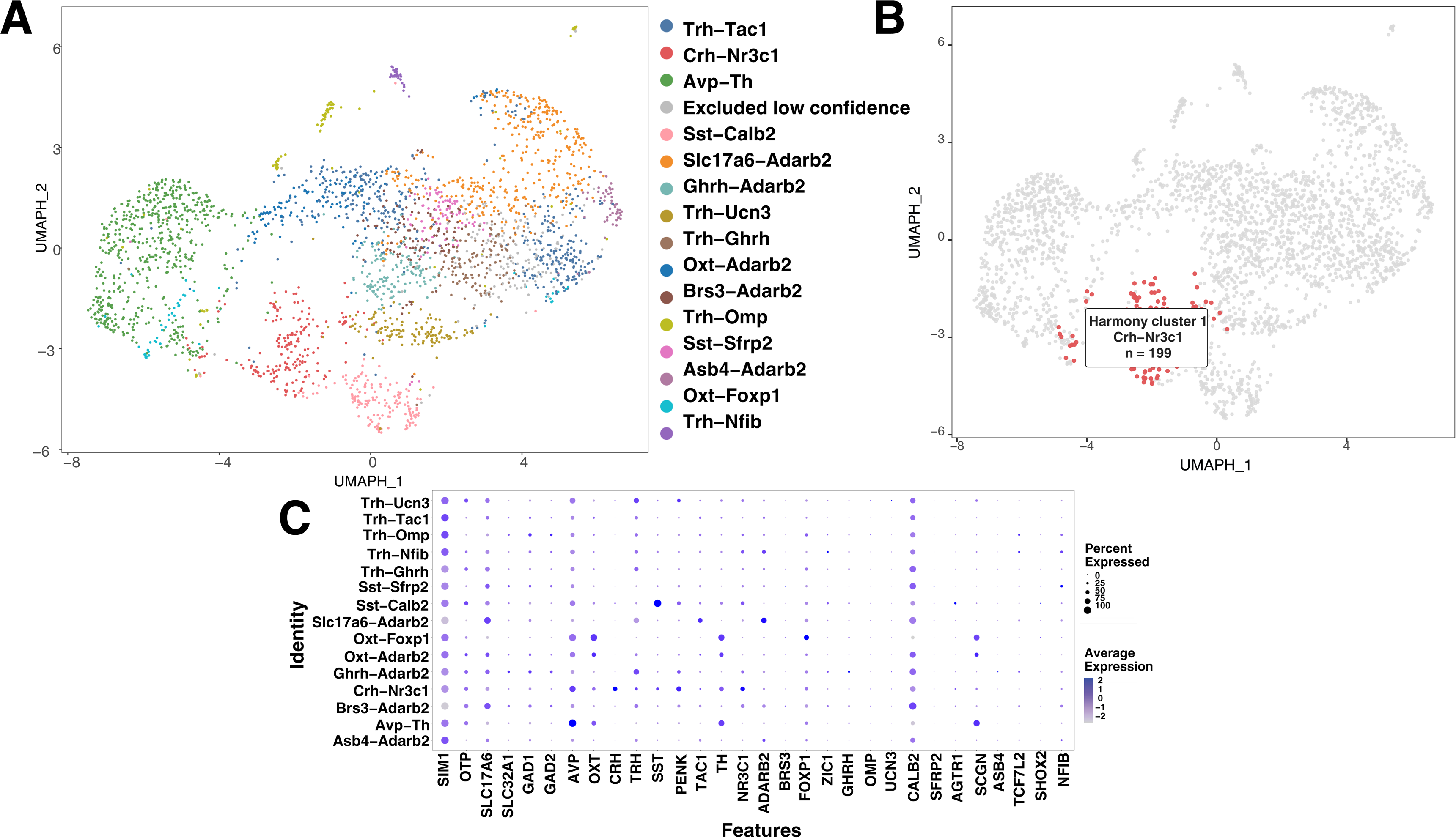
PVN Atlas-guided identification and validation of CRH-enriched SIM1-positive neuronal population. **(A) Atlas-guided identification of SIM1-positive PVN neuronal populations**. UMAP visualization of 2,847 SIM1-positive neurons following Harmony integration and unsupervised clustering. Post hoc biological labels were assigned using PVN Atlas-associated marker expression; the low-confidence population excluded from downstream label-specific analyses is shown in gray**. (B) Localization of the CRH-associated Harmony cluster.** UMAP of 2,847 SIM1-positive PVN neurons highlighting Harmony cluster 1 in red. This 199-cell cluster was assigned a high-confidence Crh-Nr3c1 biological label based on PVN Atlas-associated marker expression. **(C) PVN Atlas marker expression across SIM1-positive neuronal populations.** Dot plot showing selected PVN Atlas-associated marker genes across post hoc biological labels assigned to Harmony-derived clusters. Dot size indicates the percentage of cells expressing each marker, and color intensity indicates scaled average normalized expression.

Selection of population for differential expression. Because the HYPOMAP-guided pipeline (Pipeline 1) provided broader coverage (Splat_410_841 + Splat_410_843; ∼4,774 cells) and donor-level pseudobulk analysis benefits from larger per-donor cell counts, the Splat_410_841 and Splat_410_843 subcluster-membership definition was used for differential expression. The mouse-atlas pipeline’s convergent identification of Splat_410_843 served as independent validation that this population contains the core CRH-enriched PVN neurons.

### Differential expression analysis

Analytical design. Differential expression between OUD and HC was assessed using donor-level pseudobulk aggregation with DESeq2 (design: sex + age + PMI + brain pH + RIN + condition; apeglm shrinkage; HC as reference). Three tiers tested whether population-specific effects were diluted at broader aggregation levels: (1) general cell-type level (MMC supercluster classification); (2) the entire Splat_410 population (all 12 Splat_410 subclusters); and (3) the CRH+/PENK+ subpopulation (Splat_410_841 + Splat_410_843).

For the CRH+/PENK+ comparison, donors contributing fewer than 10 nuclei were excluded to avoid unstable pseudobulk profiles, retaining 25 donors (14 HC and 11 OUD). The rationale for pseudobulk modeling, donor representation, and sensitivity checks are provided in Supplementary Methods (“CRH+/PENK+ pseudobulk analysis”).

### Pathway analysis

Functional enrichment of the CRH+/PENK+ neuron differential-expression results was characterized across six complementary annotation sources: Gene Ontology (GO) Biological Process (BP), Cellular Component (CC), and Molecular Function (MF); KEGG; Reactome; and SynGO. For GO, KEGG, and Reactome, ranked gene set enrichment analysis (GSEA) was performed on the full gene ranking; for SynGO, over-representation analysis (ORA) was performed on thresholded foreground gene sets against a defined background. Each source was first analyzed independently and then integrated across sources using normalized enrichment score (NES), leading-edge gene overlap, and GO semantic similarity. Positive NES and log2 fold change denote higher enrichment or expression in OUD; negative values denote higher enrichment or expression in controls.

Database releases, software versions, ranking and threshold parameters, leading-edge gene processing, enrichment-map construction ***<u>(Figure 4)</u>***, GOChord ***<u>(Figure 5d)</u>*** and alluvial workflows ***<u>(Figure 5a;</u> Figure 5b)***, SynGO testing, integrated cross-modality analyses, and targeted neuropeptide/receptor visualization are provided in Supplementary Methods (“Pathway analysis and visualization”).

**Figure 4.**
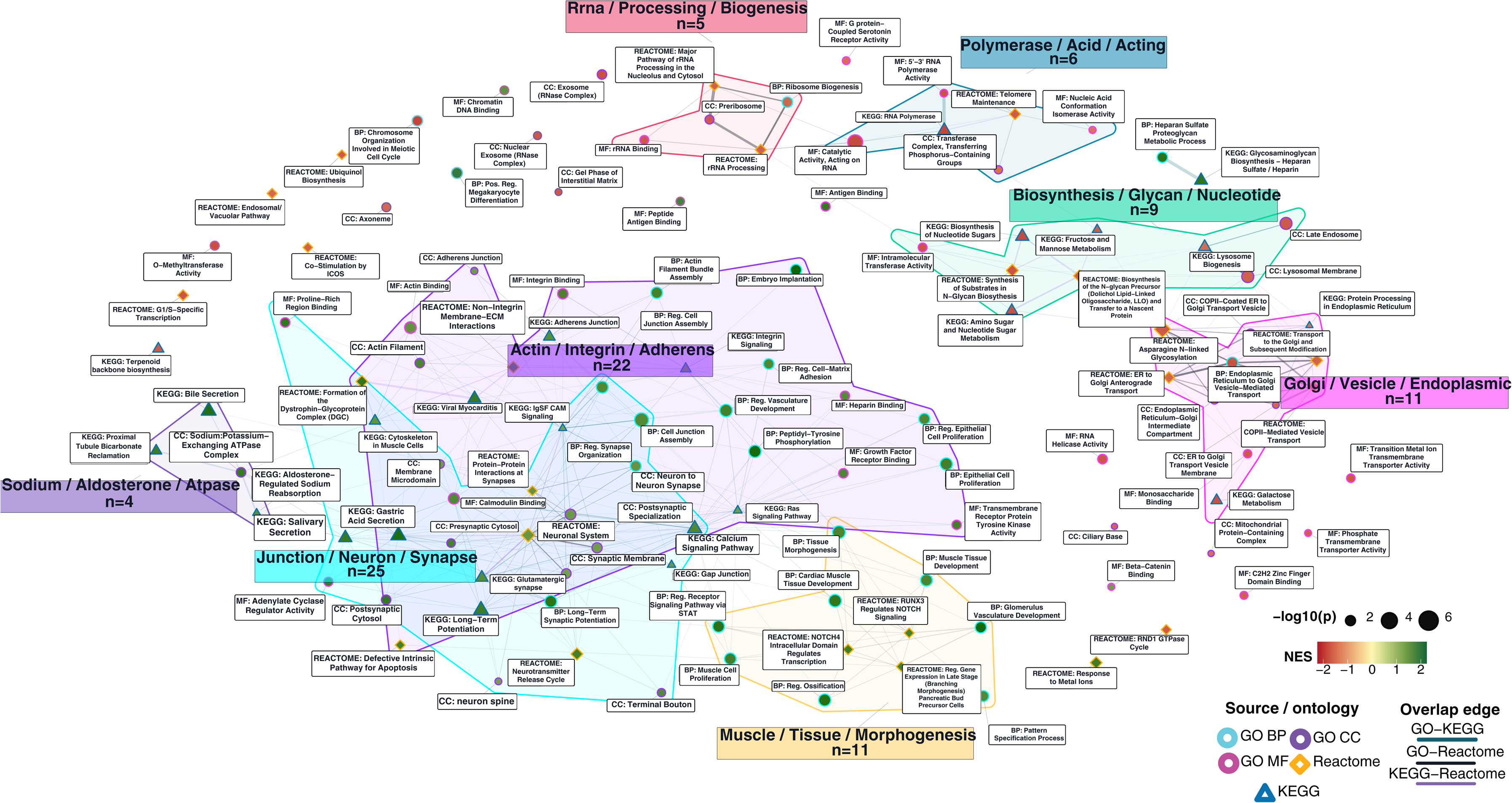
Cross-pathway enrichment map of CRH neuron pathway remodeling in OUD. Network map showing 120 enriched terms from GO BP, GO CC, GO MF, Reactome, and KEGG GSEA comparing CRH neurons from OUD donors and controls. Nodes represent enriched terms or pathways, and edges indicate overlap in leading-edge genes. Node fill indicates normalized enrichment score (red, control-enriched; yellow, near zero; green, OUD-enriched), and node size reflects -log10 nominal P value. Modules highlight recurrent biological themes, including junction/neuron/synapse, actin/integrin/adherens, Golgi/vesicle/endoplasmic, biosynthesis/glycan/nucleotide, rRNA/processing, polymerase-related, and sodium/aldosterone/ATPase processes.

**Figure 5.**
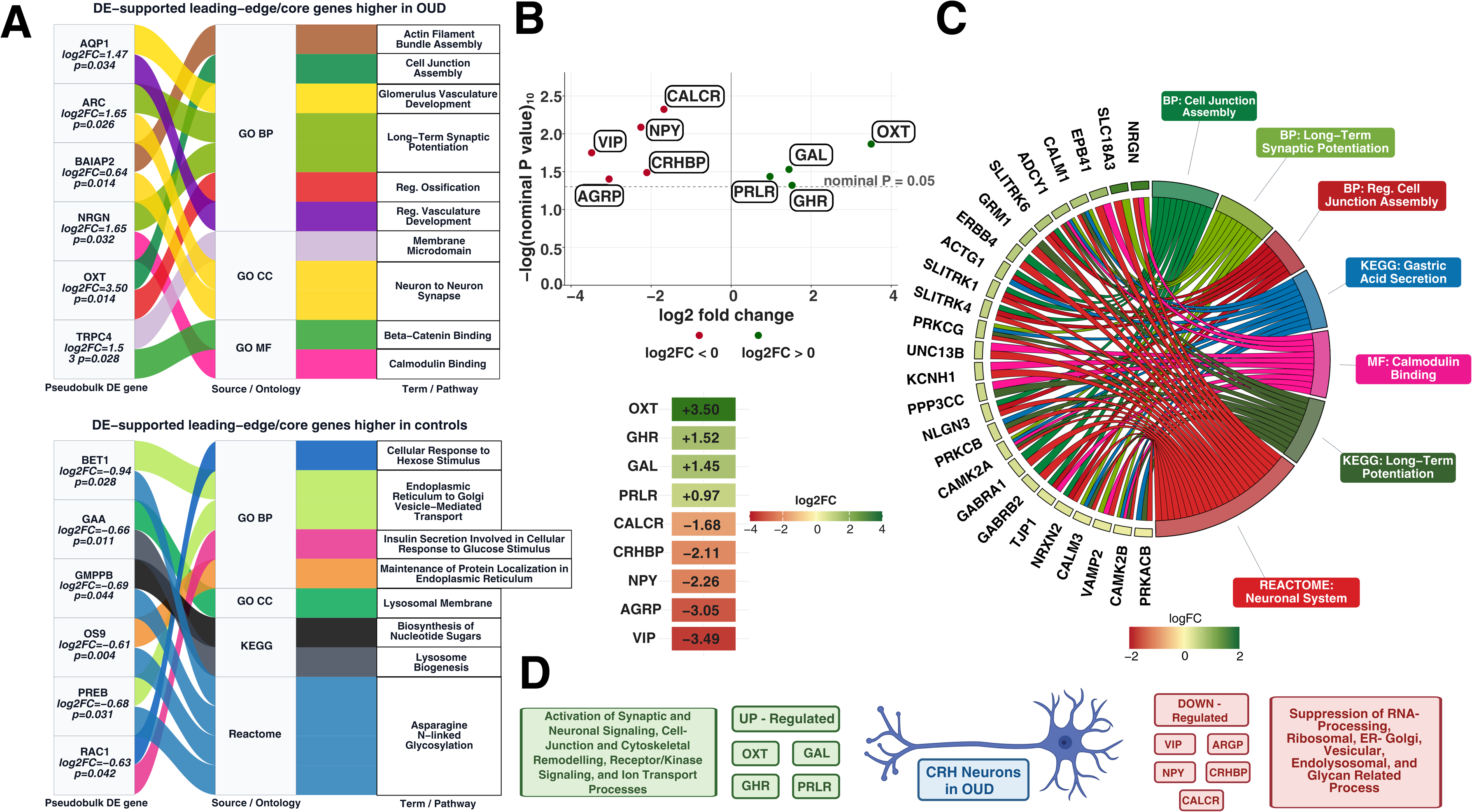
Convergent gene- and pathway-level remodeling of CRH+/PENK+ neurons in opioid use disorder. **(A) DE-supported leading-edge genes higher in OUD and higher in controls.** Alluvial plot linking pseudobulk-significant leading-edge/core genes to enriched GO, Reactome, and KEGG terms in CRH neurons from OUD donors versus controls. Genes were included if they belonged to a plotted leading-edge/core set and had a nominal pseudobulk P < 0.05. Ribbon width reflects –log10(gene P value), and ribbon color identifies the associated term or pathway. **(B) Selected pseudobulk differentially expressed neuropeptide and receptor genes in** CRH+/PENK+ **neurons**. Volcano-style plot showing log2 fold change and nominal P value for nine selected genes. Positive log2 fold change indicates higher expression in OUD, and negative log2 fold change indicates higher expression in controls; the dashed line denotes nominal P = 0.05. Heatmap of log2 fold change for the same genes, ordered from highest to lowest effect size, with color indicating relative expression change. **(C) Junction/Neuron/Secretion pathway module in CRH neurons.** GOChord visualization of the Junction/Neuron/Secretion module, comprising 27 mapped terms/pathways in the enrichment network. Seven representative sectors are shown, spanning GO, KEGG, and Reactome, with 26 genes connected through 73 gene–term/pathway associations. **(D) Summary of pathway-level remodeling in OUD CRH neurons.** Schematic summarizing convergent enrichment findings, with OUD-associated increases in synaptic, junctional, cytoskeletal, receptor-signaling, and ion-transport programs and relative enrichment of RNA-processing, ER-Golgi/vesicular, endolysosomal, and glycan-related processes in controls.

## Results

### Sample characteristics

Donors were balanced across sex and ancestry; both groups were predominantly White males. HC donors were significantly older than OUD donors (HC: 50.8 ± 13.0 years; OUD: 38.8 ± 13.3 years; Wilcoxon rank-sum, W = 325.5, P = 0.007), consistent with OUD mortality more often resulting from accidental overdose at younger ages, whereas control cohorts more often die of age-related causes (Eide et al., 2023). Within this cohort, 16 of 20 OUD donors (80%) died from overdose including an opioid versus none of the 22 HC donors. Post-mortem interval (PMI) was significantly longer in HC donors (HC: 30.5 ± 9.41 h; OUD: 23.0 ± 8.95 h; Wilcoxon W = 324, P = 0.008). Because age and PMI both differed significantly by condition, both were included as covariates in all differential-expression models ***<u>(Supplement Table 1)</u>***.

### Identification of CRH-expressing neuronal populations

Hypothalamus-specific Splatter cells were most abundant across all samples (∼114,170 cells; 33%), followed by oligodendrocytes (18%), astrocytes (11%), OPCs (5%), and microglia (4%) ***(Supplementary Figure 1).*** No cell type differed significantly in proportion between OUD and HC after multiple-testing correction (Speckle/propeller; Phipson et al., 2022; donor as replication unit). Neuronal clusters with >500 nuclei were assessed for expression of 18 hypothalamic neuropeptide and neuronal marker genes, including PVN-associated genes SIM1, CRH, AVP, OXT, SCGN, TRH, SST, VIP, ADARB2, BRS3, and NPY1R ***<u>(Figure 2d-2e) (Supplementary Figure 1c).</u>*** Splatter neurons exhibited the broadest and highest average expression of hypothalamic neuropeptides relative to other neuronal subtypes ***<u>(Supplementary Figure 2).</u>*** Splat_410 showed the highest overall expression of PVN markers and CRH ***<u>(Supplementary Figure 1c)</u>***, and Splat_410_843 and Splat_410_841 showed the greatest CRH expression among Splat_410 subclusters ***<u>(Figure 2e)</u>***. Independent human HYPOMAP and mouse PVN Atlas analyses converged on these CRH-enriched subclusters ***<u>(Figure 2b, Figure 2c)</u>*** (Supplementary Results, “Cross-atlas validation of the CRH-enriched population”).

### Cross-atlas validation of the CRH-enriched population

Human HYPOMAP-guided analysis localized a CRH+/PENK+ parvocellular-like signature to Splat_410_841 and Splat_410_843, with Splat_410_843 showing the strongest CRH enrichment. Independently ***<u>(Figure 2d-2e),</u>*** SIM1-positive re-clustering guided by the mouse PVN atlas ***<u>(Figure 3a, Figure 3c)</u>*** identified a high-confidence Crh-Nr3c1 cluster that mapped predominantly to Splat_410 and specifically to Splat_410_843. These convergent analyses support Splat_410_843 as the principal CRH-enriched PVN subcluster while preserving the broader Splat_410_841 + Splat_410_843 definition for pseudobulk analysis.

Exploratory donor-level composition analysis within the SIM1-positive PVN subset did not identify a significant OUD-control difference in the relative abundance of the CRH cluster ***<u>(Supplementary Figure 14)</u>***. Detailed atlas matching, marker profiles, donor counts, species-specific comparisons, and composition statistics are provided in Supplementary Results (“Splat_410 characterization,” “Mouse PVN Atlas-guided re-clustering,” and “Convergence of parallel pipelines”).

### Differential expression analysis of CRH neurons in OUD versus controls

Differential expression was performed on the Splat_410_841 and Splat_410_843 (CRH+/PENK+) populations using a DESeq2-based donor-level pseudobulk framework (design: Sex + Age + PMI + pH + RIN + Condition). At nominal P < 0.05 with |log FC| > 0.25, 248 genes met the prespecified nominal thresholds: 77 higher and 171 lower in OUD relative to controls. No genes survived transcriptome-wide FDR correction (minimum adjusted P = 0.999). Accordingly, gene-level findings are framed as exploratory and hypothesis-generating, consistent with the sparse representation of the CRH-enriched population.

### Pathway-level enrichment analyses

Because no individual gene survived FDR correction, pathway-level analyses were prioritized. Detailed GO, KEGG, Reactome, and SynGO results, including database-specific enrichment maps, recurrent leading-edge analyses, semantic modules, and GOChord visualizations: provided in Supplementary Results (“Detailed pathway enrichment analyses”). The main-text synthesis below focuses on signals that recur across annotation frameworks. Individual tool analysis support findings of convergent pathway analysis.

### Convergent pathway-level remodeling in OUD CRH neurons

To assess convergence across pathway databases, biological themes were compared across GO BP, CC, MF, Reactome, and KEGG using cross-tool average NES ***<u>(Supplementary Figure 3)</u>***. The most consistent OUD-associated themes involved synaptic/neuronal signaling, adhesion/cytoskeleton/ECM, receptor/kinase signaling, and ion transport/metabolism, whereas the most consistent control-associated themes involved RNA processing/ribosome and vesicle/ER-Golgi/endosome processes.

A cross-pathway enrichment map integrating 120 selected terms (24 per source) resolved OUD-enriched modules centered on junction/neuron/synapse, actin/integrin/adherens, and sodium/aldosterone/ATPase processes, while prominent control-enriched modules centered on Golgi/vesicle/endoplasmic, biosynthesis/glycan/nucleotide, rRNA/processing, and polymerase-related processes ***<u>(Figure 4)</u>***. Among the 24 selected terms per source, q < 0.05 support was present for 20 GO BP, 4 GO CC, 3 GO MF, 11 KEGG, and 1 Reactome term.

Recurrent leading-edge genes were filtered for nominal significance (P < 0.05) in pseudobulk differential expression and recurrence across at least two pathway-source modalities ***<u>(Supplementary Figure 4<u>), (</u>Supplementary Figure 5).</u>*** Genes recurring in OUD-enriched pathways included NRGN, BAIAP2, LRFN5, NECTIN1, ARC, UBE3B, ARHGAP6, GAD2, SYNPO, COL4A2, RGS2, AQP1, and GHR, implicating synaptic plasticity, postsynaptic signaling, dendritic-spine morphogenesis, cell adhesion, inhibitory neurotransmission, and growth-hormone signaling ***<u>(Supplementary Figure 5)</u>***. Control-enriched recurring genes included GMPPB and DYNC1LI2, consistent with glycosylation and microtubule-based intracellular transport ***<u>(Supplementary Figure 5)</u>***. Because no individual genes survived FDR correction, these findings are exploratory and derive support from convergent pathway enrichment rather than gene-level statistical significance.

### Cross-modality pathway visualizations

Direction-specific and Module Specific GOChord and alluvial visualizations ***<u>(Figure 5a), (Figure 5b)</u>*** recapitulated the cross-database contrast between OUD-associated Junction/Neuron/Synapse ***<u>(Figure 5d)</u>*** and cytoskeletal programs and control-associated RNA-processing, ER-Golgi/vesicular, lysosomal, and glycan-related programs. Detailed pathway identities, module panels, and supplemental figure references are provided in Supplementary Results (“Cross-modality visualizations”).

### Differentially expressed neuropeptides and receptors in PVN CRH neurons

Among selected neuropeptide and receptor genes in the CRH+/PENK+ population, pseudobulk analysis identified nominally significant upregulation of OXT (log FC = +3.50, P = 0.014), GHR (log FC = +1.52, P = 0.048), GAL (log FC = +1.45, P = 0.030), and PRLR (log FC = +0.97, P = 0.037). Nominally significant downregulation was observed for VIP (log FC = −3.49, P = 0.018), AGRP (log FC = −3.05, P = 0.040), NPY (log FC = −2.26, P = 0.008), CRHBP (log FC = −2.11, P = 0.033), and CALCR (log FC = −1.68, P = 0.005) ***<u>(Figure 5c)</u>***. These results did not survive FDR correction and should be interpreted as exploratory.

## Discussion

The PVN contains three principal neuronal classes: magnocellular neurons that produce oxytocin and AVP and project to the posterior pituitary; parvocellular neurons that release CRH, thyrotropin-releasing hormone (TRH), and somatostatin (SST) into the hypophyseal portal system; and long-projecting neurons that innervate autonomic centers in the brainstem and spinal cord. PVN CRH responds to changes in peripheral glucocorticoid status via glucocorticoid receptors, establishing these neurons as the uniquely glucocorticoid-responsive population governing HPA axis feedback control (Herman et al., 2016).

Recent single-cell transcriptomic work has shown that hypothalamic CRH expression is not confined to a single neuronal identity but spans GABAergic, glutamatergic, and dopaminergic subclasses, complicating cell-type identification (Li et al., 2026). Because CRH expression is state-dependent and unreliable as a sole marker, constitutive markers are essential: secretagogin, a calcium sensor protein, distinguishes neuroendocrine CRH stress neurons responsible for regulated exocytosis at the median eminence from transient CRH-expressing populations, and the broadly expressed transcription factor SIM1 identifies PVN neurons (Li et al., 2026). This reliance on multi-gene signatures has motivated the development of comprehensive reference atlases. Two are central to the present study. HYPOMAP is an integrated single-cell and spatial atlas of the human hypothalamus that resolves neuronal cell types, including transcriptionally distinct glutamatergic and GABAergic CRH+ populations, and provides human-specific marker definitions for cell-type assignment. In parallel, a recent spatial and projection-based PVN atlas derived from rodent tissue resolved 26 Sim1-positive PVN neuron populations, including discrete CRH subtypes defined by combinatorial markers such as Nr3c1 and Adarb2 (Li et al., 2026). Because these two atlases are built on different species and different methodological frameworks, they offer complementary but non-identical definitions of PVN CRH neurons; a distinction that directly shapes the dual-pipeline cell-identification strategy used here.

In this study, single-nucleus transcriptomic profiling of postmortem human hypothalamus was used to characterize PVN-associated CRH neurons in opioid use disorder (OUD). Rather than defining CRH neurons de novo from unsupervised clustering, identification was anchored to reproducible Allen Brain Map Splatter clusters, which provide stable, shareable identifiers. The HYPOMAP-guided analysis localized a CRH+/PENK+ parvocellular-like population primarily to Splat_410_841 and Splat_410_843, and independent re-clustering of SIM1-expressing nuclei using PVN Atlas–associated markers identified a high-confidence Crh-Nr3c1 population that again mapped preferentially to Splat_410_843. Because these strategies draw on independent references—a mouse PVN taxonomy and a human hypothalamic reference (HYPOMAP; Tadross et al., 2025)—their convergence on the same standardized Allen cluster constitutes cross-atlas, cross-species validation rather than single-pipeline annotation, illustrating how shared cluster nomenclature can make rare hypothalamic populations more tractable and reproducible (Tadross et al., 2025; Herb et al., 2023).

Two principal findings emerge. First, complementary atlas-guided strategies independently converged on Splat_410_843 as the principal CRH-enriched PVN population, providing cross-method support for the population examined downstream. Notably, the human data did not reproduce all features predicted from the mouse taxonomy. ADARB2 was detected in only ∼21% of the human Crh-Nr3c1 cluster, and several transcriptionally distinct human clusters corresponded to single marker-defined mouse identities—consistent with evidence that hypothalamic CRH expression reflects a molecularly diverse, partly state-dependent phenotype spanning glutamatergic and GABAergic lineages rather than a single conserved cell type (Romanov et al., 2017). The CRH+/PENK+ population aligned specifically with the glutamatergic CRH lineage of the human reference. HYPOMAP, generated from more than 400,000 human hypothalamic nuclei, revealed human–mouse disparities even for well-studied neuroendocrine populations (Tadross et al., 2025). Cross-species translation of PVN identities may therefore be more reliable when conserved markers are interpreted within a human transcriptional framework and mapped onto a standardized cluster reference rather than treated as one-to-one equivalents (Tadross et al., 2025; Herb et al., 2023).

These considerations bear on why PVN CRH neurons remain difficult to study in human postmortem tissue. The population is intrinsically rare: CRH-immunoreactive neurons are scattered and confined to specific periventricular parvicellular and posterior subnuclei of the PVN (Raadsheer et al., 1993; Koutcherov et al., 2000), and single-nucleus atlases of the human hypothalamus resolve the CRH/SIM1 cluster as a small fraction of hundreds of molecularly defined cell types (Herb et al., 2023; Tadross et al., 2025). Its position is also anatomically precarious: CRH-immunoreactive perikarya concentrate in the medial PVN, which abuts the wall of the third ventricle, and give rise to dense varicose fibers arching toward the infundibulum and pituitary stalk (Mihály et al., 2002; Koutcherov et al., 2000). Because this tissue lies immediately adjacent to the third ventricle, pituitary stalk, and infundibulum, it is plausibly vulnerable during brain extraction, when the pituitary and stalk are frequently torn away or left in the sella and periventricular tissue is damaged, although this failure mode has not been systematically quantified. Compounding this, the CRH subnuclei are distinguishable only by immuno- or molecular markers rather than gross landmarks, and markers for PVH neuronal populations remain largely undefined (Koutcherov et al., 2000; Li et al., 2026), so targeted microdissection of histologically defined PVN subnuclei is rarely feasible in banked specimens. The present workflow, broad hypothalamic sampling followed by atlas-guided, cross-validated computational identification (Tadross et al., 2025; Li et al., 2026), offers a practical route to recover this population when anatomical landmarks are not preserved.

Second, while no individual gene in the CRH+/PENK+ pseudobulk comparison reached significance after correction for multiple testing, possibly due to the small cell population, multiple functional-enrichment frameworks converged on a directional pattern in which OUD showed greater enrichment of synaptic and neuronal signaling, cell-junction and cytoskeletal remodeling, receptor/kinase signaling, and ion-transport processes, whereas controls showed greater enrichment of RNA-processing, ribosomal, ER-Golgi/vesicular, endolysosomal, and glycan-related functions. Together, these findings indicate that OUD is associated with a shift in the molecular state of human PVN CRH neurons rather than a large change in their relative representation or a few dominant differentially expressed genes.

The most consistent OUD-associated signal involved synaptic organization and structural remodeling and was strongest where it recurred across databases rather than within any single tool, consistent with human postmortem transcriptomic and proteomic evidence of OUD-related synaptic remodeling across brain regions (Seney et al., 2021; Puig et al., 2023). Leading-edge genes that were both differentially expressed and shared across frameworks, including ARC, BAIAP2, LRFN5, NECTIN1, SYNPO, and RGS2, repeatedly populated OUD-enriched synaptic, cell-junction, and actin/cytoskeletal modules in GO, Reactome, and cross-database integration (Levran et al., 2015; Sakloth et al., 2020). Experimental stress models offer precedent to these findings: chronic stress drives synaptic and connectional plasticity of PVN CRH neurons, including enhanced excitatory innervation with increased glutamatergic and noradrenergic terminals (Herman & Tasker, 2016; Flak et al., 2009), reorganizes excitatory transmission with increased functional synapse number (Salter et al., 2018), and up-regulates glutamate-receptor and synapse-related proteins with enhanced NMDA-receptor-dependent drive (Li et al., 2017; Zhou et al., 2018). Because remodeling is not equivalent to a simple increase in excitability but instead entails balanced excitatory and inhibitory synaptogenesis (Miklós & Kovács, 2012; Salter et al., 2018), the present enrichment is best read as altered synaptic architecture and signaling capacity rather than evidence of hyperexcitability. Future work should determine whether this reflects compensatory, maladaptive, or state-dependent remodeling.

A complementary pattern involved relative reduction of intracellular trafficking, biosynthesis, and organelle-function programs in OUD, again most defensible where conserved across GO, Reactome, and KEGG: ER-to-Golgi/COPII transport, endolysosomal function, ribosome biogenesis and RNA processing, and glycosylation/nucleotide-sugar biosynthesis. In a neuroendocrine population dependent on synthesis, processing, trafficking, and regulated peptide release, this coordinated shift raises the possibility that opioid-associated adaptation alters the balance between signaling/plasticity and biosynthetic/trafficking programs. It does not demonstrate impaired CRH processing, dense-core vesicle depletion, or reduced secretion, which require direct measurement.

Beyond biological conclusions, this study develops an analytic strategy for extracting interpretable signals from a rare population in which no individual gene survives stringent correction. Gene-set enrichment detects small but consistent coordinated changes that individual differential-expression testing misses (Subramanian et al., 2005; Qin et al., 2019). Orthogonal frameworks, GO Biological Process, Cellular Component, and Molecular Function, KEGG, and Reactome, were integrated and concordance evaluated via shared leading-edge genes (nominal P < 0.05), then localized into higher-order modules and visualized (alluvial plots, GO chord diagrams, enrichment maps). This provides a template for other rare-cell or low-power datasets (Yoon et al., 2016; Powers et al., 2018). Convergence across these tools does not, however, constitute independent replication, because GO, KEGG, Reactome, and SynGO share overlapping gene memberships, and such overlap can inflate apparent significance rather than reflect corroboration (Simillion et al., 2017; Li et al., 2021); evidence strength also differed, numerous GO and KEGG terms reached corrected significance, only one Reactome pathway reached q < 0.05, and SynGO identified none surviving threshold.

The findings are notable given the state-dependent effects of opioids on the HPA axis. Chronic morphine did not significantly alter hypothalamic CRH under baseline dependent conditions in rats, whereas naloxone-precipitated withdrawal increased hypothalamic CRH and other stress-neuropeptide transcripts (Pintér-Kübler et al., 2013; Nunez et al., 2007). Opioid exposure can therefore produce substantial stress-system adaptation without a persistent large baseline change in CRH transcript, offering one framework for why individual genes did not survive FDR correction while coordinated pathways were altered. The postmortem OUD phenotype likely reflects both chronic adaptation and acute perimortem physiology—particularly relevant here because most OUD donors died from overdose and opioid-specific versus polysubstance involvement was not established for every donor. Outside the hypothalamus, chronic morphine increases excitability in a CeA-to-VTA CRH projection during withdrawal (Jiang et al., 2021). The present results extend opioid-associated CRH plasticity to a transcriptionally defined human hypothalamic population, though PVN and extrahypothalamic CRH populations serve distinct functions and may adapt differently. Pathway findings also partially converge with prior human OUD studies implicating synaptic remodeling, neuroinflammation, and vesicle-trafficking signaling in prefrontal cortex and nucleus accumbens (Seney et al., 2021; Puig et al., 2023). Prior reports of reduced hypothalamic volume in male heroin users (Müller et al., 2018) operate at a different scale than relative cell-type proportions and together motivate a combined spatial, morphometric, and cell-type-resolved study.

Several nominally significant neuropeptide/receptor genes generate specific, exploratory hypotheses. In the CRH+/PENK+ population, pseudobulk analysis identified nominal upregulation of OXT, GHR, GAL, and PRLR, and downregulation of VIP, AGRP, NPY, CRHBP, and CALCR. Higher OXT and GAL with lower CRHBP could indicate coordinated modulation of stress-peptide signaling and CRH bioavailability, since CRH-binding protein binds CRH with sub-nanomolar affinity and limits receptor activation (Ketchesin et al., 2017; Slater et al., 2016); PVN OXT is co-distributed with and can inhibit CRH neurons and is implicated in drug-seeking (Huang et al., 2024; King, Gano & Becker, 2020), and galanin is co-expressed with enkephalin in parvocellular PVN neurons (Barson et al., 2011; Genders, Scheller & Djouma, 2020). Reduced VIP, AGRP, and NPY suggest OUD adaptation may extend to metabolic and circadian integration (Cowley et al., 1999; Blasiak et al., 2017; Engström Ruud et al., 2020; Mihály et al., 2002).

The study has several limitations. Postmortem tissue studies cannot distinguish pre-existing vulnerability from chronic-exposure or time of death effects; overdose, polysubstance exposure, agonal physiology, hypoxia, medication, and timing of last opioid exposure may all influence transcription (Ferreira et al., 2018; Dachet et al., 2021; Dai et al., 2020). The CRH+/PENK+ population is rare; requiring ≥10 nuclei per donor reduced the comparison to 25 donors, limiting power and precluding robust sex- or covariate-specific analysis. Sampling used a hypothalamic punch rather than prospective PVN microdissection. Future work should combine transcriptomic identification with spatial localization of Splat_410_843-like neurons and targeted protein/peptide measurements.

In conclusion, atlas-guided cell-specific profiling identifies a human PVN CRH-enriched population exhibiting pathway-level molecular remodeling in OUD. Convergence of two independent atlas-based strategies on a single standardized Allen Brain Map cluster demonstrates a reproducible approach for isolating rare, anatomically fragile hypothalamic populations from imperfect postmortem tissue. Across enrichment approaches, OUD was most consistently associated with greater synaptic, junctional, cytoskeletal, receptor-signaling, and ion-transport programs and relative reduction of RNA-processing, ER-Golgi/vesicular, endolysosomal, and glycan-related processes ***<u>(Figure 5e).</u>*** These results are consistent with an altered adaptive state of human CRH neurons involving synaptic and cellular remodeling.

