## Supplementary Material for "Atlas-Guided Cell-Specific Transcriptomics Identifies Pathway-Level Remodeling of Human Hypothalamic CRH Neurons in Opioid Use Disorder"

**Supplementary Information**

**Supplementary Methods**

**Library preparation and sequencing**

Nuclei were isolated using the 10x Genomics Chromium Fixed RNA Profiling tissue fixation and dissociation protocol (CG000553, "Demonstrated Protocol: Tissue Fixation & Dissociation," Revision B), performed as described by the manufacturer. Nuclei were counted using a Countess 3 FL automated cell counter with both acridine orange/propidium iodide (AO/PI) and Trypan blue staining; counts for each stain were taken in duplicate and averaged prior to loading. Samples were multiplexed 16-plex using probe barcodes and processed with the Chromium Fixed RNA Profiling Reagent Kit for Multiplexed Samples (Flex v1 chemistry; CG000527). Sequencing was performed on a NovaSeq X Plus using a 10B flowcell. The 42-donor cohort was distributed across three sequencing pools: Batch1_S1_S16, Batch2_S17_S32, and Batch4_S49_S64 .

**Read alignment and quantification**

Libraries were processed with cellranger multi (Cell Ranger 7.1.0) against the GRCh38-2020-A reference using the Chromium Human Transcriptome Probe Set v1.0.1 (targeting 18,082 genes). Mean reads per cell at the pool level ranged from 8,062 (Batch1) to 11,046 (Batch4). Two pools showed donor-level unique molecular identifier (UMI) imbalance: in Batch1_S1_S16, donor 926 contributed 56.12% of total UMIs; in Batch4_S49_S64, three donors (217, 230, 234) together contributed 59.55% of UMIs. These same three donors dominated clusters 39, 40, and 41 in downstream analysis, though a causal relationship between pool loading and cluster composition was not tested. None of these donors, nor clusters 39, 40, or 41, were excluded from any downstream analysis. These clusters do not meaningfully contribute to this study's central findings: Splat_410, the population of primary interest, draws only 14 and 4 cells from clusters 41 and 42, respectively, out of 16,596 total cells. Cells from clusters 39, 40, and 41 contribute zero cells to either of the two most significant general cell-type differential expression categories (Amygdala excitatory, LAMP5-LHX6 and Chandelier) and only 3 of 12,531 tested cells to the third (Vascular).

**Quality control and filtering**

*Nuclei counts.* A total of 408,684 nuclei were analyzed after donor exclusion and before mitochondrial filtering; 346,435 remained after mitochondrial filtering; and 343,819 were retained after targeted, cell-level doublet removal ***(Supplementary Figure 6)***.

*Per-nucleus metrics.* Donor-level median nCount_RNA (post-mitochondrial filtering) was 1,046 (HC) versus 1,106 (OUD), a non-significant ~5.7% difference (Wilcoxon W = 197, P = 0.575); donor-level median nFeature_RNA (post-mitochondrial filtering) was likewise non-significant. (Wilcoxon W = 198, P = 0.587). Dataset-wide summaries were: pre-mitochondrial filtering, nCount_RNA median 1,237 (mean 2,941), nFeature_RNA median 952 (mean 1,539); post-mitochondrial filtering, nCount_RNA median 1,152 (mean 2,734), nFeature_RNA median 902 (mean 1,470) ***(Supplementary Table 2)***. The median-versus-mean gap reflects heavy right skew from a small number of extreme-count nuclei (maximum nCount_RNA = 425,714 in a single nucleus, classified as a doublet by scDblFinder but retained in the final dataset, as its cluster (14) fell outside the scope of the targeted doublet pruning described below.

*Mitochondrial threshold.* A minimum of 500 nuclei per donor was required; all 42 donors met this criterion. Nuclei exceeding 20% mitochondrial content had been excluded during initial object assembly by the sequencing core, so the distribution reported here is conditional on that prior filter. Among retained nuclei, those with ≥5% mitochondrial gene content were excluded (percent.mt: median 1.063%, mean 2.311%, IQR 0.410–2.970%); 84.77% fell below the 5% threshold and were retained ***(Supplementary Figure 6)***.

*Doublet identification and removal.* Doublets were detected using scDblFinder on the fully clustered dataset (overall doublet rate 10.62%). Rather than applying a universal doublet-rate filter, a cluster-specific diagnostic approach distinguished true rare populations from technical artifacts: flagged clusters were re-embedded, re-clustered in isolation, and examined for marker identity and donor composition. Three clusters showed high doublet fractions (44–56% versus the 10.62% dataset average; ***(Supplementary Figure 7)***) and were predominantly doublets and stress artifacts; scDblFinder-flagged doublets within these three clusters (2,616 nuclei) were removed at the cell level. Singlets within these clusters, and all nuclei in every other cluster, were retained regardless of per-cell doublet status. Five additional small clusters (38-42) flagged for donor dominance rather than doublet fraction were examined identically ***(Supplementary Figure 8)***. None of these clusters were found to be predominantly doublet driven.

*Final nuclei count.* 343,819 nuclei were retained (346,435 passing QC and mitochondrial filtering, minus 2,616 removed by targeted cell-level doublet pruning).

*High-count-cell follow-up.* A total of 5,331 nuclei (1.54% of 346,435) exceeded 20,000 UMIs; of these, 3,840 were classified as singlets by scDblFinder (1.12% of the 343,819-nucleus post-pruning dataset), consistent with the known reduced sensitivity of doublet detection for homotypic doublets. Rates of high-count singlets by cell type were highest in choroid plexus (9.05%, more than double the next-highest category; all others 1–3.6%). Choroid plexus epithelium is well documented as highly secretory (Damkier, Brown & Praetorius, 2013, *Physiol Rev*; Thouvenot et al., 2006, *Proteomics*, PMID 17051638), though this literature establishes secretory and proteomic capacity rather than elevated per-nucleus transcript counts in snRNA-seq specifically; this is therefore a plausible but unverified explanation. Choroid plexus was in any case already excluded from between-group comparisons on donor-concentration grounds. High-count singlets within splat_410 (1.09%) sat at the dataset background rate, and the CRH+/PENK+ subset (0.42%) below it ***(Supplementary Figure 9)***. No upper-bound count filter was applied.

Sex-gene concordance. Donor sex records were cross-referenced against Y-chromosome gene expression (UTY, DDX3Y, KDM5D). Sex identification was fully concordant across all 42 donors (29 male, 13 female), with no mismatches or borderline cases: donors recorded as female showed 0.0–0.5% of nuclei expressing each Y-chromosome gene, consistent with ambient background, whereas donors recorded as male showed 4.0–51.8%. The lowest single male value (4.0%, donor 67940) exceeded the highest female value (0.5%, donor 67992) by eightfold ***(Supplementary Figure 10)***.

**Batch integration**

Harmony (v2.0.5) was used for batch integration (`group.by.vars = orig.ident`; convergence at 6 iterations within a 100-iteration ceiling). Batch mixing was evaluated with the Local Inverse Simpson's Index (LISI; Korsunsky et al., 2019). Donor mixing (iLISI) improved ~89% after integration (baseline PCA median 3.249, IQR 2.015–4.828; post-Harmony median 6.136, IQR 4.494–7.696; ***(Supplementary Figure 11)***). iLISI was interpreted relative to this pre/post baseline rather than the theoretical maximum of 42, because rare, unevenly distributed populations naturally show lower local donor diversity without indicating a batch effect (Andreatta et al., 2024). Cluster purity (cLISI) was assessed with a stratified five-replicate design (retaining all cells from small/rare clusters, down sampling larger clusters), confirming high reproducibility (SD < 0.013 per cluster; dataset median 1.026, IQR 1.000–1.360) and correcting an initial single-sample analysis that overestimated cLISI for small clusters due to a down sampling artifact ***(Supplementary Figure 11)***.

**Dimensionality reduction and clustering**

PCA used the first 50 principal components (`npcs = 50`). Harmony integration and downstream UMAP and neighbor-finding used the first 30 Harmony-corrected dimensions (`dims = 1:30`). `FindClusters` (Louvain, resolution = 0.8) initially produced 61 communities; 18 singletons were collapsed, yielding 43 final clusters (modularity 0.9212).

**Neuronal and non-neuronal cell populations**

Neuronal nuclei were subset and visualized by UMAP, revealing 21 neuronal superclusters: upper-layer intratelencephalic, upper rhombic lip, thalamic excitatory, Splatter, miscellaneous, midbrain-derived inhibitory, mammillary body, lower rhombic lip, LAMP5-LHX6 and Chandelier, hippocampal dentate gyrus, hippocampal CA4, hippocampal CA1–3, medium spiny neuron, eccentric medium spiny neuron, deep-layer near-projecting, deep-layer intratelencephalic, deep-layer corticothalamic and 6b, CGE interneuron, cerebellar inhibitory, and amygdala excitatory ***(Figure 2a)*.** Non-neuronal nuclei comprised vascular cells, oligodendrocyte precursors, oligodendrocytes, microglia, fibroblasts, ependymal cells, committed oligodendrocyte precursors, choroid plexus epithelium, Bergmann glia, and astrocytes ***(Figure 2a)*.**

*Gene Marker Panel. Canonical panel:* DotPlot across 43 Louvain clusters confirmed broad cell identities ***(Supplementary Figure 12)***: oligodendrocytes (MBP, PLP1, MOBP, MOG), OPCs (PDGFRA, CSPG4, OLIG1, OLIG2), astrocytes (AQP4, GFAP, SLC1A2), microglia (CSF1R, P2RY12, CX3CR1, C1QA), vascular cells (CLDN5, FLT1, PDGFRB), ependymal cells (FOXJ1, CCDC153, PIFO), choroid plexus (TTR, FOLR1, PRLR), fibroblasts (DCN, COL1A1), GABAergic neurons (GAD1, GAD2, SLC32A1), and thalamic-excitatory neurons (SLC17A6, SLC17A7, GRIN2B).

*Gene Marker Panel. Neuropeptide panel:* (KISS1, GNRH1, OXT, AGRP, POMC, GHRH, PDYN, TH, AVP, SST, PENK; ***(Supplementary Figure 13)***) combined with an MMC atlas top-enriched-gene reference panel (AVP, OXT, OTP, SIM1, TH, TRH, CRH, PDYN, SCGN, CARTPT, GHSR; Siletti et al., 2023; ***(Supplementary Figure 13)***) identified hypothalamic neuropeptide-expressing neurons within Splat_410. AVP was expressed in 57.5–88.2% of cells across all 12 Splat_410 subclusters, with 12- to 75-fold enrichment relative to other dataset cells (AVP ~75×, OXT ~53×, TH ~42×, CARTPT ~26×, CRH ~20×).

**CRH neuron identification**

**Human HYPOMAP-guided Splat_410 subcluster-membership approach (Pipeline 1)**

Pipeline 1 Splat_410 was identified within the MMC-derived Splatter supercluster (16,596 cells; 12 subclusters; Siletti et al. taxonomy aliases 724, 725, 833, 834, 837–844), all predominantly hypothalamic in the reference atlas (66–98% hypothalamus per subcluster). Specificity analyses showed 12- to 75-fold enrichment for hypothalamic neurosecretory markers (AVP ~75×, OXT ~53×, TH ~42×, CARTPT ~26×, CRH ~20×, SCGN ~19×, SIM1 ~16×, OTP ~14×, TRH ~12×), with the master developmental transcription factors SIM1 and OTP among the most enriched. The 12 subclusters split into an AVP/OXT-dominant magnocellular-type majority and a smaller, transcriptionally distinct CRH+/PENK+ parvocellular-type signature concentrated in Splat_410_841 (~1,254 cells) and Splat_410_843 (~3,520 cells), in which CRH was ~13-fold higher in Splat_410_843 than other subclusters and PENK was expressed in ~46–51% of cells (versus <10–20% elsewhere). Cross-validation against HYPOMAP confirmed this identity: marker genes (CRH, CARTPT, AVP, PGF, TH) of HYPOMAP cluster C4-297 ("Mid-3 GLU-3 AVP PGF") corresponded to Splat_410_841 and Splat_410_843, aligning with the glutamatergic CRH⁺ lineage by manually referencing the Tadross supplementary data cluster_tree_annotations tab. PENK was not among C4-297 top markers, and a separate PENK-specific HYPOMAP cluster (C4-354) showed no overlap with AVP or CRH, indicating PENK is a secondary co-expressed feature rather than an independent population ***(Figure 2c)***. To avoid inadvertent inclusion of CRH⁺ neurons outside this group, the CRH+/PENK+ population was defined by subcluster membership (Splat_410_841 + Splat_410_843) rather than expression thresholds. Subcluster-level MMC mapping confidence was below the dataset average (median bootstrapping probability ~0.48 vs. ~0.62), but was retained without filtering given strong marker and cross-atlas support.

**Mouse PVN Atlas-guided SIM1-positive re-clustering (Pipeline 2)**

Nuclei with detectable *SIM1* expression (normalized expression >0) were subset (2,847 nuclei from 25 donors) ***(Supplementary Figure 14)*** and independently re-clustered on Harmony-corrected embeddings ***(Supplementary Figure 15)*** (30 Harmony dimensions, k = 30 nearest neighbors, SNN resolution = 3.5), yielding 22 Harmony clusters ***(Supplementary Figure 16)***. Post hoc biological labels were assigned using a marker-based strategy informed by the mouse PVN Atlas ***(Figure 3a)*** (Li et al., 2026). For each cluster, normalized RNA expression of candidate marker genes was summarized by two metrics: average normalized expression and the percentage of cells with detectable expression. Relative marker enrichment across Harmony clusters was quantified using a combined marker score, defined as the gene-wise z-scored average expression plus 0.75 times the gene-wise z-scored detection frequency (marker score = z-scored average expression + 0.75 * z-scored detection frequency) ***(Supplementary Figure 17)***.

Biological labels were assigned using rule-based marker combinations, each consisting of a primary marker paired with a label-refining support marker. Primary markers comprised neuropeptide and neurotransmitter associated genes, including CRH, AVP, OXT, TRH, SST, GHRH, SLC17A6, ASB4, and BRS3. Support markers included NR3C1, ADARB2, TH, TAC1, UCN3, GHRH, CALB2, SFRP2, FOXP1, OMP, and NFIB ***(Figure 3c)***.

Confidence tiers reflected the strength of marker support stratified by high, medium and low confidence. High confidence: strong primary-marker enrichment together with a biologically relevant support marker in a sufficiently sized cluster. Medium confidence: interpretable marker support with weaker, sparser, or less complete support-marker evidence. Low confidence: partial resemblance to a PVN Atlas-associated marker pattern but insufficient marker support for robust downstream label-specific interpretation ***(Supplementary Figure 17)***. High- and medium-confidence clusters were retained for downstream biologically labelled analyses. Low-confidence clusters were retained within the unsupervised Harmony clustering results but excluded from downstream label-specific analyses.

This pipeline identified a high-confidence Crh-Nr3c1 cluster ***(Supplementary Figure 1a)*** (Harmony cluster 1; 199 nuclei; CRH detected in 57.8% of nuclei, NR3C1 in 56.3%, and ADARB2 in 20.6%) ***(Figure 3b; Supplementary Figure 1b)***.

*Convergence between pipelines.* Post hoc cross-referencing of Harmony cluster 1 with MMC annotations showed 162 of 199 nuclei (81.4%) mapped to Splat_410 at the broad cluster level ***(Supplementary Figure 18)***, and 110 of 199 (55.3%) to Splat_410_843 at the subcluster level ***(Supplementary Figure 19)***, the same subcluster identified as CRH-enriched by Pipeline 1, independently localizing the CRH-enriched PVN population to Splat_410_843 per MMC.

*Species-specific differences in PVN cell-type resolution.* Several differences between mouse-atlas predictions and the human data emerged: 1) the mouse atlas defines CRH neurons as a single Crh-Nr3c1 population with an ADARB2-co-expressing subset, but ADARB2 was detectable in only 20.6% of human Crh-Nr3c1 nuclei ***(Supplementary Figure 17)***, suggesting the Crh-Adarb2 subtype is less prominent or distinct in human PVN; 2) the mouse atlas separates CRH, AVP, OXT, TRH, and SST into discrete clusters, whereas multiple human Harmony clusters shared one marker-defined identity (e.g., Avp-Th: clusters 2, 4, 11, 16), indicating greater human transcriptional heterogeneity ***(Figure 3)***; 3) the glutamatergic/GABAergic CRH⁺ distinction emphasized by human HYPOMAP is not a primary organizing feature of the mouse atlas, and the CRH⁺/PENK⁺ subclusters aligned specifically with the glutamatergic lineage; 4) CRH-PENK co-expression (~46–51%) was not recapitulated as an independent cell type in either atlas, suggesting a species-specific or context-dependent feature.

**CRH+/PENK+** **pseudobulk analysis**

CRH+/PENK+ subcluster (Splat_410_843 and Splat_410_841). Two considerations were addressed before finalizing. First, the choice between pseudobulk DESeq2 and a mixed-effects model was resolved by literature review: the donor (not the nucleus) is the correct unit of biological replication even at smaller sample sizes, and properly implemented pseudobulk methods are statistically comparable to mixed-effects models (Lee & Han, 2024); a revised benchmark found pseudobulk superior in both type-I and type-II error after correcting methodological issues in an earlier benchmark that favored mixed models (Murphy & Skene, 2022). Although DESeq2 has underperformed some pseudobulk tools in certain benchmarks (Squair et al., 2021), it was retained because it accommodates donor-level covariates directly rather than regressing them out. Second, 17 of 42 donors had fewer than 10 cells in this subpopulation (some 1–2), yielding unstable pseudobulk profiles; applying a ≥10-cell-per-donor minimum (no directional bias) retained 25 donors (14 HC / 11 OUD) with balanced subcluster representation (Splat_410_841: 494 HC / 756 OUD; Splat_410_843: 1,667 HC / 1,849 OUD). No genes reached FDR significance among 10,214 tested ***(Supplementary Figure 33).***

**Pathway analysis and visualization**

Functional enrichment was characterized across GO Biological Process, Cellular Component, and Molecular Function; KEGG; Reactome; and SynGO (58). GO, KEGG, and Reactome used ranked GSEA on the full pseudobulk gene ranking, whereas SynGO used over-representation analysis of thresholded foreground genes. Analyses were first performed independently and then integrated across sources using NES, leading-edge gene overlap, and GO semantic similarity.

Enrichment was performed at the pseudobulk level; genes were ranked or thresholded from a single per-sample pseudobulk differential expression analysis rather than by per-cell gene set scoring. Analyses were performed in R 4.6.1 using clusterProfiler 4.20.0, org.Hs.eg.db 3.23.1, AnnotationDbi 1.74.0, GOSemSim 2.38.3, enrichplot 1.32.0, GOplot 1.0.2, and igraph 2.3.3, together with the plotting packages listed below. The org.Hs.eg.db annotation package used Entrez Gene source date 2026-Mar18, GO source date 2026-01-23 from go-basic.obo, and GO-to-Entrez source date 2026-Mar18. Annotation database releases were as follows: GO/org.Hs.eg.db 3.23.1, built locally under R 4.6.1 on 2026-07-22 UTC; KEGG release 119.0 (July 1, 2026), accessed through clusterProfiler/KEGGREST when the KEGG GSEA outputs were generated on 2026-08-05; ReactomePA 1.56.0 using Reactome version 96, source date 2026-04-06, with the local Reactome GSEA outputs generated on 2026-08-05; and SynGO release 1.3 (portal release 20250226; SynGO_bulk_20250218.json and genes_lookup.json cached locally on 2026-08-03).

**GO Enrichment (BP, CC, MF)**

GO enrichment derived from clusterProfiler gseGO results (clusterProfiler 4.20.0; minGSSize = 10 and maxGSSize = 500) for the CRH+/PENK+ neuronal pseudobulked DEGs, analyzed separately for BP, CC, and MF. Wald statistic GSEA ranking metric from the pseudobulk DESeq2 differential expression workflow, and genes were ranked across the full gene list. Simplified GO result tables spanned observed setSize values of 1 to 360 after result generation and filtering. GSEA used weighted enrichment with exponent p = 1. ClusterProfiler defaults were method = "multilevel", nPerm = 1000, adaptive = FALSE, minPerm = 101, maxPerm = 1e5, and pvalThreshold = 0.1 ***(16)***.

Result tables were imported, cleaned, and combined across ontologies (GO BP, CC, MF). Leading-edge genes were extracted from the core_enrichment field, mapped from Entrez IDs to gene symbols using org.Hs.eg.db, and joined to the pseudobulk DESeq2 results where available; gene-level log2FC and nominal p values were used for gene coloring and cross-validation in leading-edge figures. Terms were ranked within ontology by q value, then nominal p value, then absolute NES. Unless otherwise stated, q value was used as the FDR metric for ranking and display, with q <= 0.25 as the primary FDR threshold and nominal p < 0.05 as the nominal-evidence threshold.

For recurrent leading-edge heatmaps, top terms were selected within each ontology and leading-edge genes were ranked by recurrence across terms, recurrence across ontologies, direction span, pseudobulk nominal p value, and absolute log2FC ***(Supplementary Figure 20)***. The expanded figure used the top 20 terms per ontology and up to 54 recurrent genes; a DE-validated version retained only recurrent genes present in the pseudobulk DE table with nominal gene-level p < 0.05 and omitted terms without retained genes ***(Supplementary Figure 21), (Supplementary Figure 22)***.

Enrichment maps were generated in an enrichplot-inspired format. The primary map connected the top 20 terms per ontology when they shared ≥ 2 leading-edge genes or had Jaccard overlap ≥ 0.04 (up to 300 edges). Louvain community detection was applied to the Jaccard-weighted term-overlap graph, and layout used Fruchterman–Reingold (4,000 iterations, seed 149). Node fill encoded NES, node size encoded −log10(q value), and node border color indicated ontology. For the redundancy-reduced map, candidate terms were selected at q ≤ 0.25 with a minimum of 24 terms per ontology, redundant terms were collapsed within ontology using Wang similarity (cutoff 0.70), and representative terms were chosen by q value, nominal p value, and absolute NES. The final reduced map displayed 72 terms (24 per ontology) with edges retained at minshared = 1, minjaccard = 0.025, and max_edges = 240; ontology border colors were BP #00FFFF, CC #9933FF, and MF #FF33FF, and module boundaries were drawn as transparent hulls around Louvain communities ***(Supplementary Figure 23)***.

GOChord plots were generated with the GOplot workflow (`circledat`, `chorddat`, `GOChord`). GO-term sectors represented selected enriched terms, ribbons represented leading-edge gene membership, and gene sectors were colored by CRH pseudobulk log2FC; terms were assigned unique colors by GO ID to make term-specific overlaps traceable. The recurrent GOChord used the top 18 candidate terms per ontology, selected recurrent genes appearing in ≥ 2 selected terms where possible (up to 28 genes), and displayed terms supported by ≥ 2 selected genes. Direction-specific panels were generated separately for positive-NES (higher in OUD) and negative-NES (higher in controls) terms using q ≤ 0.25 or nominal p < 0.05, with up to 3 terms per ontology and up to 24 genes per panel ***(Supplementary Figure 24)***. Semantic-module panels were derived from enrichment-map modules containing ≥ 4 terms, ranked by term count, minimum q value, and absolute mean NES, and displayed up to 7 terms and 24 genes per module ***(Supplementary Figure 25)***. GOChord rendering parameters were space = 0.02, nlfc = 1, lfc.min = −2, lfc.max = 2, with genes ordered by log2FC and ribbon colors assigned uniquely by term; GO labels were placed outside the chord circle with repelled labels. Across all figures, a shared palette was used in which red denotes higher enrichment/expression in controls, yellow denotes a near-zero effect, and green denotes higher enrichment/expression in OUD.

**KEGG Pathway Analysis**

KEGG enrichment was evaluated from preranked GSEA output for the OUD-versus-control contrast, using the Wald-statistic ranking over the full tested gene universe (KEGG release 119.0, accessed via clusterProfiler/KEGGREST on 2026-08-05; GSEA parameters as specified in the GO subsection unless otherwise noted). Pathways were retained for visualization if non-empty and nominally significant (p < 0.05) and were ranked by q value, nominal p value, and absolute NES. Leading-edge genes were extracted from the `core_enrichment` field, mapped from Entrez IDs to symbols via org.Hs.eg.db, and joined to the CRH pseudobulk DE table by symbol; gene-level validation retained only genes present in the pseudobulk table with nominal gene-level p < 0.05.

A KEGG enrichment map represented each retained pathway as a node, with pairwise similarity from shared leading-edge genes (shared count and Jaccard overlap). Edges were drawn for pathway pairs sharing ≥ 1 gene or with Jaccard ≥ 0.025, limited to the top 220 edges (ranked by shared count and Jaccard) and weighted by Jaccard overlap. Modules were identified by Louvain community detection on the Jaccard-weighted graph, and the network was laid out with Fruchterman–Reingold (seed 149, 4,000 iterations). Node fill encoded NES, node size encoded −log10(nominal p value), and boundaries were drawn around Louvain communities. Because KEGG lacks a GO-style ontology graph for Wang similarity, modules were defined by gene overlap rather than semantic similarity; module labels were generated post hoc from recurrent informative keywords in member-pathway descriptions after removal of common pathway stop words, and serve as descriptive summaries rather than formal semantic classes ***(Supplementary Figure 26)***.

Additional KEGG visualizations included direction-specific NES heatmaps ***(Supplementary Figure 27)***, recurring ***(Supplementary Figure 28)*** and DE-validated ***(Supplementary Figure 29)*** leading-edge gene heatmaps, effect–evidence plots ***(Supplementary Figure 30)***, and GOChord diagrams ***(Supplementary Figure 31), (Supplementary Figure 32)***. Direction-specific heatmaps grouped retained pathways by NES sign; recurring-gene heatmaps displayed selected leading-edge genes across pathways with tile color encoding pathway NES; GOChord diagrams connected selected pathways to their leading-edge genes, with unique pathway-sector colors and gene sectors ordered and colored by CRH pseudobulk log2FC.

**Reactome Pathway Analysis**

Reactome GSEA results for the CRH-neuron OUD-versus-control comparison were generated with ReactomePA 1.56.0 (Reactome version 96, source date 2026-04-06; outputs generated 2026-08-05) and imported, and pathways with non-empty identifiers and descriptions were retained and ranked by q value, nominal p value, and absolute NES. For visualization, nominally enriched pathways (p < 0.05) were selected, with up to 72 pathways used for the enrichment map. Direction was assigned by NES sign (positive = higher in OUD; negative = higher in controls). Leading-edge genes were parsed from `core_enrichment`, mapped from Entrez IDs to HGNC symbols via AnnotationDbi and org.Hs.eg.db, and joined to the pseudobulk DE table for gene-level validation and GOChord log2FC coloring; DE-validated heatmaps retained only genes with nominal pseudobulk p < 0.05.

The enrichment map represented pathways as nodes and shared leading-edge genes as edges, with overlap quantified by shared gene count and Jaccard index (shared core genes ÷ union of the two pathway core sets). Edges were retained for shared genes ≥ 1 or Jaccard ≥ 0.025 (maximum 280 edges; observed network, 168 edges), with edge width proportional to Jaccard overlap and edge transparency reflecting shared-gene count. Layout used Fruchterman–Reingold (seed 149, niter = 4,000). Louvain modules were detected with `igraph::cluster_louvain` weighted by Jaccard overlap and were therefore defined by core-gene overlap rather than by pathway names. Module labels were assigned post hoc by a keyword procedure: member descriptions were lowercased, non-alphanumeric characters replaced with spaces, tokens shorter than four characters removed, and a Reactome-specific stop-word list excluded (generic terms such as reactome, pathway, process, regulation, response, protein, transport, activity, activation, from, alpha, linked, intracellular, initiated, initiation, processing, regulate/regulated/regulates, specific, system, and related non-biological connectors). The three most frequent remaining terms per module were title-cased and slash-joined to form the label; these are descriptive summaries of Louvain overlap neighborhoods and should not be interpreted as formal Reactome ontology classes ***(Supplementary Figure 34)***.

Direction-specific heatmaps displayed the top 56 nominal pathways (ranked by q value, nominal p value, absolute NES) ***(Supplementary Figure 35)***. Recurring leading-edge gene heatmaps displayed the top 36 pathways and up to 56 genes ranked by recurrence across pathways, maximum absolute NES, best pathway p value, and gene symbol; the DE-validated version restricted genes to pseudobulk DE p < 0.05 ***(Supplementary Figure 36), (Supplementary Figure 37)***. GOChord diagrams (`GOplot::circledat`, `chorddat`, `GOChord`) included direction-specific panels selecting up to seven OUD-enriched and seven control-enriched pathways (ranked by q value, nominal p value, absolute NES) with sufficient leading-edge gene representation; gene sectors were ordered by logFC and colored by pseudobulk log2FC (red/yellow/green), and pathway sectors were uniquely colored by pathway ID ***(Supplementary Figure 38)***. Module-driven panels used Louvain modules from the enrichment map (up to seven pathways per module, up to 28 genes per panel) ***(Supplementary Figure 39)***. Biological-theme heatmaps used a separate keyword-based description classifier—independent of Louvain clustering—to group pathways into high-level themes, with tile fill encoding mean NES per theme-by-direction group and text labels reporting pathway counts ***(Supplementary Figure 40)***. Effect evidence plot was also generated ***(Supplementary Figure 41).***

**SynGO Synaptic Ontology Analysis**

Synaptic annotation enrichment (SynGO release 1.3) was assessed by ORA across the synaptic CC and BP domains, using DEGs from the CRH-PENK+ pseudobulk analysis. UP and DOWN foreground sets thresholds: nominal p < 0.05 and |log2FC| > 0.25. Background included all genes testable in the pseudobulk differential expression analysis, augmented with foreground genes to match the SynGO webtool foreground/background structure; the whole genome was not used as background.

SynGO CC and BP ontology structures derived from SynGO bulk annotation file, and annotations were propagated recursively so that genes annotated to child terms also counted toward parent terms. Separate over-representation analyses were run for UP and DOWN sets within each domain. For each term, a one-sided Fisher exact test was applied to a 2 × 2 contingency table comparing, for the term in question, the number of foreground genes recursively annotated to the term versus foreground genes not annotated to it, against the corresponding counts among background genes; the test was competitive and one-sided for over-representation. Terms were tested when at least 3 foreground genes were recursively annotated, and redundant parent terms with foreground counts identical to a child term were excluded. P values were adjusted by the Benjamini–Hochberg method separately within each direction and domain, with q ≤ 0.01 as the significance threshold.

*Sunburst plots.* Display the SynGO ontology hierarchy was generated in R. Filled sectors represent terms containing foreground genes, colored by the mean foreground log2FC of recursively annotated genes; unfilled sectors represent terms without sufficient foreground annotation ***(Supplementary Figure 42), (Supplementary Figure 43)***. Figures used data.table 1.18.4, jsonlite 2.0.0, dplyr 1.2.1, tidyr 1.3.2, ggplot2 4.0.3, ggforce 0.5.0, patchwork 1.3.2, scales 1.4.0, and ggrepel 0.9.8. SynGO was cited per portal recommendation (Koopmans et al., 2019, Neuron).

**Cross-Modality (Integrated) Pathway Analysis**

Integrated pathway figures combined five sources for the CRH-neuron OUD-versus-control contrast: GO BP, GO CC, GO MF, Reactome, and KEGG (positive NES/log2FC = higher in OUD; negative = higher in controls). GO inputs used the redundancy-reduced representative terms from the GO-only analysis; Reactome and KEGG inputs used the precomputed GSEA Wald-ranked result files. For each source, terms/pathways were filtered at nominal p < 0.05, and the integrated set retained the top 24 per source (24 GO BP, 24 GO CC, 24 GO MF, 24 Reactome, 24 KEGG; 120 total). GO terms were ranked by q value, nominal p value, then absolute NES; Reactome by nominal p value, q value, then absolute NES; KEGG by q value, nominal p value, then absolute NES. Leading-edge genes were parsed from `core_enrichment`, mapped from Entrez IDs to symbols via org.Hs.eg.db, uppercased, and joined to pseudobulk DE statistics by symbol; gene-level log2FC was used for GOChord coloring and nominal gene-level p value for DE-validated filtering.

For the integrated enrichment map, nodes represented retained GO terms, Reactome pathways, or KEGG pathways. Edges connected pairs sharing at least 1 leading-edge gene or with Jaccard overlap ≥ 0.025 (fallback Jaccard cutoff 0.018 if fewer than 40 edges were recovered) and were ranked with cross-source edges prioritized, then by shared gene count and Jaccard overlap (max_edges = 320; final network, 320 edges). Node fill encoded NES, node size encoded −log10(nominal p value), node shape encoded source class, and node border color encoded source/ontology (GO BP #00FFFF, GO CC #9933FF, GO MF #FF33FF, Reactome #FFB000, KEGG #0072B2). Louvain community detection, weighted by Jaccard overlap, and Fruchterman–Reingold layout (seed 149, 4,000 iterations) resolved 8 modules; module labels were generated from frequent non-generic terms in member descriptions, module hulls were drawn as transparent colored boundaries, and labels were placed with ggrepel (label seed 1491) ***(Figure 4)***.

Recurring leading-edge gene heatmaps used the top 10 terms/pathways per source and up to 56 genes ranked by number of source modalities, recurrence across terms/pathways, maximum absolute NES, best term p value, and gene symbol; a DE-validated heatmap retained only genes with nominal pseudobulk p < 0.05, and a multimodal DE-validated heatmap additionally required genes to appear in at least 2 source modalities ***(Supplementary Figure 4), (Supplementary Figure 5)***. Biological themes were assigned by keyword matching of GO, Reactome, and KEGG descriptions into high-level categories (synaptic/neuronal signaling; ion transport/metabolism; vesicle/ER-Golgi/endosome; adhesion/cytoskeleton/ECM; RNA processing/ribosome; immune/antigen; receptor/kinase signaling; development/morphogenesis; cell cycle/death/stress; other), with tile color encoding mean NES per theme-source bin and labels reporting retained counts; a companion version added an average-NES column computed as the unweighted mean of populated source-level mean NES values across the five sources ***(Supplementary Figure 3)***.

GO semantic-similarity modules were computed only within GO ontologies (Wang similarity, cutoff 0.70), and Reactome and KEGG pathways were then projected onto GO semantic modules by shared leading-edge genes (minshared = 2 or Jaccard ≥ 0.025); the semantic comparison displayed the top 14 GO semantic modules ranked by assigned source count, total assigned term/pathway count, and minimum p value. Integrated GOChord analyses (`GOplot::circledat`, `chord_dat`, `GOChord`) used term/pathway sectors for selected GO, Reactome, or KEGG entries, ribbons for leading-edge gene membership, and gene sectors colored by pseudobulk log2FC. Direction-specific panels were generated for OUD-enriched and control-enriched entries (nominal p < 0.05; up to 2 per source, with fallback selection if fewer than 4 were available) ***(Supplementary Figure 44)***, and module-driven panels were generated from Louvain modules with at least 4 mapped terms/pathways (up to 7 terms/pathways and 26 genes per module) ***(Supplementary Figure 45)***. GOChord parameters were space = 0.02, gene.order = "logFC", gene.space = 0.34, gene.size = 3.55–3.65, nlfc = 1, lfc.min = −2, and lfc.max = 2, with unique sector colors by term/pathway ID.

Cross-pathway alluvial visualizations were generated for CRH neuron pathway enrichment results comparing OUD patients with controls. Nominally significant GO BP, GO CC, GO MF, Reactome, and KEGG terms/pathways were retained using nominal pathway `p < 0.05`. Positive NES values were coded as higher enrichment in OUD, whereas negative NES values were coded as higher enrichment in controls ***(Figure 5a;5b), (Supplementary Figure 46)***. For the pathway-level alluvial summary, all nominally significant terms/pathways were exported to CSV (`n = 346`), and a readable plotting subset was selected using the top 2 terms/pathways per enrichment direction and source/ontology. Alluvial ribbons connected enrichment direction, pathway source/ontology, keyword-derived biological theme, and term/pathway identity. Ribbon width was proportional to capped `-log10(nominal pathway p value)`.

For gene-linked alluvial analyses, leading-edge/core genes from nominally enriched GO, Reactome, and KEGG terms/pathways were joined to the CRH neuron pseudobulk differential expression table. Genes were retained for visualization if they had non-missing log2 fold change and nominal pseudobulk `p < 0.05`. Positive pseudobulk log2FC indicated higher expression in OUD, and negative log2FC indicated higher expression in controls. General DE-supported alluvial panels selected up to 6 genes per direction and up to 2 term/pathway links per gene ***(Figure 5a;5b)***. Multi-modal gene panels retained genes represented in leading-edge/core sets from at least two pathway-analysis source columns and plotted up to 5 genes per direction with up to 2 links per gene ***(Supplementary Figure 47)***.

**Targeted Differential Expression Visualization of Selected Genes**

Pseudobulk differential expression results were generated upstream and provided as a DESeq2-style table (gene symbol, log2FC, nominal p value, adjusted p value). For the targeted visualization, the pseudobulk DEG list was filtered to a prespecified set of nine genes (AGRP, VIP, NPY, OXT, GAL, CRHBP, GHR, PRLR, CALCR); no additional genes were included. For each gene, the source log2FC was used as the effect-size estimate and the source nominal p value as the significance value, with −log10(nominal p value) computed for the volcano-style panel. Adjusted p values were retained in the exported filtered table but were not used for color scaling. Figures were generated in R using ggplot2 4.0.3, ggrepel 0.9.8, patchwork 1.3.2, dplyr 1.2.1, readr 2.2.0, and scales 1.4.0.

The volcano-style panel plotted log2FC on the x-axis against −log10(nominal p value) on the y-axis. A dashed horizontal line marked nominal p = 0.05, and a vertical reference line marked log2FC = 0. Points were colored by direction of effect only (log2FC > 0 in green #006400; log2FC < 0 in red #B40426), not by FDR status or a fold-change threshold. All nine prespecified genes were plotted and labeled with ggrepel (seed = 7, max.overlaps = Inf). The targeted heatmap displayed the same genes ordered by descending log2FC, with tile color representing pseudobulk log2FC only ***(Figure 5c)***.

**Supplementary Results**

**Cross-atlas validation of the CRH-enriched population**

Independent human HYPOMAP and mouse PVN Atlas analyses confirmed the CRH-enriched identity of Splat_410_841 and Splat_410_843; detailed results are presented below.

**Splat_410 characterization and human HYPOMAP-guided CRH population definition (Pipeline 1)**

The Splat_410 cluster comprised 16,596 cells across 12 subclusters (Siletti et al. taxonomy aliases 724, 725, 833, 834, 837–844), all predominantly hypothalamic in origin (66–98% hypothalamus per subcluster in the reference atlas). AVP was detected in 57.5–88.2% of cells across all 12 subclusters, with 12- to 75-fold enrichment for hypothalamic markers relative to background (AVP ~75×, OXT ~53×, TH ~42×, CARTPT ~26×, CRH ~20×, SCGN ~19×, SIM1 ~16×, OTP ~14×, TRH ~12×). The master hypothalamic PVN developmental transcription factors *SIM1* and *OTP* were among the most highly enriched genes.

The 12 subclusters split into at least two signatures: a majority AVP/OXT-dominant, magnocellular-type signature and a smaller, transcriptionally distinct CRH+/PENK+ parvocellular-type signature concentrated in Splat_410_841 (~1,254 cells) and Splat_410_843 (~3,520 cells). In these subclusters, *CRH* was ~13-fold higher than in other Splat_410 subclusters, and PENK was expressed in ~46–51% of cells (versus <10–20% elsewhere).

Cross-validation against the independent human HYPOMAP atlas confirmed this identity: marker genes (CRH, CARTPT, AVP, PGF, TH) of HYPOMAP cluster C4-297 ("Mid-3 GLU-3 AVP PGF") directly corresponded to Splat_410_841 and Splat_410_843, aligning specifically with the glutamatergic CRH⁺ lineage. PENK was not among the top markers of C4-297, and a separate PENK-specific HYPOMAP cluster (C4-354, "Mid-2 GLU-2 PGR PENK") showed no marker overlap with AVP or CRH, suggesting *PENK* represents a secondary co-expressed feature rather than an independently defined population. Subcluster-level MMC mapping confidence was somewhat below the dataset average (median bootstrapping probability ~0.48 vs. ~0.62), but was retained without filtering given strong marker and cross-atlas support.

**Mouse PVN Atlas-guided SIM1-positive re-clustering (Pipeline 2)**

Nuclei with detectable SIM1 expression (normalized expression >0) were subset, yielding 2,847 SIM1-positive nuclei from 25 donors. Condition was derived from the Opioid metadata column, comprising 1,488 HC-associated nuclei from 15 donors and 1,359 OUD-associated nuclei from 10 donors. Median detected genes per nucleus was 3,407, median UMIs 6,310, and median mitochondrial percentage 1.05%. QC distributions for this subset are shown in ***(Supplementary Figure 48)***.

The first 30 PCs explained 91.1% of PCA variance ***(Supplementary Figure 49)***. Unsupervised clustering on Harmony-corrected embeddings (30 Harmony dimensions, k = 30 nearest neighbors, SNN resolution = 3.5) identified 22 de novo Harmony clusters ***(Supplementary Figure 16)***. Harmony correction reduced donor-associated structure (variance-weighted donor R² from 0.149 in PCA space to 0.099 in Harmony space; ***(Supplementary Figure 15)***.

Post hoc biological labels were assigned using a marker-based annotation strategy informed by the mouse PVN Atlas. For each cluster, normalized RNA expression was summarized for PVN Atlas-associated marker genes by calculating both average expression and the proportion of cells with detectable expression, converted to gene-wise relative enrichment scores across clusters. Primary markers included AVP, OXT, CRH, TRH, SST, PENK, GHRH, SLC17A6, ASB4, and BRS3. Secondary markers included ADARB2, NR3C1, SCGN, AGTR1, FOXP1, TAC1, TH, UCN3, NFIB, SFRP2, CALB2, and OMP ***(Figure 3c)***. Of the 22 clusters, 10 received high-confidence and 11 medium-confidence labels; these 21 clusters were retained for downstream analyses ***(Supplementary Figure 17)***. Harmony cluster 5 (164 nuclei) received a low-confidence Asb4-Adarb2-like label and was excluded from label-specific interpretation ***(Figure 3a)***.

Multiple Harmony clusters shared the same marker-defined identity (e.g., Avp-Th: clusters 2, 4, 11, 16; Slc17a6-Adarb2: clusters 7, 13, 14), indicating greater transcriptional heterogeneity within human PVN cell types than predicted by the mouse atlas, where these populations resolve as single clusters. Donor-, condition-, and label-stratified cell counts are provided in ***(Supplementary Figure 50)***.

Species-specific differences in PVN cell-type definitions. Several discrepancies between mouse PVN Atlas predictions and the present human data emerged through re-clustering. The mouse atlas defines CRH neurons as a single Crh-Nr3c1 population co-expressing NR3C1 with an ADARB2-co-expressing subset; in the human data, ADARB2 was detectable in only 20.6% of Harmony cluster 1 nuclei, suggesting the Crh-Adarb2 subtype is less prominent or less transcriptionally distinct in human PVN ***(Figure 3b; 3d)***. The mouse atlas separates CRH, AVP, OXT, TRH, and SST into discrete clusters, whereas multiple human Harmony clusters mapped to the same marker-defined identity, indicating greater human transcriptional heterogeneity. The glutamatergic versus GABAergic CRH⁺ distinction emphasized by human HYPOMAP is not a primary organizing feature of the mouse atlas; the CRH+/PENK+ subclusters aligned specifically with the glutamatergic lineage; a distinction resolvable only through the parallel human atlas-referenced pipeline.

**Convergence of parallel pipelines on Splat_410_843**

The CRH-associated population in Pipeline 2 localized to Harmony cluster 1 (199 nuclei; high-confidence Crh-Nr3c1 label). CRH was detected in 57.8% of nuclei, NR3C1 in 56.3%, and ADARB2 in 20.6% ***(Figure 3b; 3d)***. Violin plots across all Harmony clusters further supported this identity assignment ***(Supplementary Figure 1a)***.

Post hoc cross-referencing with Allen MMC annotations confirmed that Harmony cluster 1 mapped predominantly to the Splat-*410* lineage: 162 of 199 nuclei (81.4%) at the broad cluster level ***(Supplementary Figure 18)***, and 110 of 199 (55.3%) specifically to Splat_410_843 at the subcluster level, accounting for 53.9% of all Splat_410_843 nuclei in the dataset ***(Supplementary Figure 19).***

Thus, both the human HYPOMAP-guided pipeline (defining CRH+/PENK+ neurons by Splat_410_841 and Splat_410_843 subcluster assignment) and the mouse PVN Atlas-guided pipeline (identifying CRH-enriched neurons through de novo SIM1-positive re-clustering) independently converged on Splat_410_843 as the principal CRH-enriched PVN subcluster ***(Figure 3)***.

Donor-level representation of Harmony cluster 1 is shown in ***(Supplementary Figure 51)***. In the OUD-control analytic subset, Harmony cluster 1 contained 193 nuclei, including 106 control-associated and 87 OUD-associated nuclei. Median cluster 1 nuclei per donor was 5 in controls and 3 in OUD, and no individual donor contributed ≥500 nuclei.

**Exploratory CRH neuron composition analysis**

As an exploratory donor-level composition check within the SIM1-positive PVN subset, CRH cluster 1 proportions did not differ significantly between OUD and controls ***(Supplementary Figure 14)***. Median CRH cluster 1 proportion was 4.48% in controls and 3.95% in OUD. Neither a condition-only quasibinomial model (P = 0.288; Benjamini-Hochberg adjusted P = 0.627) nor a Wilcoxon rank-sum test (P = 0.420) identified a significant difference.

**Detailed pathway enrichment analyses**

**Gene set enrichment analysis**

Preranked GSEA was performed using two ranking metrics: the Wald statistic and shrunken log₂ fold-change. Wald-ranked *GSEA* GO analysis identified 251 Biological Process (BP) terms, 67 Molecular Function (MF) terms, and 50 Cellular Component (CC) terms at nominal P < 0.05 after semantic simplification. FDR support varied by ontology: 20 BP, 3 MF, and 4 CC terms reached q < 0.05; 127 BP, 21 MF, and 35 CC terms reached q < 0.25. KEGG identified 53 nominally enriched pathways (11 at q < 0.05; 53 at q < 0.25). Reactome identified 107 nominally enriched pathways (1 at q < 0.05; 2 at q < 0.25). Nominal and FDR-supported pathway counts are summarized in ***(Supplementary Figure 52)***. GO analysis identified 154 BP, 54 MF, and 47 CC terms at nominal P < 0.05 after semantic simplification. KEGG identified 17 of 22 tested and Reactome 77 of 84 tested nominally significant pathways. Shrunken log₂FC-ranked not used in further results because the Wald-ranked analysis was prioritized because the Wald statistic incorporates both effect size and estimation uncertainty, providing a more robust ranking for pathway-level inference in DESeq2-based workflows.

**Biological theme analysis of GO enrichment**

Significant GO terms from Wald-ranked GSEA were assigned to biological themes by keyword-derived categories, and mean NES was calculated across themes and ontologies ***(Supplementary Figure 20)***. Themes enriched in OUD included developmental/morphogenesis, synaptic/neuronal signaling, receptor/kinase signaling, adhesion/cytoskeleton/ECM, ion transport/metabolism, and immune/antigen processes. Themes suppressed in OUD included vesicle/ER-Golgi/endosome and RNA processing/ribosome functions.

An enrichment map was constructed after semantic redundancy reduction (Wang similarity ≥ 0.70) using leading-edge gene overlap (Jaccard ≥ 0.025) and Louvain community detection ***(Supplementary Figure 23)***. Five functional modules emerged: two OUD-enriched modules dominated by muscle/proliferation/receptor, actin/junction/neuron, and three control-enriched modules encompassing exosome/RNase/transferase, Golgi/vesicle/endoplasmic, and endosome/late/lysosomal semantic terms.

**Recurrent leading-edge gene analysis GO GSEA**

Leading-edge genes across the top 20 GO BP, CC, and MF terms (ranked by q-value, then P-value, then |NES|) were identified and ranked by cross-ontology recurrence and effect size ***(Supplementary Figure 21).*** Cross-validation against pseudobulk DE results (nominal P < 0.05) retained 10 of 54 recurrent genes ***(Supplementary Figure 22)***: BAIAP2, ARC, RGS2, NECTIN1, and LRFN5 (positive NES; upregulated in OUD) and GOPC, BET1, EIF4A1, GOLT1B, and PREB (negative NES; downregulated in OUD).

**GOChord direction-specific GO GSEA pathway analysis**

GOChord analysis of direction-stratified enriched terms ***(Supplementary Figure 24)*** revealed that OUD-enriched pathways include cell junction assembly, tissue morphogenesis, actin/integrin/calmodulin binding, and neuronal synaptic functions (membrane, postsynaptic, and neuron-to-neuron interactions). Control-enriched pathways converged on catalytic RNA activity and RNA helicase function, lysosomal and endosomal activity, ER-to-Golgi vesicular transport, ribosome biogenesis, and transferase activity.

**Semantic-topic GOChord analysis of CRH neuron GO enrichment modules**

Semantic-topic decomposition of the CRH+-PENK+ neuron GO enrichment map resolved two positively enriched modules in OUD (positive mean NES) and two enriched modules in controls (negative mean NES). Each GOChord displayed the module's top 7 GO terms and 24 leading-edge genes ***(Supplementary Figure 25).*** OUD enriched terms included: Muscle/ proliferation/ receptor (Semantic topic 1) and Actin/ Junction/ Filament (Semantic topic 2). Control enriched terms included: Exosome/ RNAase/ Transferase (Semantic topic 3) and Golgi/ Endoplasmic/ Reticulum (Semantic topic 4).

Together these findings could indicate upregulation of pathways related to morphogenesis/ receptor signaling and actin-junction/synaptic structure in OUD. Downregulated pathways in OUD could include those related to RNA-processing/exosome activity and ER-Golgi/ endolysosomal trafficking.

**Reactome pathway analysis**

The Reactome input contained 107 pathways, all met nominal pvalue < 0.05, and 1 pathway met qvalue < 0.05. Across all Reactome pathways, 67 had positive NES values and 40 had negative NES values ***(Supplementary Figure 53)***.

The enrichment map displayed 72 nominally enriched Reactome pathways connected by 168 shared-core-gene overlap edges. Louvain community detection identified nine modules. OUD-enriched (positive mean NES) modules were Apoptotic/Diseases/Neurodegenerative, Extracellular/Matrix/Degradation, Notch1/Signaling/Transcription, Release/Canonical/cGMP, and Erythrocytes/Aquaporins/Carbon. Control-enriched (negative mean NES) modules were Cycle/GTPase/ICOS, Polymerase/Promoter/Transcription, Glycan/Golgi/Anterograde, and rRNA/Cytosol/Nucleus (***Supplementary Figure 34)***.

The direction-specific NES heatmap displayed 56 top-ranked nominal Reactome pathways ***(Supplementary Figure 35)***. The recurring leading-edge/core gene heatmap displayed 56 recurrent core genes across 36 selected Reactome pathways ***(Supplementary Figure 36).*** The pseudobulk DE-validated recurring gene heatmap retained 28 genes across 20 Reactome pathways after filtering to genes with pseudobulk DE pvalue < 0.05 ***(Supplementary Figure 37)***.

The direction-specific Reactome GOChord analysis selected 14 Reactome pathways for direction-stratified visualization. After gene/pathway representation filtering, the OUD-enriched panel displayed 4 pathways and 28 genes, while the control-enriched panel displayed 6 pathways and 28 genes ***(Supplementary Figure 38)***. Module-driven GOChord analysis used 9 Louvain modules and 47 selected module pathways. The combined module GOChord figure displayed the top 4 modules, and standalone GOChord files were generated for 8 modules with sufficient pathway and gene representation ***(Supplementary Figure 39)***.

The biological-theme comparison summarized Reactome pathways by direction and high-level keyword-inferred theme ***(Supplementary Figure 40)***. Two themes were uniformly OUD-enriched: synaptic/neuronal signaling (7 pathways, all higher in OUD, mean NES = 1.711) and Cell cycle/ death/ stress. Two themes were uniformly control-enriched: vesicle/ER-Golgi/endosome (3 pathways, mean NES = −1.581) and RNA processing/ribosome (6 pathways, mean NES = −1.588).

**KEGG Pathway Analysis**

KEGG analysis retained 53 enriched pathways, of which 11 met q < 0.05. Among these, 39 had positive NES and 14 negative NES ***(Supplementary Figure 27), (Supplementary Figure 30)***. The strongest OUD-enriched pathways were gastric acid secretion, bile secretion, long-term potentiation, salivary secretion, viral myocarditis, glycosaminoglycan biosynthesis (heparan sulfate-heparin), and calcium signaling ***(Supplementary Figure 27), (Supplementary Figure 30), (30)***. The strongest control-enriched pathways were RNA polymerase, biosynthesis of nucleotide sugars, amino sugar and nucleotide sugar metabolism, galactose metabolism, lysosome biogenesis, terpenoid backbone biosynthesis, and fructose/mannose metabolism ***(Supplementary Figure 27), (Supplementary Figure 30), (Supplementary Figure 31)***. The convergence of nucleotide-sugar biosynthesis, amino/nucleotide-sugar metabolism, and fructose/mannose metabolism could indicate coordinated relative downregulation of glycosylation-substrate precursors in OUD CRH neurons.

The KEGG enrichment map contained 220 overlap edges connecting pathways with shared leading-edge genes and identified 5 Louvain modules ***(Supplementary Figure 26)***. The three positive modules centered on secretion/ion transport, adhesion/signaling, and cardiomyopathy/cytoskeleton-related pathways; the two negative modules concentrated on biosynthesis/metabolism/nucleotide and lysosome-related pathways, with mean NES of 1.55, 1.36, 1.55, −1.68, and −1.44, respectively. Module-driven GOChord visualization linked module-associated pathways to shared leading-edge genes across the top 4 modules ***(Supplementary Figure 32)***.

Keyword-inferred theme summaries showed enrichment across ion transport/secretion, synaptic/neuronal signaling, signaling pathways, metabolism/biosynthesis, immune/infection, adhesion/cytoskeleton, RNA/transcription, organelle/protein processing, and disease/tissue remodeling themes ***(Supplementary Figure 54)***. Ion transport/secretion contained 15 pathways (14 higher in OUD, 1 higher in controls), while metabolism/biosynthesis contained 9 pathways (7 higher in controls, 2 higher in OUD) ***(Supplementary Figure 54)***.

Recurring KEGG leading-edge genes were visualized before and after pseudobulk DE validation ***(Supplementary Figure 28), (Supplementary Figure 29)***. The unfiltered heatmap displayed 56 recurring genes across 32 pathways ***(Supplementary Figure 28)***; after filtering on pseudobulk DE nominal P < 0.05, 22 genes across 18 pathways were retained (Supplementary Figure 29): COL4A2, PGM1, NECTIN1, GMPPB, GAA, BAIAP2, LMNA, AQP1, OS9, CD63, RELN, ALDH7A1, MAP2K7, AP5Z1, PMAIP1, GSTO1, LRFN5, PREB, REL, COL27A1, GALNS, and VPS16. These span structural/ECM and adhesion (COL4A2, COL27A1, NECTIN1, RELN, LRFN5, LMNA, BAIAP2), glycosylation and glycan-processing/lysosomal metabolism (PGM1, GMPPB, GAA, GALNS, OS9, PREB, VPS16, CD63), stress/apoptosis and MAPK-inflammatory signaling (MAP2K7, PMAIP1, REL, GSTO1, ALDH7A1), and water/ion homeostasis (AQP1).

**SynGO Pathway Analysis**

SynGO analysis was performed on nominally significant genes (P < 0.05, |log₂FC| > 0.25). Of 77 upregulated and 171 downregulated genes submitted, 13 and 16 mapped to SynGO, respectively. Upregulated genes annotated to postsynaptic density, synapse assembly, synapse organization, and chemical synaptic transmission. Downregulated genes annotated to synaptic vesicle, synapse organization, synaptic signaling, and regulation of postsynaptic membrane neurotransmitter receptor levels ***(Supplementary Figure 42), (Supplementary Figure 43)***. No SynGO terms reached significance after Benjamini-Hochberg correction (q ≤ 0.01), consistent with the modest gene-level signal and limited CRH-neuron sample size.

**Cross-modality visualizations**

Direction-specific cross-pathway GOChord plots integrating GO, Reactome, and KEGG results ***(Supplementary Figure 44)*** identified OUD-enriched terms including cell junction assembly, tissue morphogenesis, actin binding, calmodulin binding, neuron-to-neuron synapse, postsynaptic specialization, neuronal system, and neurotransmitter release cycle. Control-enriched terms included catalytic activity acting on RNA, RNA helicase activity, lysosomal membrane, ER-to-Golgi vesicle-mediated transport, ribosome biogenesis, asparagine N-linked glycosylation, synthesis of substrates in N-glycan biosynthesis, and biosynthesis of nucleotide sugars.

Module-driven GOChord visualization ***(Supplementary Figure 45)*** identified upregulation of neuronal junction/secretory function, actin/integrin/adherens, and muscle/regulation/tissue modules in OUD, with concurrent downregulation of Golgi/endoplasmic/vesicular function.

Similarly, cross-pathway alluvial analysis summarized nominally enriched GO BP, GO CC, GO MF, Reactome, and KEGG terms/pathways in CRH neurons from OUD patients and controls ***(Supplementary Figure 46)***. Higher-in-OUD terms/pathways included: cell junction assembly and tissue morphogenesis; neuron to neuron synapse, postsynaptic specialization; GO MF: actin binding, calmodulin binding; Reactome: Neuronal System, Neurotransmitter release cycle; KEGG: Bile secretion; Gastric acid secretion ***(Figure 5a)***. Higher-in-controls terms/pathways shown: GO BP: endoplasmic reticulum to Golgi vesicle-mediated transport, ribosome biogenesis; GO CC: late endosome, lysosomal membrane; GO MF: catalytic activity, acting on RNA, RNA helicase activity, Asparagine N-linked glycosylation; Reactome: Synthesis of substrates in N-glycan biosynthesis; KEGG: Biosynthesis of nucleotide sugars, RNA polymerase ***(Figure 5b)***.

**Supplementary Figures and Tables**

Supplementary Table 1:

| Characteristic | Overall (N = 42) | HC (N = 22) | OUD (N = 20) | p-value |
| --- | --- | --- | --- | --- |
| Age (years) | 45 (14) | 51 (13) | 39 (13) | 0.007 |
| Sex |  |  |  | 0.7 |
| Female | 13 (31%) | 6 (27%) | 7 (35%) |  |
| Male | 29 (69%) | 16 (73%) | 13 (65%) |  |
| PMI (hours) | 27 (10) | 31 (9) | 23 (9) | 0.008 |
| Died of overdose | 16 (38%) | 0 (0%) | 16 (80%) | <0.001* |

*Died of overdose was tested using Fisher's exact test for completeness of reporting; because overdose is a near-definitional consequence of OUD status in this cohort rather than an independent covariate, this comparison is not interpreted as evidence of confounding and was not treated as a balance-check variable in the differential expression covariate design.

Cohort demographics by condition group. HC = healthy control; OUD = opioid use disorder. Values are mean (SD) or n (%). p-values: Wilcoxon rank-sum exact test (Age, PMI); Fisher's exact test (Sex, Died of overdose).

Supplementary Table 2:

| Donor | Condition | Nuclei (pre-filter) | Nuclei (post-filter) | % Retained | Median nCount (post-filter) | Median %MT (post-filter) |
| --- | --- | --- | --- | --- | --- | --- |
| 900 | HC | 15493 | 15449 | 99.7% | 1443 | 0.78 |
| 916 | OUD | 28576 | 22795 | 79.8% | 928 | 2.90 |
| 922 | OUD | 3690 | 3543 | 96% | 1092 | 0.93 |
| 926 | HC | 47742 | 7059 | 14.8% | 6875 | 3.27 |
| 931 | OUD | 6407 | 4753 | 74.2% | 566 | 1.84 |
| 933 | OUD | 7365 | 6913 | 93.9% | 1177 | 1.57 |
| 934 | HC | 9791 | 9776 | 99.8% | 1178 | 0.30 |
| 936 | HC | 10038 | 9872 | 98.3% | 1035 | 0.89 |
| 940 | HC | 3118 | 1618 | 51.9% | 381.5 | 1.95 |
| 945 | HC | 11513 | 11449 | 99.4% | 784 | 0.76 |
| 958 | OUD | 8290 | 8198 | 98.9% | 1319 | 0.83 |
| 960 | OUD | 13341 | 13301 | 99.7% | 1257 | 0.68 |
| 961 | OUD | 8771 | 8275 | 94.3% | 795 | 1.95 |
| 964 | HC | 5620 | 4435 | 78.9% | 1482 | 1.56 |
| 966 | HC | 12334 | 12332 | 100% | 876 | 0.27 |
| 967 | HC | 16155 | 16155 | 100% | 746 | 0.18 |
| 974 | OUD | 8138 | 8137 | 100% | 859 | 0.18 |
| 983 | OUD | 13999 | 13972 | 99.8% | 1026 | 0.65 |
| 987 | OUD | 2642 | 2544 | 96.3% | 351 | 0.91 |
| 992 | HC | 3011 | 2656 | 88.2% | 722.5 | 1.32 |
| 993 | HC | 7354 | 7302 | 99.3% | 990 | 0.66 |
| 995 | HC | 14436 | 13434 | 93.1% | 1190.5 | 1.63 |
| 996 | OUD | 4916 | 4856 | 98.8% | 821 | 0.43 |
| 201 | HC | 11061 | 10634 | 96.1% | 1229 | 0.96 |
| 203 | OUD | 6281 | 6280 | 100% | 1212.5 | 0.12 |
| 204 | HC | 1251 | 1227 | 98.1% | 324 | 0.40 |
| 206 | HC | 8085 | 6198 | 76.7% | 2798 | 1.56 |
| 211 | OUD | 7546 | 5530 | 73.3% | 2259 | 2.19 |
| 215 | HC | 8183 | 7121 | 87% | 1906 | 1.58 |
| 217 | OUD | 19093 | 18937 | 99.2% | 2752 | 0.77 |
| 218 | OUD | 8929 | 8560 | 95.9% | 1025 | 0.65 |
| 230 | OUD | 22323 | 20740 | 92.9% | 2364 | 1.72 |
| 234 | OUD | 12892 | 12874 | 99.9% | 2728 | 0.60 |
| 239 | HC | 3904 | 3485 | 89.3% | 917 | 1.11 |
| 244 | HC | 3787 | 3787 | 100% | 1057 | 0.14 |
| 245 | OUD | 7886 | 7721 | 97.9% | 863 | 0.64 |
| 248 | OUD | 3808 | 3793 | 99.6% | 1231 | 0.41 |
| 249 | HC | 3381 | 3379 | 99.9% | 1180 | 0.12 |
| 250 | OUD | 3057 | 3000 | 98.1% | 1120 | 0.47 |
| 251 | HC | 5631 | 5601 | 99.5% | 1215 | 0.32 |
| 252 | HC | 5992 | 5960 | 99.5% | 739 | 0.17 |
| 255 | HC | 2854 | 2784 | 97.5% | 478 | 0.61 |

Per-donor QC summary (n = 42 donors), pre- and post-mitochondrial-filtering. HC = healthy control; OUD = opioid use disorder. Full 11-column table (including median nFeature and pre-filter metrics) available as a standalone supplementary file.

Supplementary Figure 1a.

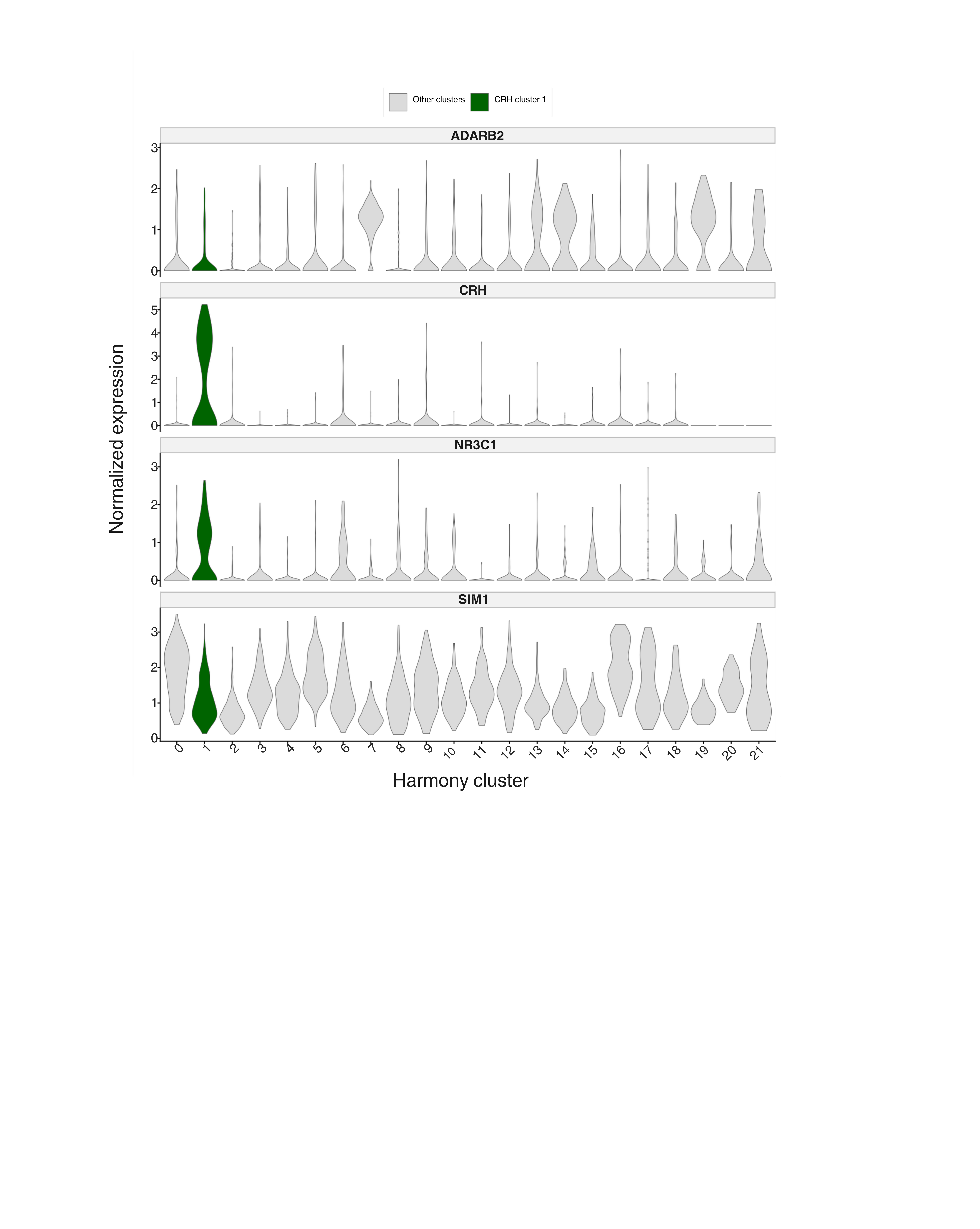

Marker validation of the CRH-enriched PVN cluster.

Violin plots showing normalized expression of key PVN identity markers across Harmony clusters within the SIM1-positive subset. Harmony cluster 1 showed high CRH expression with broad SIM1 and NR3C1 detection, supporting its assignment as a CRH-enriched Crh-Nr3c1 PVN population.

Supplementary Figure 1b.

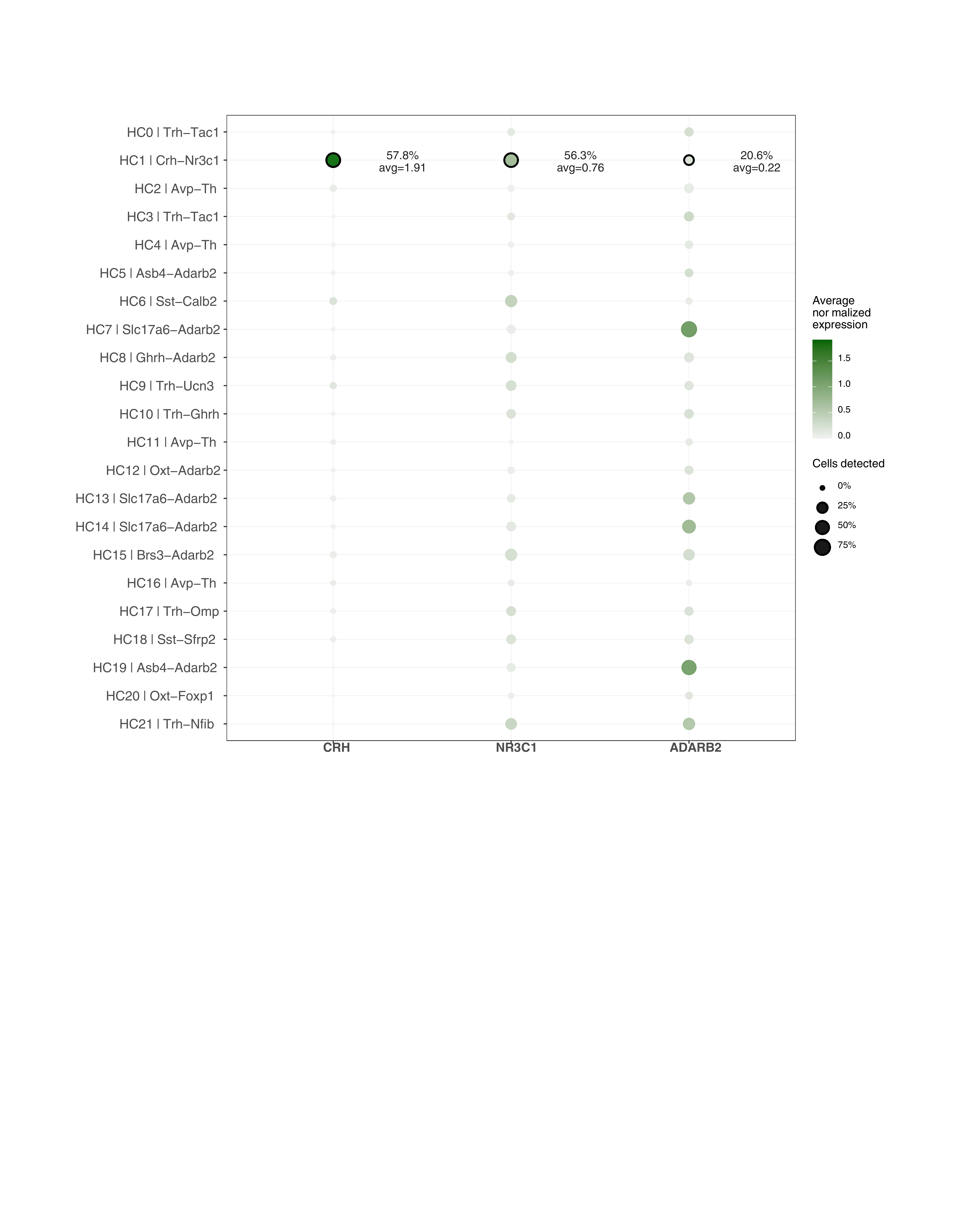

Marker support for the CRH-associated Harmony cluster.

Dot plot showing CRH, NR3C1, and ADARB2 expression across 22 Harmony clusters. Dot size indicates the percentage of cells expressing each marker and color indicates average normalized expression. Harmony cluster 1 showed the strongest combined CRH and NR3C1 expression, with detectable ADARB2 expression.

Supplementary Figure 1c

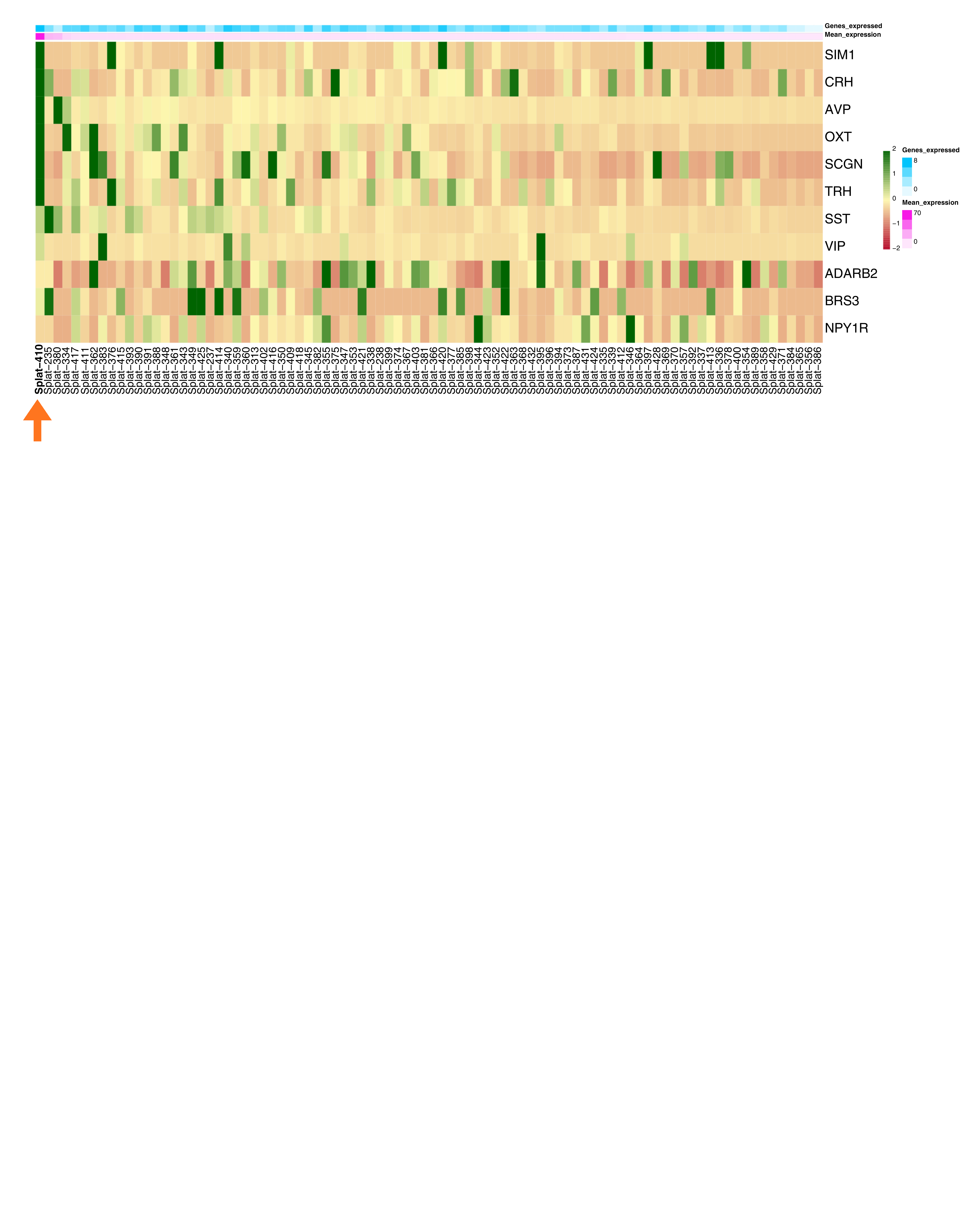

PVN marker expression across Splatter clusters.

Heatmap showing scaled average expression of PVN-associated markers across MMC Splatter clusters. Splat_410 showed the highest overall PVN marker expression, with SIM1 also enriched across several additional Splatter clusters.

Supplementary Figure 1d:

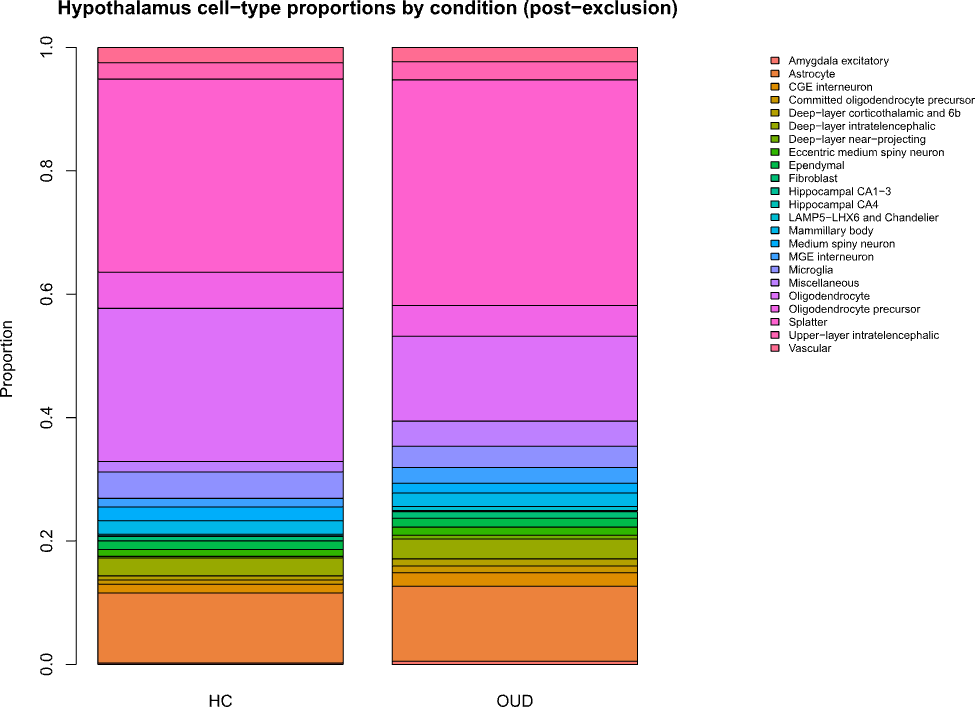

**Broad hypothalamic cell-type proportions are unchanged in opioid use disorder.**

Cell-type proportions do not differ between OUD and HC. Estimated proportions of each of 23 broad cell types, compared between opioid use disorder (OUD) and healthy control (HC) donors using propeller (logit-transformed proportions; donor, not cell, as the unit of replication). No cell type showed a statistically significant proportion difference after FDR correction (lowest FDR ≈ 0.19). This null result was robust to doublet filtering (complete agreement between all-cells and singlets-only analyses) and to per-donor doublet-rate differences by condition (not significant, Wilcoxon test). The Splatter category, containing Splat_410, the population of primary interest, showed no proportion shift (ratio ≈ 0.96, FDR ≈ 0.95), indicating that downstream expression-level findings in this population are not confounded by a change in its representation between conditions.

Supplementary Figure 2:

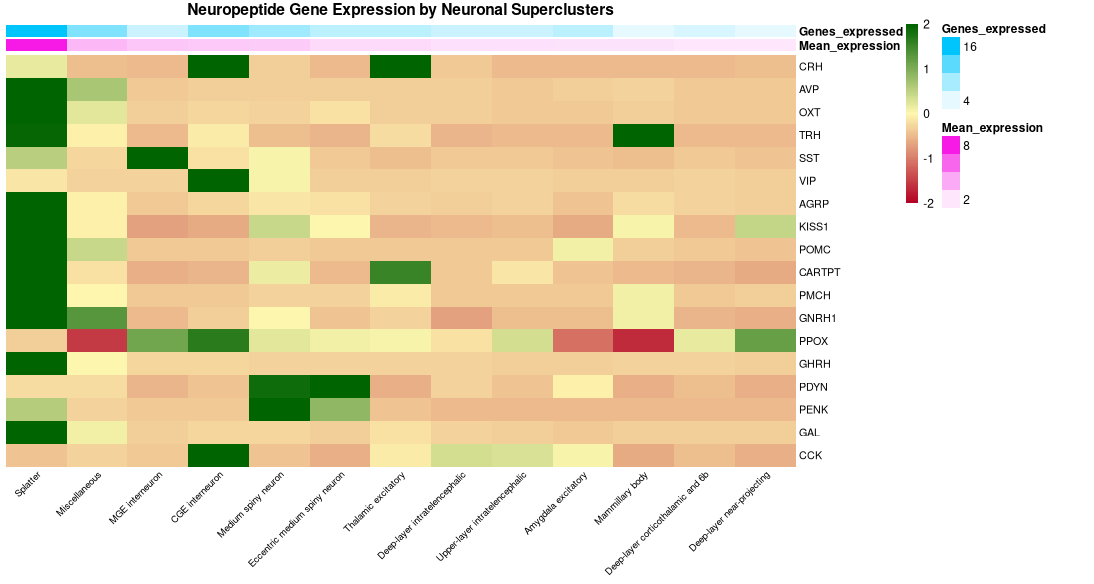

**Neuropeptide marker expression across neuronal cell types.**

Heatmap showing average normalized expression of neuropeptide marker genes of interest across MMC neuronal superclusters. Expression values were averaged by MMC supercluster, scaled per gene across superclusters, and expressed on Red to Green scale with average log2fold change ranging from -2 to 2 for visualization. Red indicates lower-than-average expression, yellow indicates average expression, and green indicates higher-than-average expression for each gene. Columns are ordered by decreasing mean neuropeptide marker expression. Top annotations show the unscaled mean marker expression and the number of neuropeptide marker genes expressed above 0.25 in each supercluster. Deep layer corticothalamic and 6b neurons and deep layer near projecting neurons notably are missing highly enriched expression in any neuropeptide markers

Supplement Figure 3:

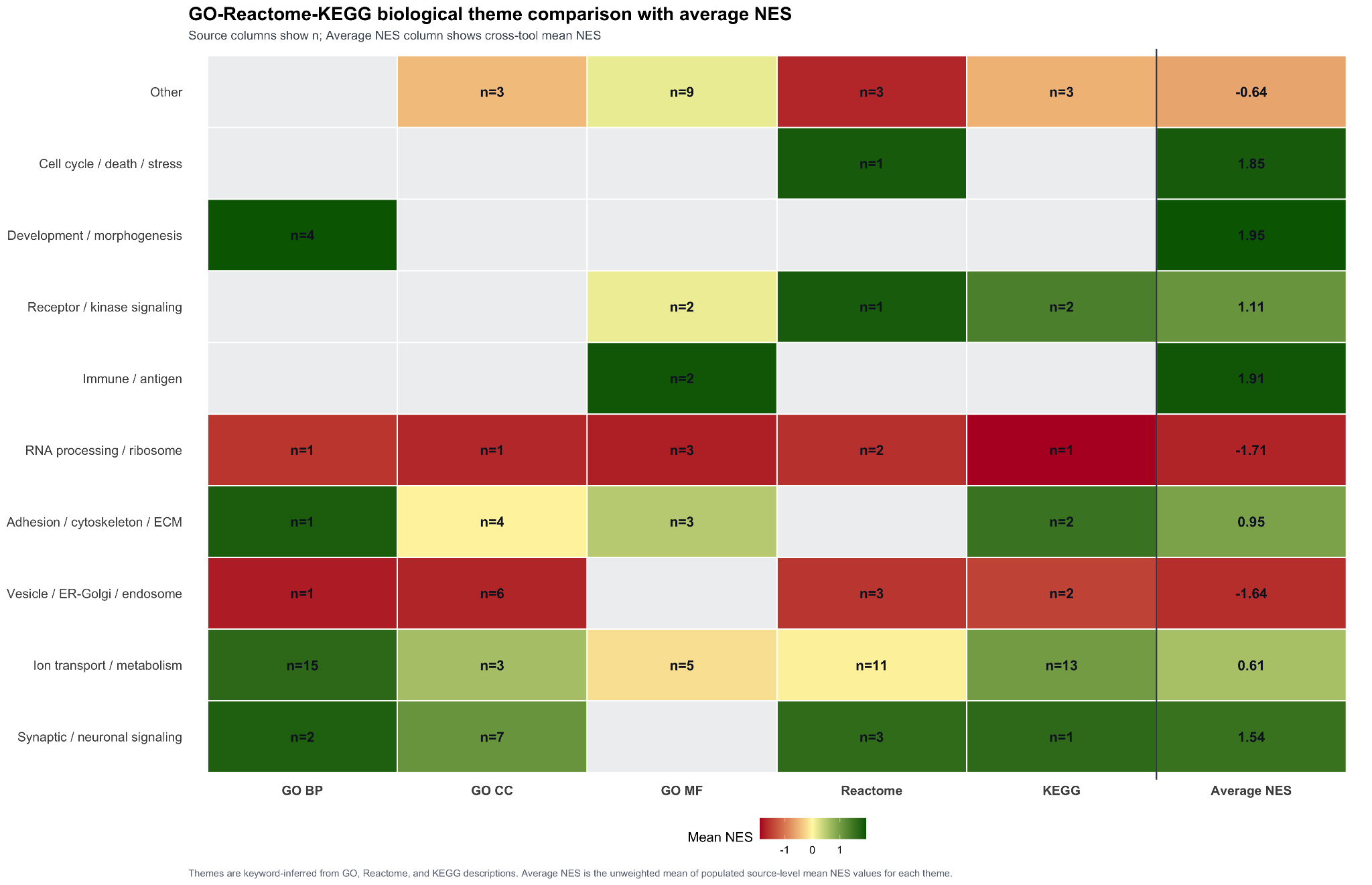

**GO-Reactome-KEGG Biological Theme Comparison with Cross-Tool Average NES.**

Heatmap summarizing enriched biological themes across GO Biological Process, GO Cellular Component, GO Molecular Function, Reactome, and KEGG pathway analyses in CRH neurons from the single-nucleus RNA-seq comparison of opioid use disorder patients versus controls. Rows represent keyword-inferred biological themes assigned from term/pathway descriptions. Source columns show the number of retained enriched terms/pathways assigned to each theme, while tile color represents the mean normalized enrichment score (NES) within that source-theme bin. Positive NES values indicate enrichment higher in OUD, and negative NES values indicate enrichment higher in controls. The rightmost column reports the cross-tool average NES for each biological theme, calculated as the unweighted mean of populated source-level mean NES values across GO BP, GO CC, GO MF, Reactome, and KEGG. Gray cells indicate no retained term/pathway for that source-theme combination.

Supplement Figure 4:

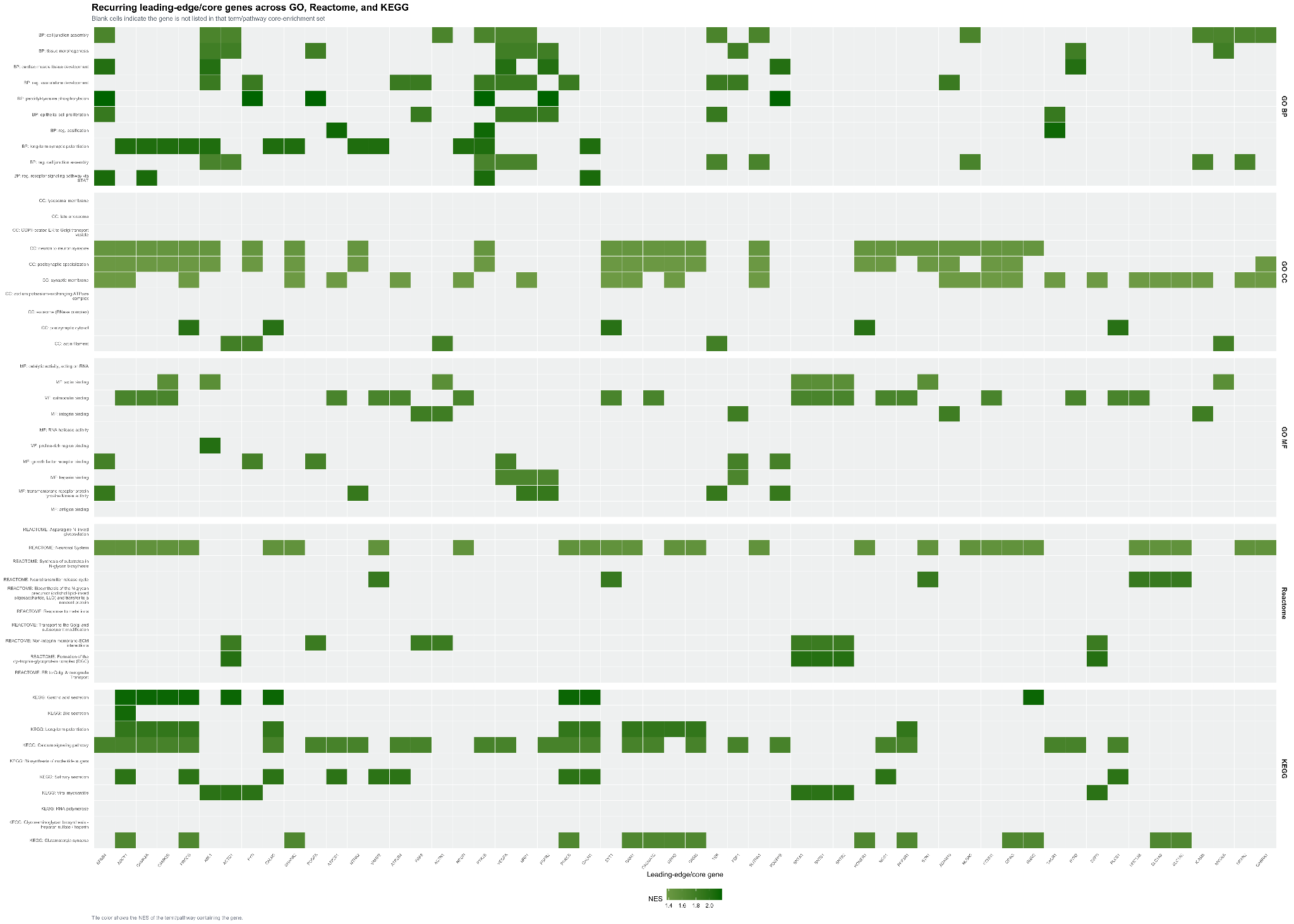

**Recurring Leading-Edge/Core Genes Across GO, Reactome, and KEGG in CRH Neurons.**

Heatmap showing recurrent leading-edge/core-enrichment genes across selected GO BP, GO CC, GO MF, Reactome, and KEGG enrichment results for CRH neurons comparing OUD patients with controls. Rows represent enriched GO terms or pathways, grouped by source, and columns represent recurring leading-edge/core genes. Colored cells indicate that a gene was present in the core-enrichment set for that term/pathway; blank cells indicate absence. Tile color represents the NES of the term/pathway containing the gene. The displayed matrix includes 50 terms/pathways, with 10 each from GO BP, GO CC, GO MF, Reactome, and KEGG, and 56 recurrent genes. Across the plotted matrix, 337 of 2,800 possible gene-term/pathway cells were populated.

Supplementary Figure 5:

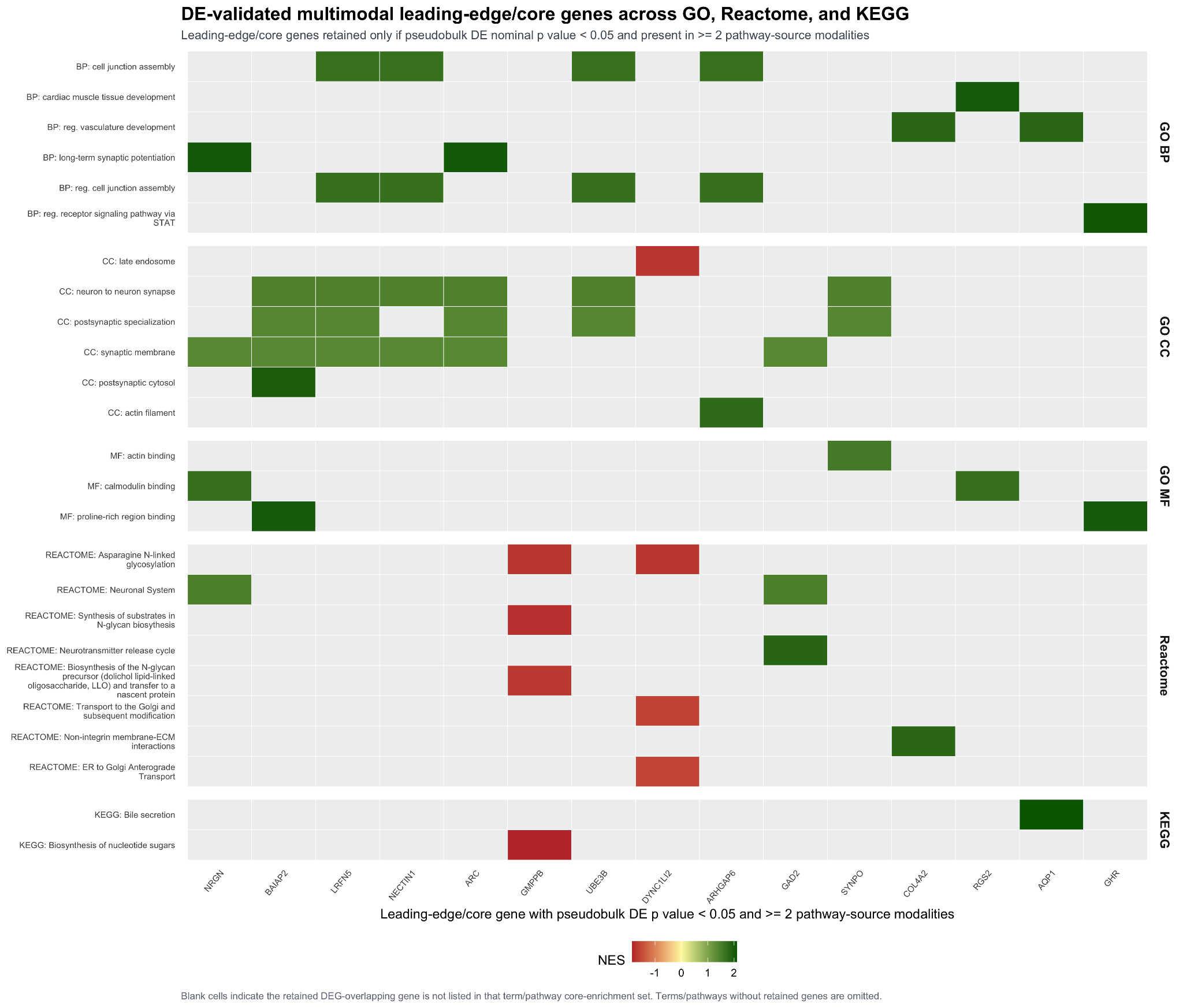

**DE-Validated Multimodal Leading-Edge/Core Genes Across GO, Reactome, and KEGG.**

Heatmap showing leading-edge/core-enrichment genes that were also nominally significant in pseudobulk differential expression analysis and recurred across at least two pathway-source modalities in CRH neurons. Genes were retained if they were present in the pseudobulk differential expression table with nominal p < 0.05 and appeared in leading-edge/core-enrichment sets from at least two sources among GO BP, GO CC, GO MF, Reactome, and KEGG. Rows represent enriched terms/pathways retained after filtering, grouped by source, and columns represent retained multimodal genes. Colored cells indicate that the gene was present in the core-enrichment set for that term/pathway; blank cells indicate absence. Retained terms/pathways included 6 GO BP, 6 GO CC, 3 GO MF, 8 Reactome, and 2 KEGG entries. The retained genes were NRGN, BAIAP2, LRFN5, NECTIN1, ARC, GMPPB, UBE3B, DYNC1LI2, ARHGAP6, GAD2, SYNPO, COL4A2, RGS2, AQP1, and GHR.

Supplementary Figure 6:

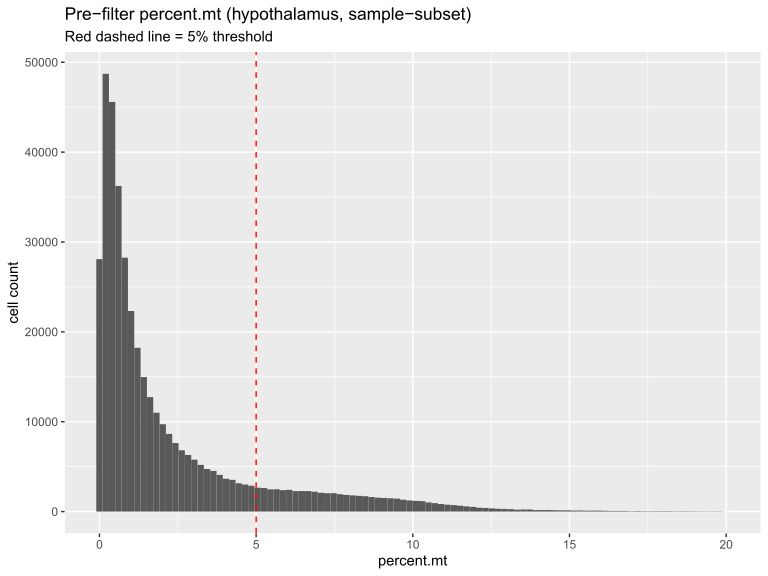

**Mitochondrial read distribution and secondary quality-control threshold.**

Mitochondrial content distribution across nuclei prior to secondary filtering. Histogram of percent mitochondrial reads per nucleus across all 408,684 nuclei (post-donor-exclusion). Nuclei exceeding 20% mitochondrial content had already been excluded during initial object assembly by the sequencing core; the distribution shown is therefore truncated at that upstream threshold, and summary statistics are conditional on it. The 5% threshold applied in this study is marked (dashed line). Median 1.063%, mean 2.311%, IQR 0.410–2.970%; 84.77% of the nuclei shown passed the 5% threshold and were retained.

Supplementary Figure 7:

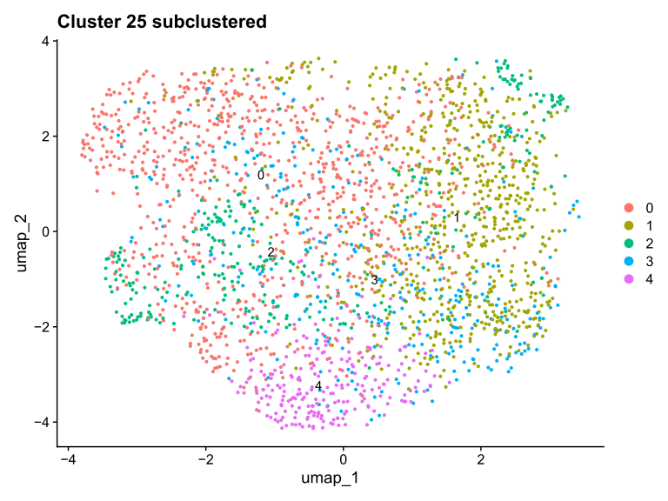

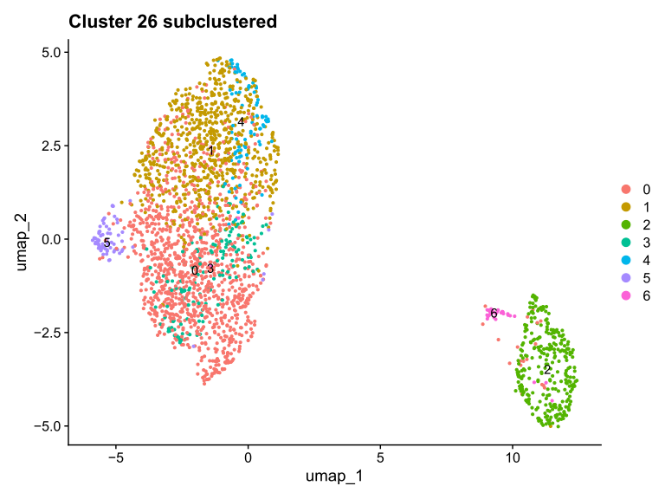

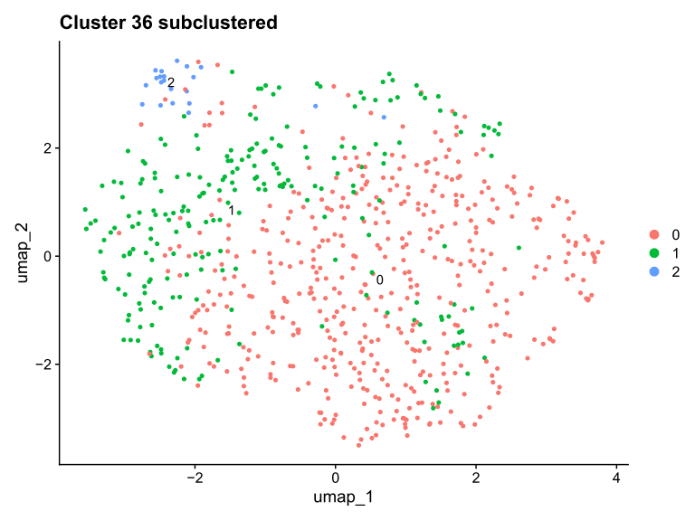

**Re-clustering and doublet identification in high-doublet-fraction clusters.**

Re-embedded and re-clustered UMAP of cells from target clusters identified for doublet fractions (44–56% compared to the dataset mean of 10.62%). Three clusters with high doublet fractions were labeled according to the new sub-community IDs, which were identified after re-running PCA, UMAP, and FindClusters on that subset. These clusters mostly consisted of doublets or stress artifacts. Cells flagged by scDblFinder were removed (2,616 nuclei across the three clusters.

Supplementary Figure 8:

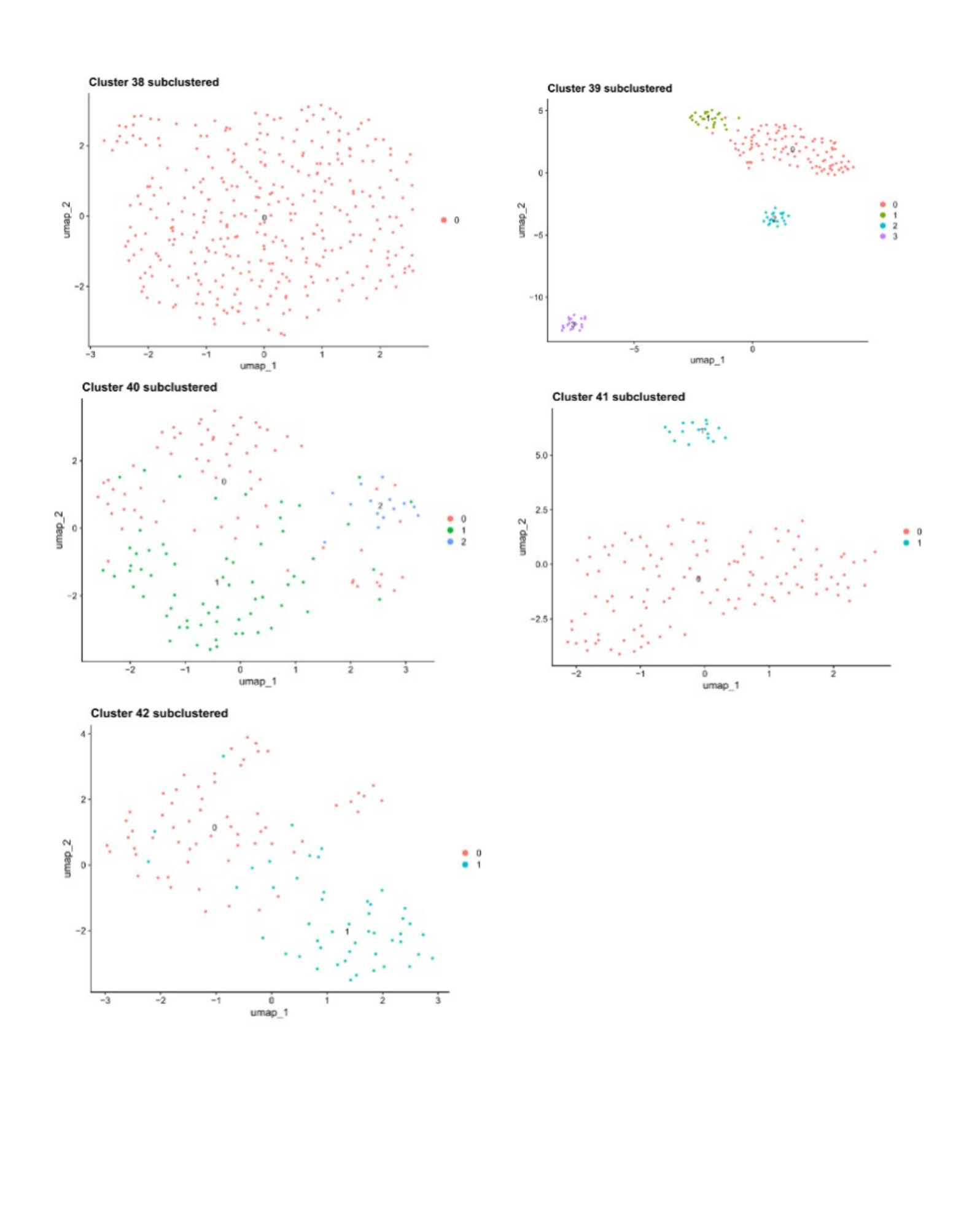

**Re-clustering of donor-dominance-flagged clusters.**

Targeted subcluster diagnostics of donor-dominance-flagged clusters 38–42. Five Louvain clusters were chosen for further analysis because each was dominated by one or two donors (cluster 38: 88% from a single donor; 39: 97% and 40: 100% from the same two donors; 41: 99% and 42: 97% each from a single donor), rather than because of elevated doublet fractions. Each panel displays cells from one cluster, which were re-embedded and re-clustered separately using independent PCA, UMAP, and FindClusters on that subset). Cells are colored by sub-community identities within each cluster. The coordinate spaces and sub-community labels are specific to each panel and are not directly comparable across panels or with the overall UMAP embedding (see Fig. 26). Clusters 39–42 split into 2–4 sub-communities with distinct marker profiles, including glial, endothelial, immune, and neuronal signatures, while cluster 38 remained unsplit (modularity = 0.60), and its identity is unresolved. None exhibited the doublet-driven profile that led to pruning in Figure 24, as their doublet fractions ranged from 2.7% to 19.4%, compared to 44%–56% for the pruned clusters and a dataset average of 10.62%. No cells were removed. Cluster 42 is predominantly from donor 67926, whose nuclei showed unusually low mitochondrial-filter survival, suggesting a possible technical factor influencing its composition.

Supplementary Figure 9:

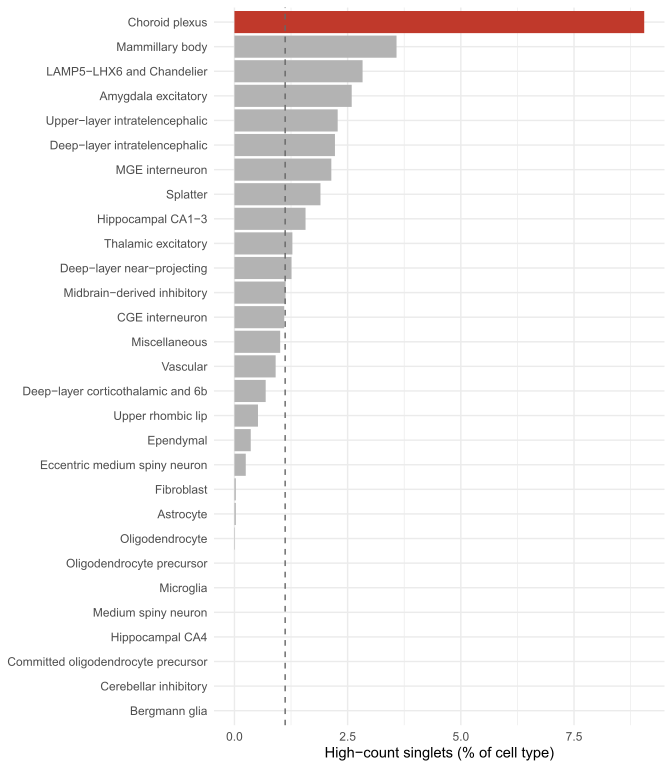

**High-count singlet nuclei concentrate specifically in choroid plexus.**

Percentage of each cell type’s nuclei with elevated UMI counts (nCount_RNA > 20,000) that were nonetheless classified as singlets by scDblFinder, plotted by annotated cell type (categories with fewer than 100 nuclei excluded). Choroid plexus shows a markedly elevated rate (9.05%, red) relative to the dataset-wide average (1.12%, dashed line) and more than double the next-highest category (Mammillary body, 3.58%; all other categories fall within a narrow 0-3.6% range. Choroid plexus epithelial cells are well documented as an unusually high-secretory cell type (Damkier, Brown & Praetorius, 2013; Thouvenot et al., 2006), though this literature establishes secretory and proteomic capacity rather than confirmed elevated per-nucleus transcript counts specifically; this remains a plausible, unverified explanation. Choroid plexus was in any case already excluded from between-group comparisons on donor-concentration grounds (see Methods). Splatter—containing Splat_410—sits close to the dataset-wide rate (1.90%), and its CRH+/PENK+ subset specifically falls below background (0.42%, not shown), indicating this phenomenon does not concentrate in the cell populations central to this study’s findings.

Supplementary Figure 10:

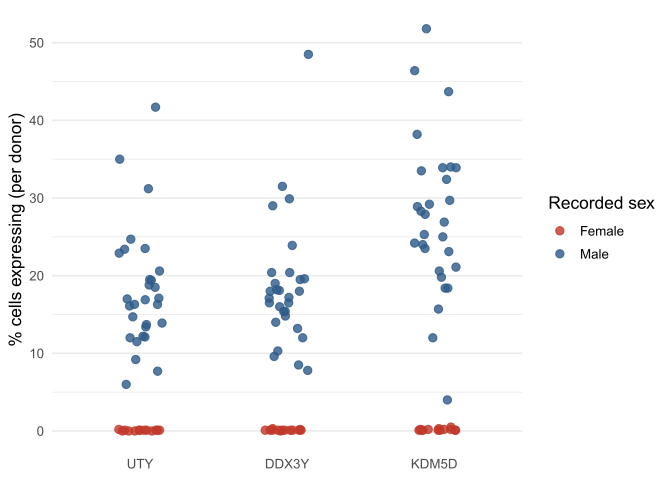

**Donor-level sex-gene concordance is consistent across all 42 donors.**

The percentage of cells expressing each of the three Y-chromosome genes—UTY, DDX3Y, and KDM5D—is shown per donor, with colors indicating recorded sex. Female donors cluster near background or ambient levels across all three genes (0.0-0.5%), distinctly separated from male donors (4.0-51.8%), with no borderline or misclassified cases. XIST and RPS4Y1 were not included in this dataset’s gene set and are therefore not displayed (see Supplementary Methods).

Supplementary Figure 11:

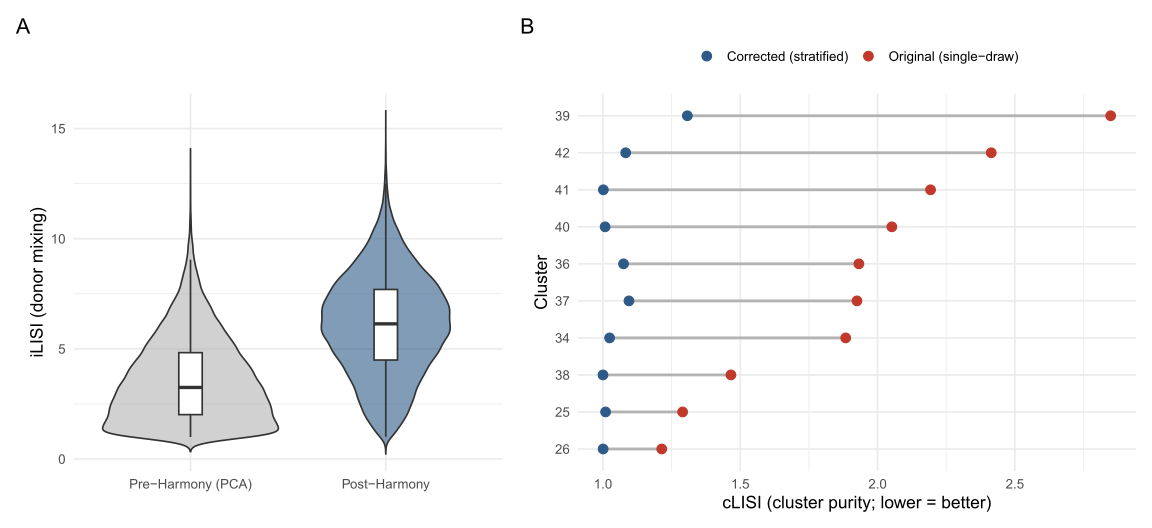

**Assessment and correction of batch integration quality and down sampling artifacts.**

(A) iLISI (donor mixing) calculated on a stratified cell sample, shown before (pre-Harmony, PCA embedding) and after (post-Harmony) batch integration. The median iLISI rose from 3.429 to 6.136, representing about an 89% improvement, with the upper distribution also increasing (max 13.424 to 15.024). This demonstrates a meaningful enhancement in donor mixing, not just a shift in the central tendency. (B) Per-cluster cLISI (cluster purity; lower indicates less local mixing) for ten clusters flagged during quality control diagnostics, comparing the initial single-draw calculation (red) to a stratified, full-inclusion calculation across five replicate seeds (blue). The original single-draw approach substantially inflated apparent cLISI for small clusters due to a down sampling artifact—cluster 39 saw the largest correction (2.849 to 1.307)—while cluster 38 remained almost unchanged (1.466 to 1.000). The corrected values, averaged over five seeds with SD < 0.013, demonstrate high reproducibility of the stratified correction.

Supplementary Figure 12:

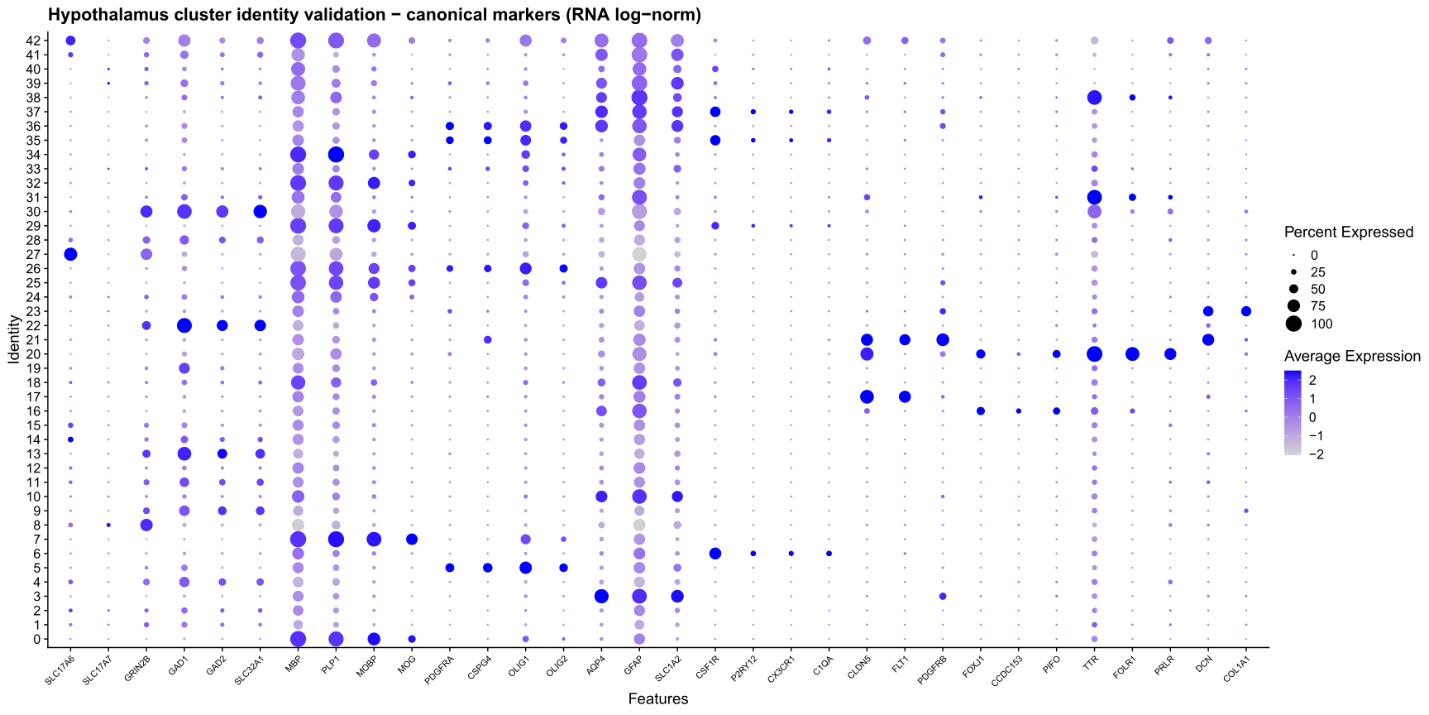

**Canonical marker validation of glial and non-neuronal cell identities.**

A dot plot shows expression levels of canonical marker genes across cell types, verifying glial and non-neuronal identities: oligodendrocytes (MBP, PLP1, MOBP, MOG), astrocytes (AQP4, GFAP, SLC1A2), microglia (CSF1R, P2RY12, CX3CR1, C1QA), vascular cells (CLDN5, FLT1, PDBFRB), ependymal cells (FOXJ1, CCDC153, PIFO), and choroid plexus cells (TTR, FOLR1, PRLR).

Supplementary Figure 13:

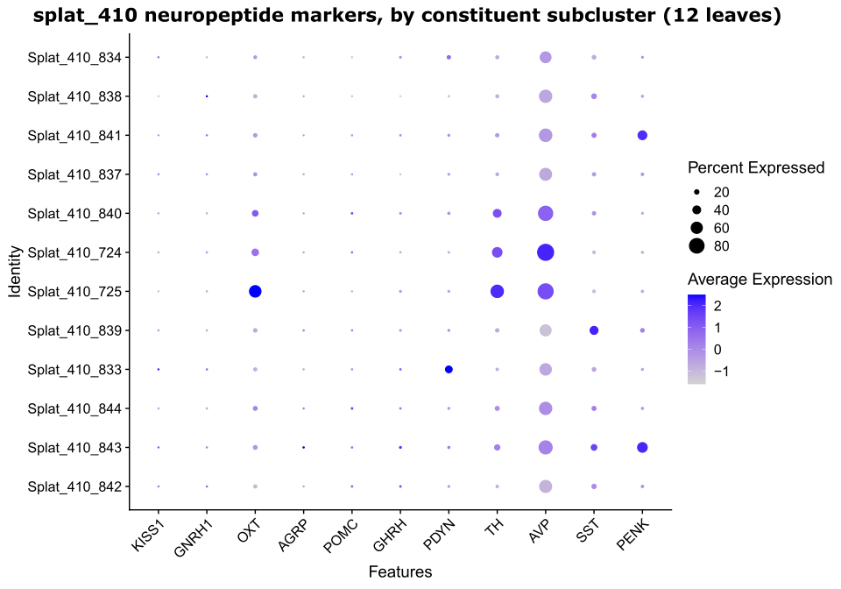

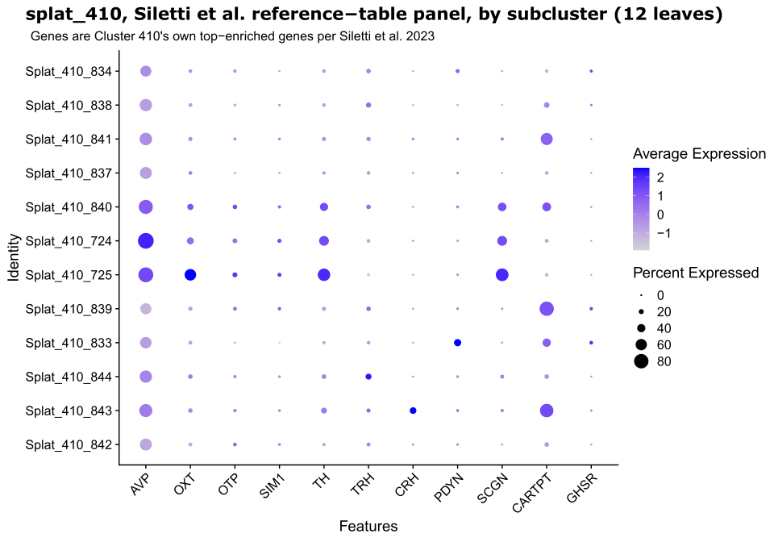

**Neuropeptides and atlas-reference marker expression across Splat_410 subclusters.**

(A) Presents a dot plot showing canonical hypothalamic neuropeptide markers—such as KISS1, GNRH1, OXT, AGRP, POMC, GHRH, PDYN, TH, AVP, SST, and PENK—across the 12 subclusters of Splat_410. (B) Displays a dot plot of the top-enriched genes associated with Cluster 410, as reported in the Allen Brain Cell Atlas reference table (including AVP, OXT, OTP, SIM1, TH, TRH, CRH, PDYN, SCGN, CARTPT, GHSR; Siletti et al., 2023), measured across the same 12 subclusters. In both plots, dot size reflects the percentage of cells expressing the gene within each subcluster, while color indicates the average scaled expression. The subclusters are arranged in the same row order in both panels.

Supplementary Figure 14:

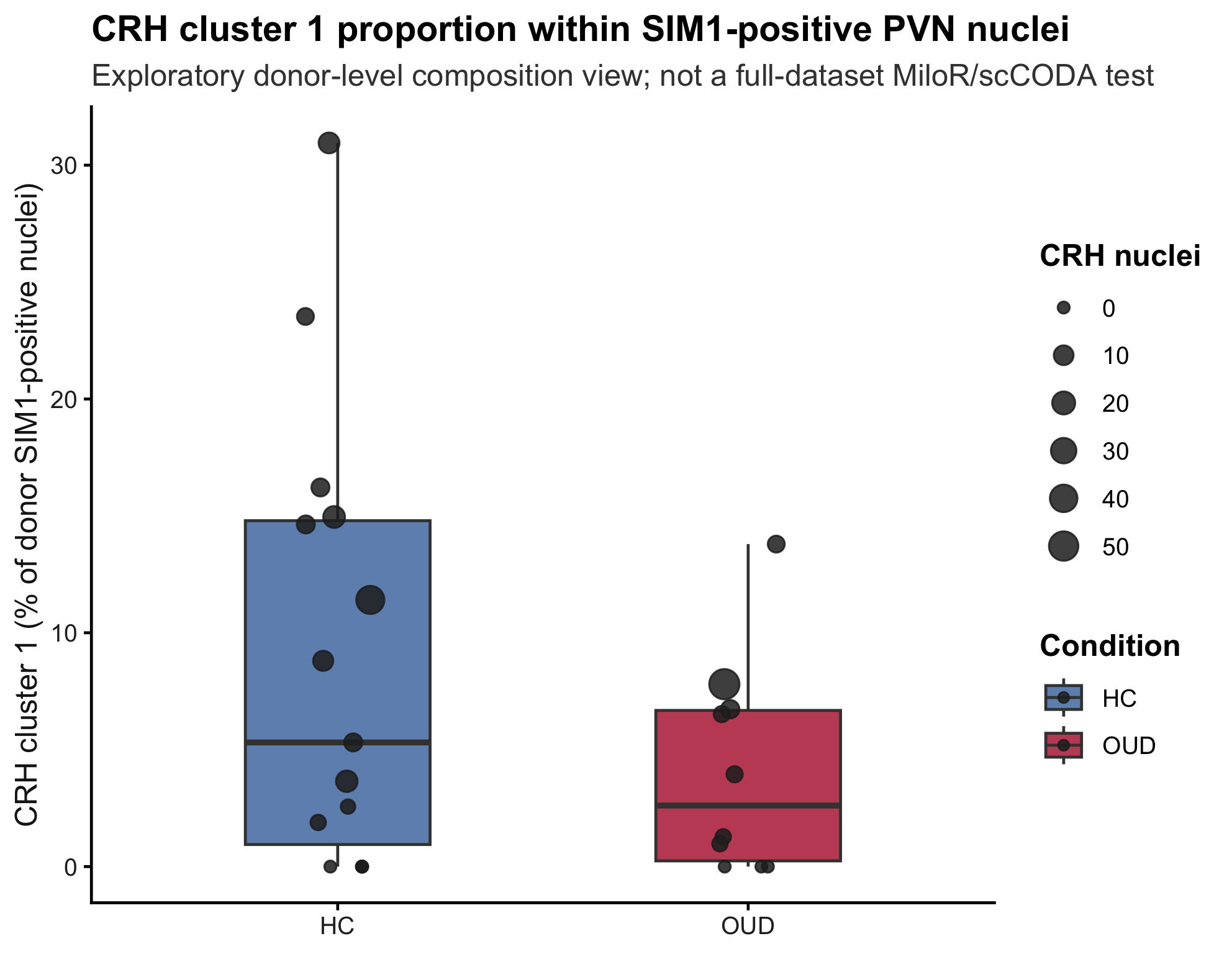

**Exploratory composition analysis of the CRH-enriched Crh-Nr3c1 population within the SIM1-positive PVN subset.**

Points represent donor-level proportions of SIM1-positive PVN nuclei assigned to Harmony cluster 1, stratified by metadata-derived HC/OUD condition from the Opioid column. Median CRH cluster 1 proportion was 5.31% in HC and 2.61% in OUD. A condition-only donor-level quasibinomial model did not identify a significant OUD-HC difference for this cluster (P = 0.204; Benjamini-Hochberg adjusted P = 0.454), and a Wilcoxon test was similarly non-significant (P = 0.274). This analysis is exploratory and does not replace full-dataset differential abundance testing such as MiloR or scCODA.

Supplementary Figure 15:

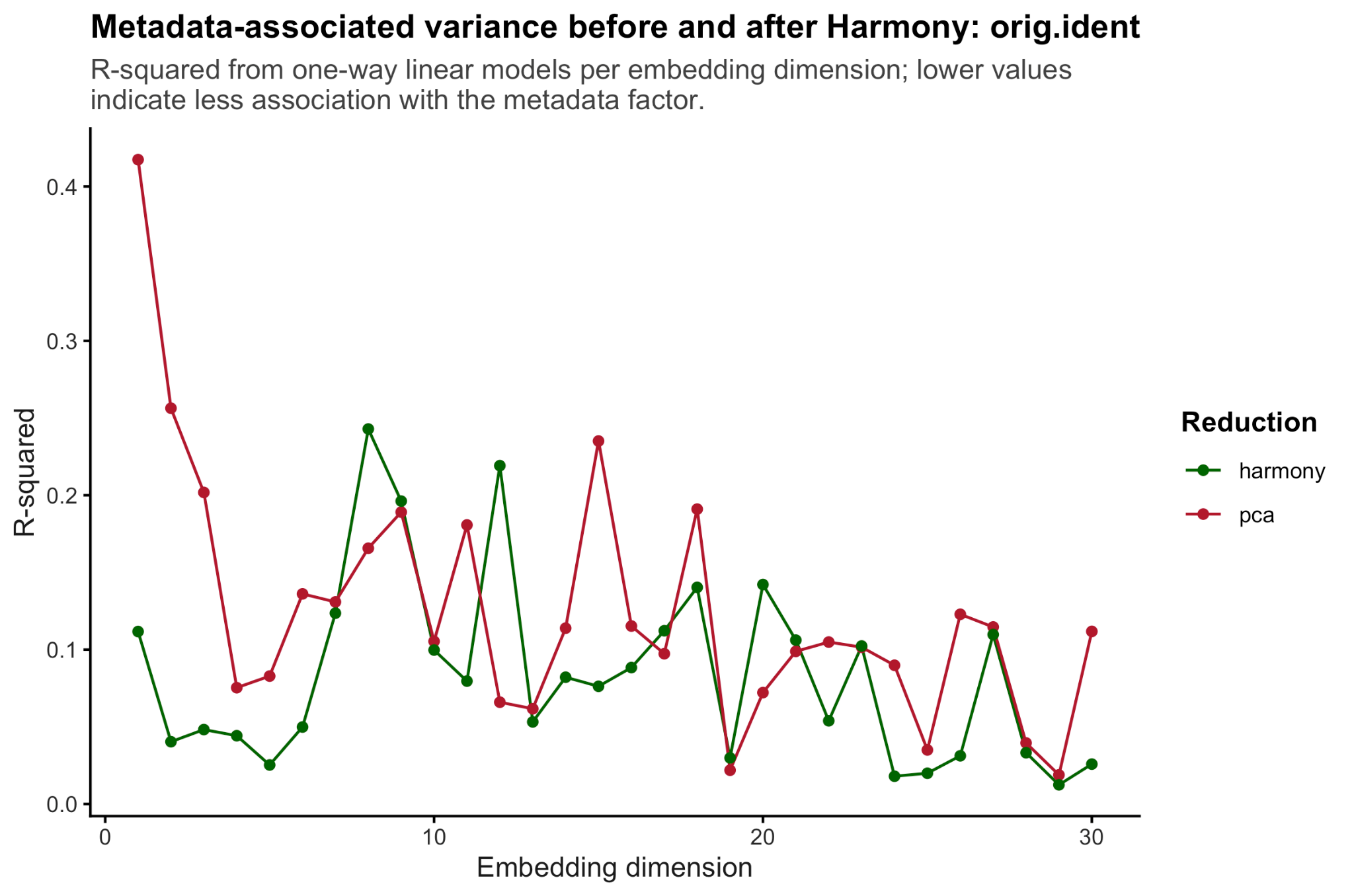

**Metadata-associated variance before and after Harmony correction in the available SIM1-positive PVN subset.**

Linear-model R-squared values were calculated for metadata covariates across PCA and Harmony dimensions. Variance-weighted donor/sample-associated R-squared decreased from 0.149 in PCA space to 0.099 in Harmony space, consistent with reduced donor/sample structure after Harmony correction. Diagnosis-associated R-squared remained low in both PCA and Harmony space.

Supplementary Figure 16:

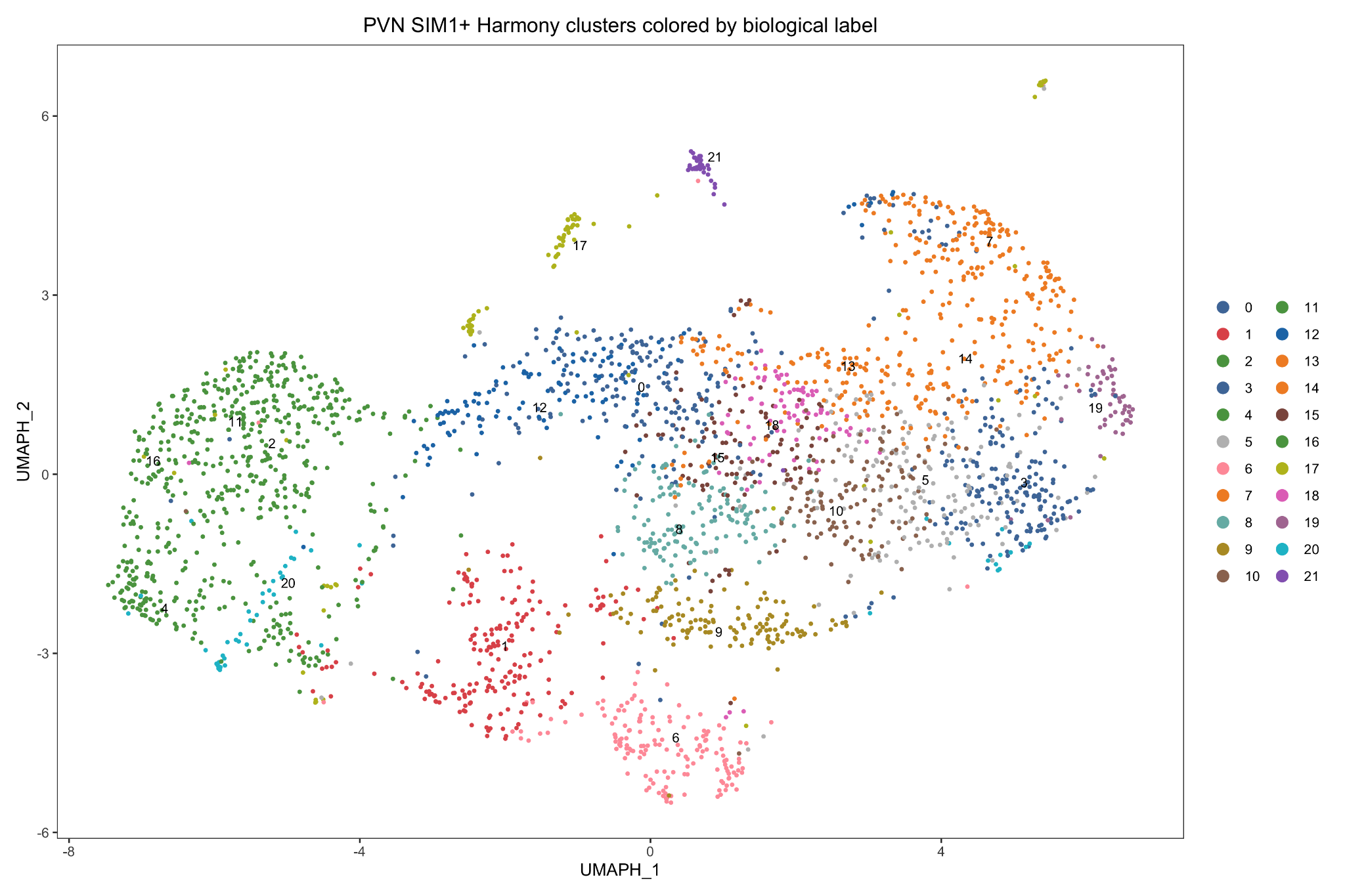

**UMAP visualization of unsupervised Harmony clusters colored by corresponding biological label.**

UMAP projection of the same 2,847 SIM1-positive PVN neurons, colored by the original unsupervised Harmony cluster assignments generated from the Harmony SNN graph at resolution 3.5. Harmony clusters are numbered 0-21. Colors correspond to the post hoc biological label assigned to each Harmony cluster, using the same palette as Figure 1A. Harmony clusters sharing the same biological annotation are shown in the same color, highlighting cases where multiple transcriptionally distinct Harmony clusters map to the same marker-defined PVN Atlas-associated identity. Harmony cluster 5, assigned a low-confidence Asb4-Adarb2-like label and excluded from downstream label-specific analyses, is shown in gray.

Supplementary Figure 17:

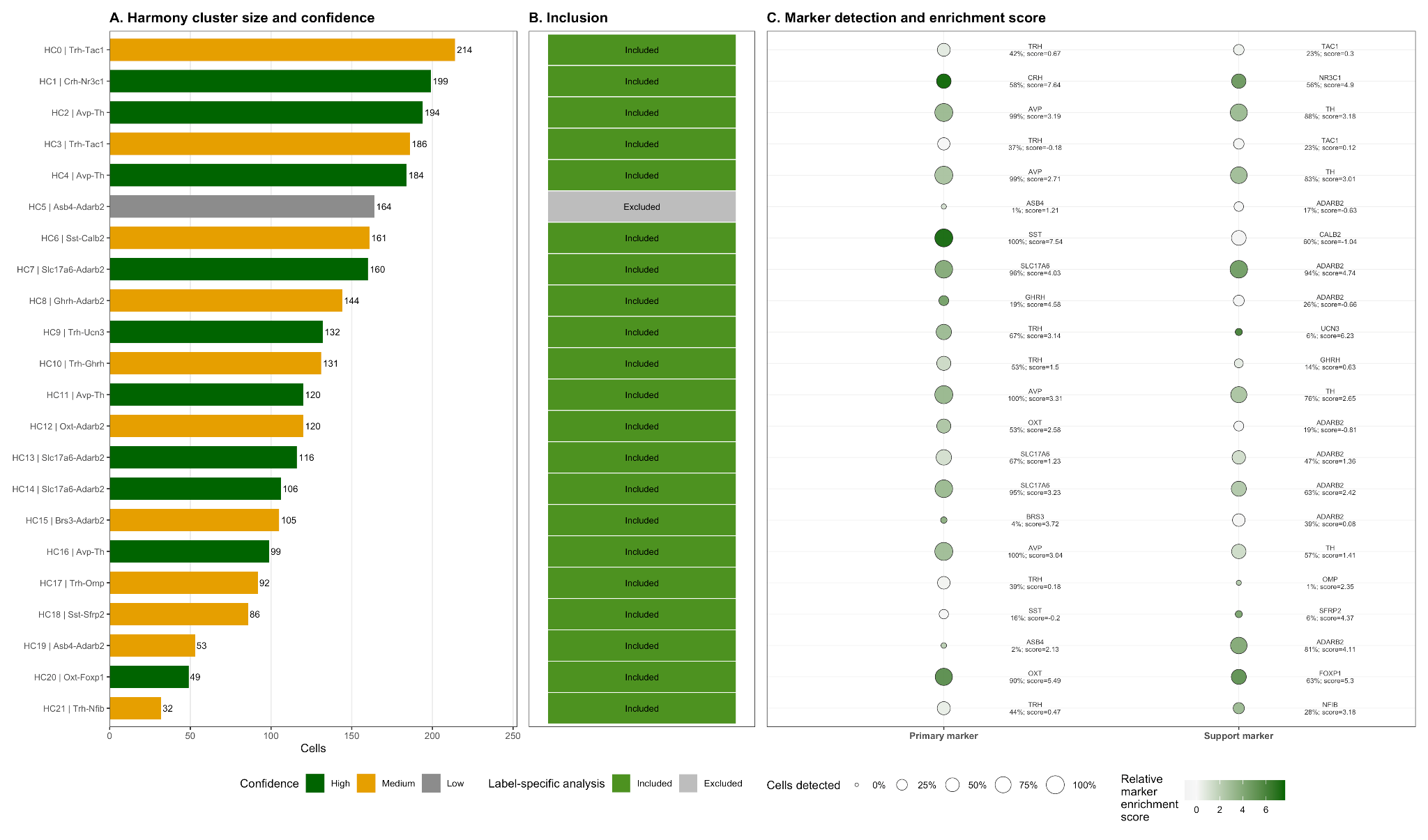

**Marker detection and enrichment scores underlying post hoc PVN biological-label confidence assignments.**

Bar plot showing the number of SIM1-positive PVN neurons in each unsupervised Harmony cluster, colored by post hoc biological-label confidence tier. High-confidence clusters are shown in dark green, medium-confidence clusters in orange, and the low-confidence cluster in gray. (B) Inclusion status for downstream label-specific analyses. High- and medium-confidence clusters were retained for downstream biologically labelled analyses, whereas the single low-confidence cluster, Harmony cluster 5, was excluded from label-specific analyses. (C) Marker evidence supporting each assigned biological label. For each Harmony cluster, the primary marker and label-refining support marker are shown. Dot size represents the percentage of cells within the cluster with detectable marker expression. Green intensity represents the relative marker enrichment score, calculated as gene-wise z-scored average normalized expression plus 0.75 times gene-wise z-scored detection frequency. Numeric labels indicate percent detected and relative marker enrichment score for each marker. Together, these panels show that confidence assignments were based on cluster size, marker detection frequency, and relative marker enrichment rather than forced assignment to PVN Atlas cell types.

Supplementary Figure 18:

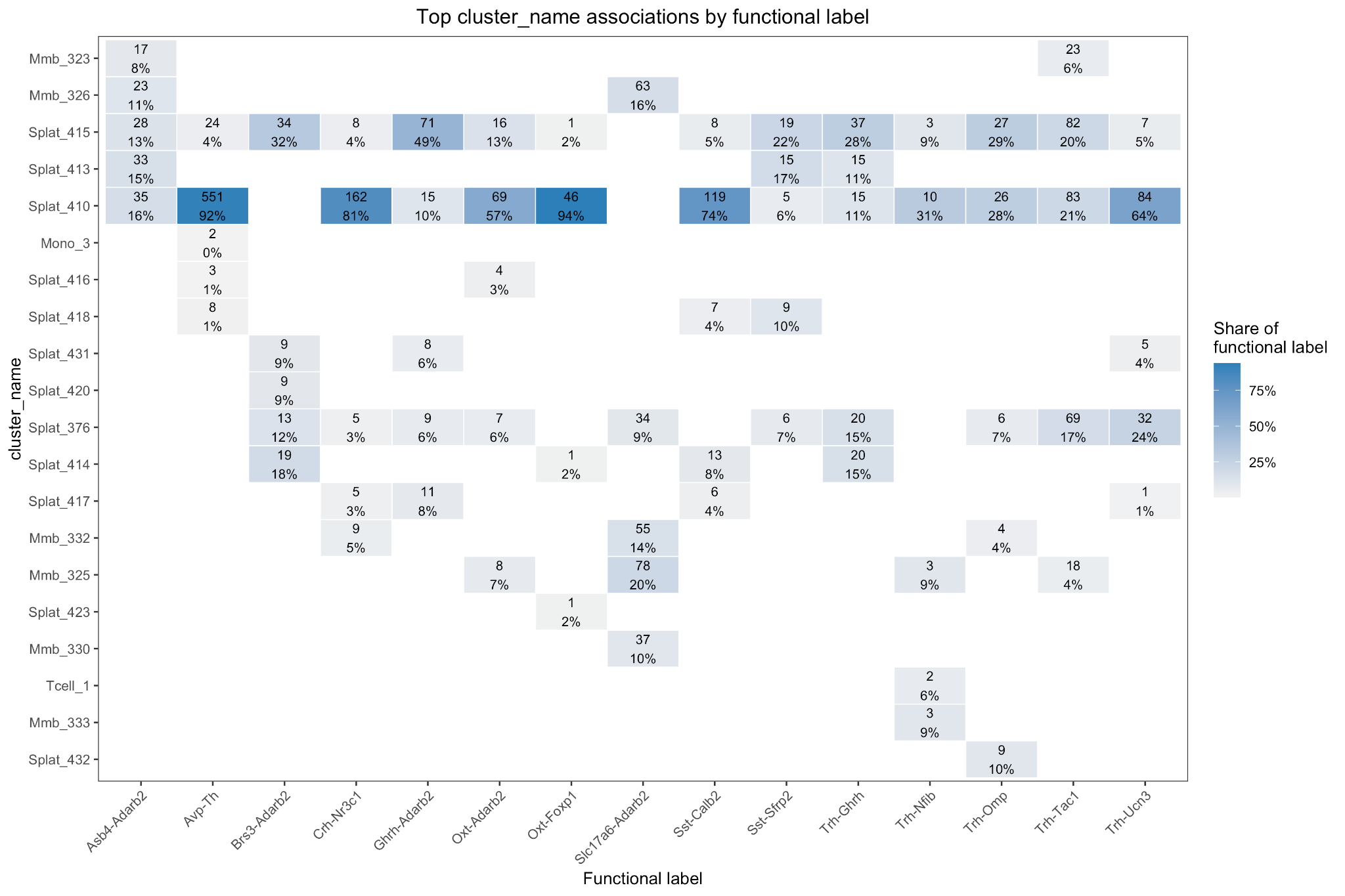

**Correspondence between Harmony-derived PVN clusters and broad Allen MapMyCells cluster assignments.**

Heatmap showing the top broad Allen MapMyCells cluster_name associations for each Harmony-derived functional cluster. Columns represent the 22 unsupervised Harmony clusters, labeled by Harmony cluster ID and post hoc biological marker annotation. Rows represent prior Allen MapMyCells broad cluster assignments. Tile color indicates the fraction of cells within each Harmony cluster assigned to the corresponding Allen MapMyCells cluster_name; overlaid text indicates the number of overlapping cells and the percentage of the Harmony cluster represented by that overlap. This analysis was used as a post hoc validation step to compare unsupervised Harmony cluster structure with prior Allen MapMyCells annotations. Harmony cluster 1, assigned a high-confidence Crh-Nr3c1 label, showed predominant overlap with the broad Splat_410 lineage.

Supplementary Figure 19:

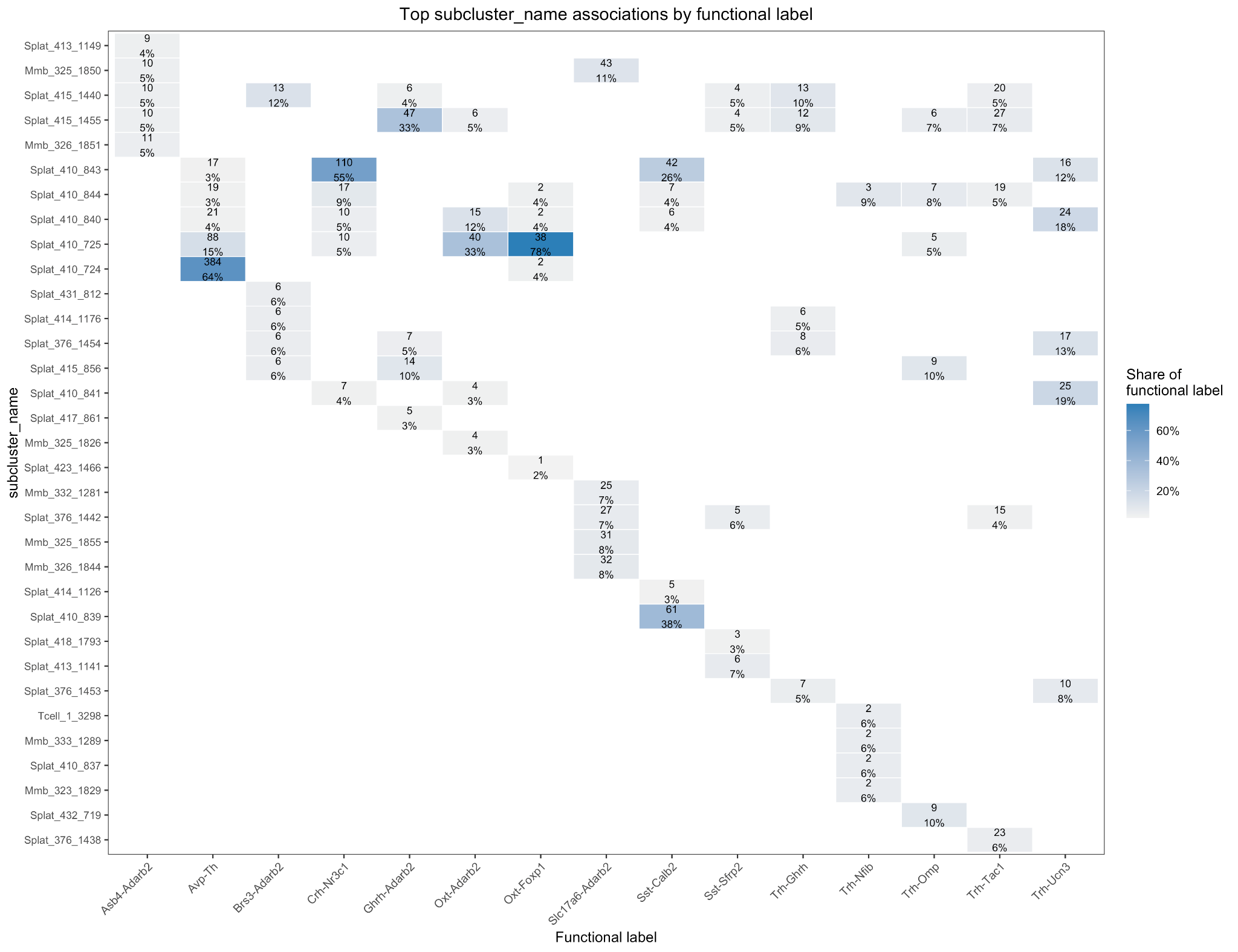

**Correspondence between Harmony-derived PVN clusters and fine Allen MapMyCells subcluster assignments.**

Heatmap showing the top Allen MapMyCells subcluster_name associations for each Harmony-derived functional cluster. Columns represent the 22 unsupervised Harmony clusters, labeled by Harmony cluster ID and post hoc biological marker annotation. Rows represent prior Allen MapMyCells subcluster assignments. Tile color indicates the fraction of cells within each Harmony cluster assigned to the corresponding Allen MapMyCells subcluster_name; overlaid text indicates the number of overlapping cells and the percentage of the Harmony cluster represented by that overlap. This analysis was used as a post hoc validation step to determine whether Harmony-derived PVN clusters localized to specific Allen MapMyCells fine subclusters. Harmony cluster 1, assigned a high-confidence Crh-Nr3c1 label, showed its strongest fine-level overlap with Splat_410_843, supporting localization of the CRH-enriched, ADARB2-detectable SIM1-positive PVN population to this Allen MapMyCells subcluster.

Supplementary Figure 20:

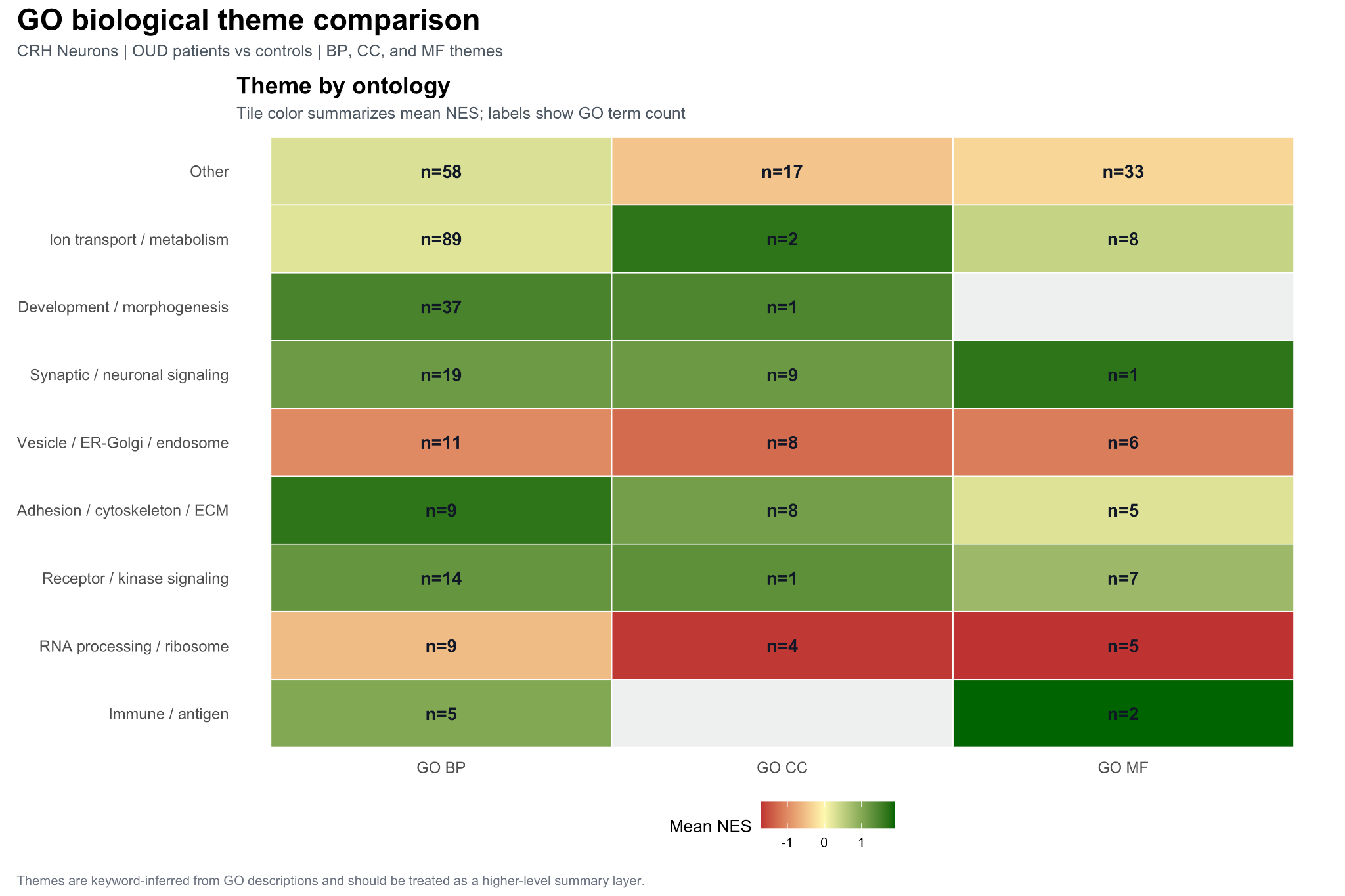

**GO GSEA analysis of Pathways Across**

Biological-theme summary of GO GSEA results in CRH neurons from OUD patients versus controls. GO terms passing q ≤ 0.25 or nominal p < 0.05 were assigned to keyword-derived biological themes based on GO descriptions. Tile color represents the mean NES across terms within each theme and ontology, with green indicating enrichment in OUD and red indicating enrichment in controls. Labels indicate the number of GO terms assigned to each theme. Overall, OUD-associated positive enrichment is strongest across developmental/ morphogenesis, synaptic/ neuronal signaling, receptor/ kinase signaling, adhesion/ cytoskeleton/ ECM, and immune/ antigen themes. In contrast, vesicle/ ER-Golgi/ endosome and RNA processing/ ribosome themes show negative mean NES, indicating relatively stronger enrichment in controls.

Supplementary Figure 21:

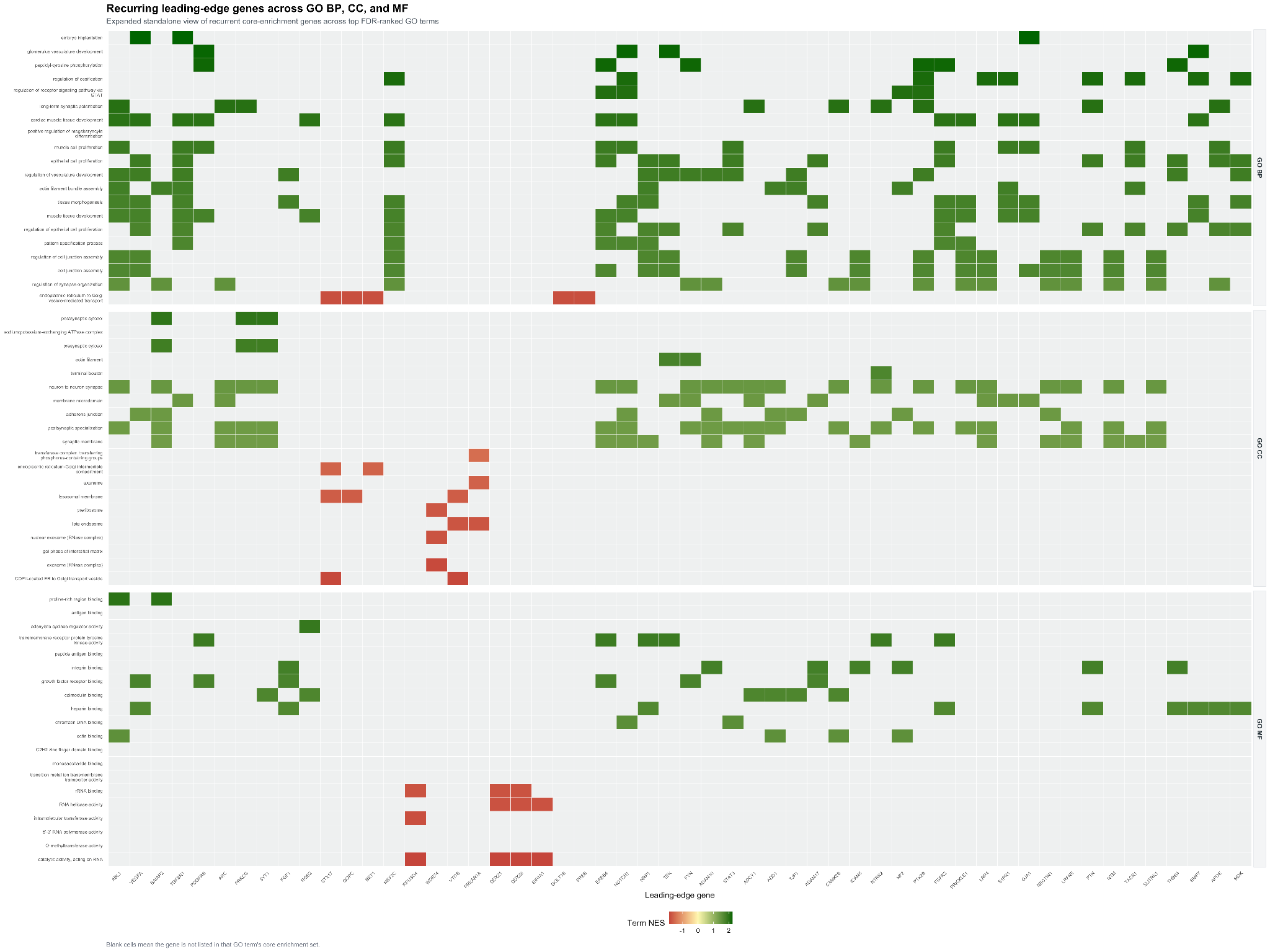

**Heatmap showing recurrent leading-edge/core-enrichment genes across enriched Gene Ontology terms in CRH neurons from OUD patients compared with controls.**

GO Biological Process, Cellular Component, and Molecular Function terms were ranked within each ontology by qvalue, then pvalue, then absolute normalized enrichment score, abs(NES). The top 20 terms per ontology were included, yielding 60 GO terms total. Leading-edge genes were extracted from the core_enrichmentfield of the GSEA output and mapped from Entrez identifiers to gene symbols. Genes were then prioritized for display by recurrence across GO terms, span across GO ontologies, span across enrichment direction, minimum gene-level p value, and strongest absolute log2 fold-change, with up to 54 recurrent genes shown.

Supplementary Figure 22:

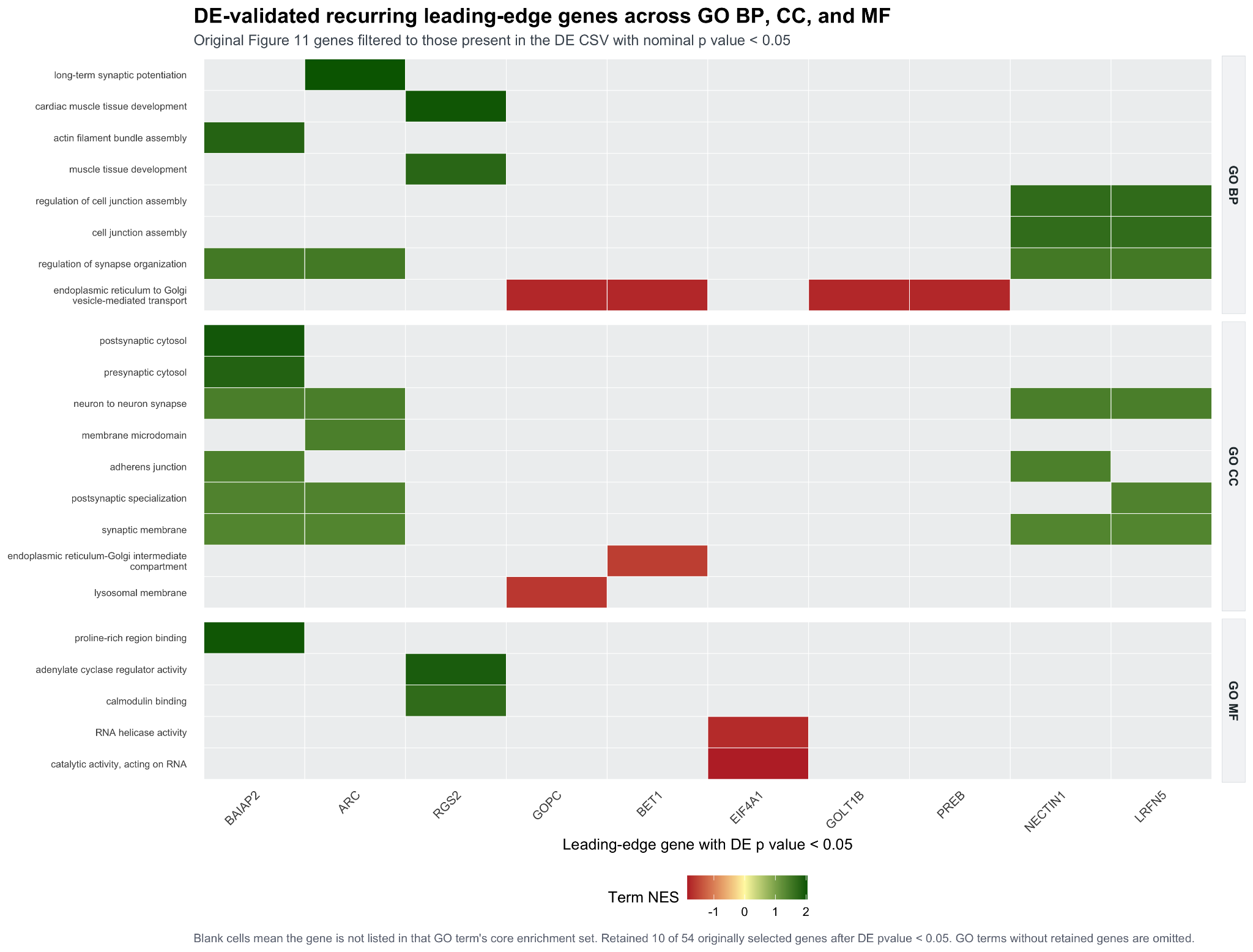

**Heatmap showing recurrent GO leading-edge/core-enrichment genes in CRH neurons from OUD patients compared with controls.**

GO Biological Process, Cellular Component, and Molecular Function terms were ranked within ontology by qvalue, then pvalue, then absolute normalized enrichment score, abs(NES), and the top 20 terms per ontology were considered. Recurrent leading-edge genes from the original Figure 11a selection were cross-validated against the CRH pseudobulk differential expression table, retaining only genes present in the DE CSV with nominal pvalue < 0.05. This filter retained 10 of 54 originally selected genes: BAIAP2, ARC, RGS2, GOPC, BET1, EIF4A1, GOLT1B, PREB, NECTIN1, and LRFN5.

Supplementary Figure 23:

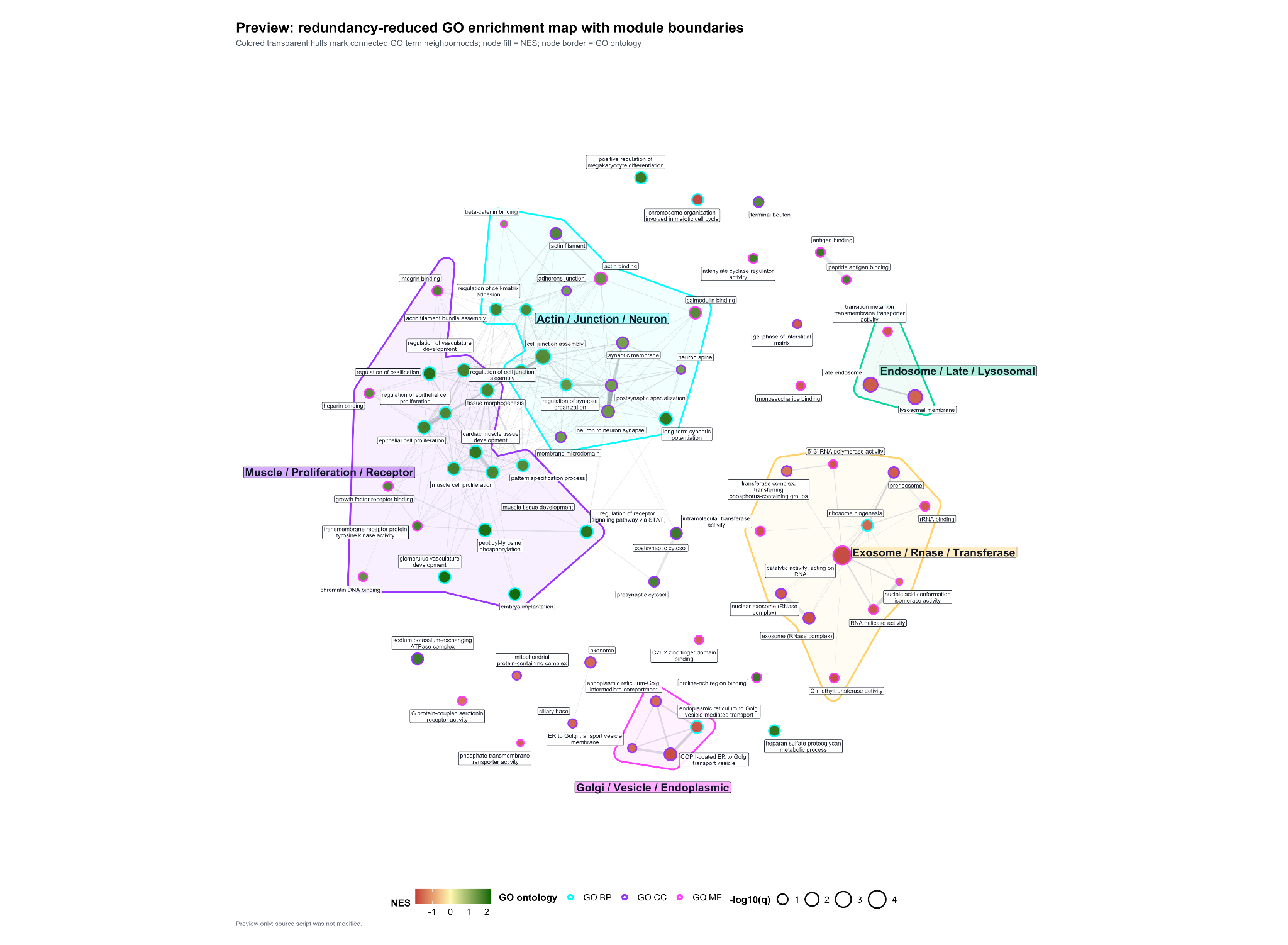

**Enrichment map of GO BP, CC, MF by semantic reduction and leading-edge/core-enrichment genes**

This enrichment map shows representative GO Biological Process, Cellular Component, and Molecular Function terms after semantic redundancy reduction. Each node is one GO term; edges connect terms sharing leading-edge/core-enrichment genes, so clustered nodes represent biologically related GO programs.

The map highlights five main functional neighborhoods. OUD-enriched modules are dominated by Muscle / Proliferation / Receptor and Actin / Junction / Neuron themes, suggesting coordinated shifts in morphogenesis, receptor/phosphorylation, adhesion, cytoskeletal, junctional, and synaptic/neuronal processes. Control-enriched modules include Exosome / RNase / Transferase, Golgi / Vesicle / Endoplasmic, and Endosome / Late / Lysosomal, indicating reduced or oppositely directed enrichment of RNA-processing, vesicular trafficking, endosomal/lysosomal, and ER-Golgi-associated terms in OUD CRH neurons.

For this figure, GO terms were first ranked within each ontology by statistical evidence and effect strength, prioritizing lower qvalue, then lower pvalue, then larger absolute normalized enrichment score, abs(NES). To reduce redundancy among closely related GO terms, semantic similarity was calculated using the Wang method, and terms with similarity >= 0.70 were grouped; the highest-ranked term within each redundancy group was retained as the representative term. From these redundancy-reduced terms, the top 24 GO BP, 24 GO CC, and 24 GO MF terms were selected for visualization.

The enrichment map was then constructed as a term-overlap network. Each GO term was treated as a node, and pairs of GO terms were connected when they shared at least one leading-edge/core-enrichment gene. Edge strength was quantified using Jaccard overlap, defined as the number of shared leading-edge genes divided by the total number of unique leading-edge genes across the two terms. Edges were retained if Jaccard >= 0.025, with a maximum of 240 edges shown.

Network topology was visualized using a Fruchterman-Reingold force-directed layout with niter = 4000 and seed = 149, so terms sharing more core genes were positioned closer together. Functional modules were detected using Louvain community detection weighted by Jaccard overlap. Transparent colored hulls were drawn around multi-term modules to emphasize clustered biological neighborhoods, while bold module labels summarize recurring words across the GO terms in each cluster.

Node aesthetics encode the main biological interpretation: node fill represents NES, with red indicating enrichment higher in controls, yellow near zero, and green enrichment higher in OUD; node border color identifies ontology, with GO BP in cyan, GO CC in purple, and GO MF in magenta. Node size represents -log10(q), so larger nodes correspond to stronger enrichment evidence.

Supplementary Figure 24:

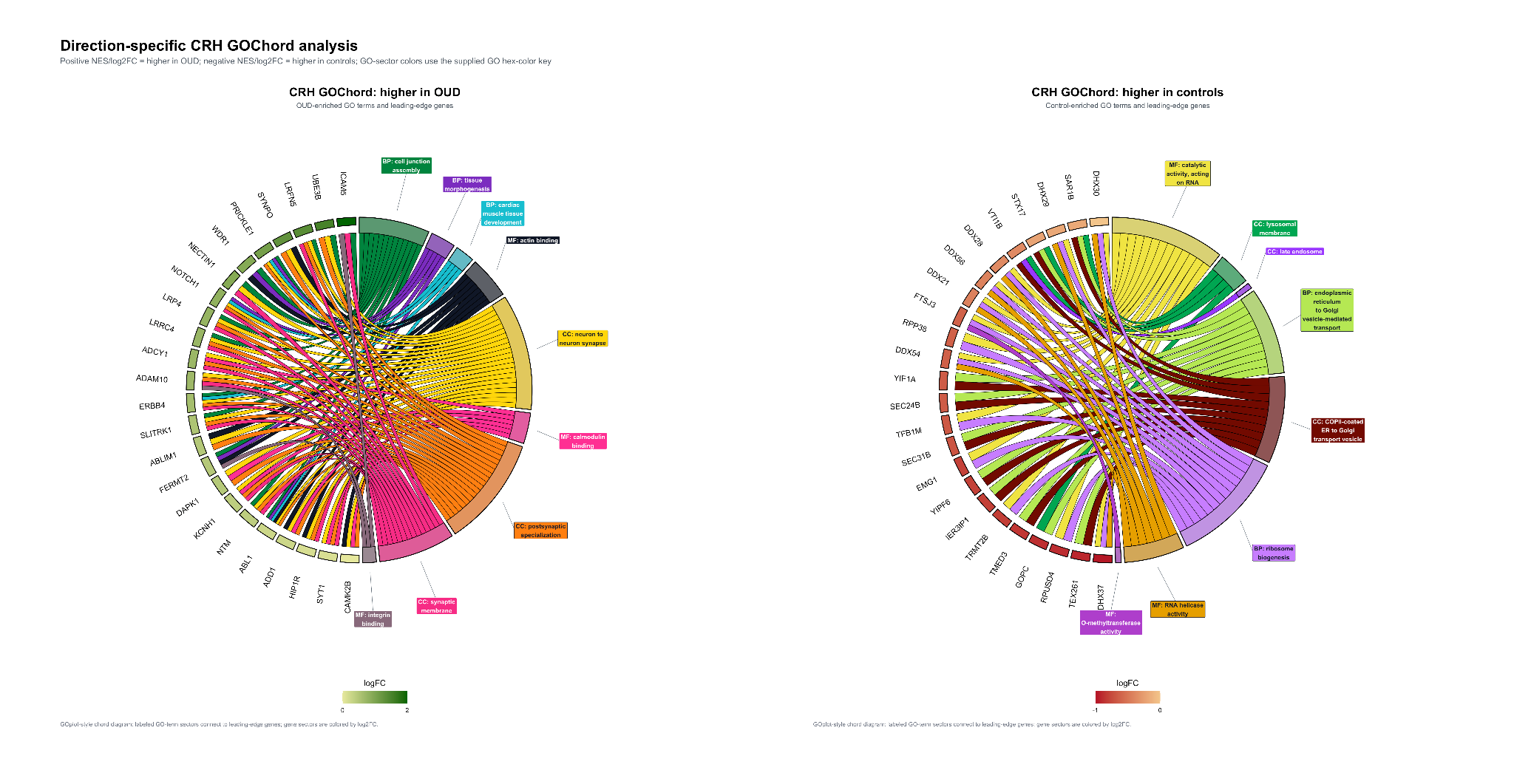

**Direction-specific GOChord analysis of enriched GO terms in CRH neurons.**

All GOChord figures used CRH neuron single-nucleus RNA-seq GSEA outputs for GO BP, GO CC, and GO MF from the OUD patients vs controls Wald-ranked contrast. Positive NES/log2FC denotes higher signal in OUD; negative NES/log2FC denotes higher signal in controls. Leading-edge/core genes were parsed from the GSEA core-enrichment field and linked to selected GO terms using GOplot::circle_dat, GOplot::chord_dat, and GOplot::GOChord. Genes were ranked by recurrence across selected GO terms: selected_term_count = n distinct GO terms, selected_ontology_count = n distinct GO ontologies, logFC = first finite log2FoldChange with shrunk log2FoldChange fallback, max_abs_logFC = max(abs(log2FC), abs(shrunk log2FC)), and best_qvalue = min(qvalue). Genes with selected_term_count >= 2 were retained when at least 8 recurrent genes were available; the top 24 genes were then selected by descending term count, descending ontology count, descending absolute logFC, ascending q-value, and gene symbol.
GOChord diagrams show direction-specific GO enrichment in CRH neurons from the OUD patients vs controls single-nucleus RNA-seq contrast. The left panel includes GO terms with positive NES, corresponding to higher enrichment in OUD; the right panel includes GO terms with negative NES, corresponding to higher enrichment in controls. Terms were eligible if qvalue <= 0.25 or nominal pvalue < 0.05, then ranked within each ontology by qvalue, pvalue, and abs(NES). Up to 3 terms per ontology were selected. The OUD-enriched panel contains 9 GO terms, including 3 BP, 3 CC, and 3 MF terms, with NES range 1.368 to 2.010, q-value range 0.00169 to 0.11288, and p-value range 2.89e-7 to 0.00104. The control-enriched panel contains 8 GO terms, including 2 BP, 3 CC, and 3 MF terms, with NES range -1.866 to -1.583, q-value range 2.70e-5 to 0.16964, and p-value range 2.29e-8 to 0.00228. Each panel displays 24 leading-edge/core genes. GOChord parameters were space = 0.02, gene.order = "logFC", gene.space = 0.32, gene.size = 3.7, nlfc = 1, lfc.min = -2, lfc.max = 2, and unique GO-sector colors assigned by GO ID.

Supplementary Figure 25:

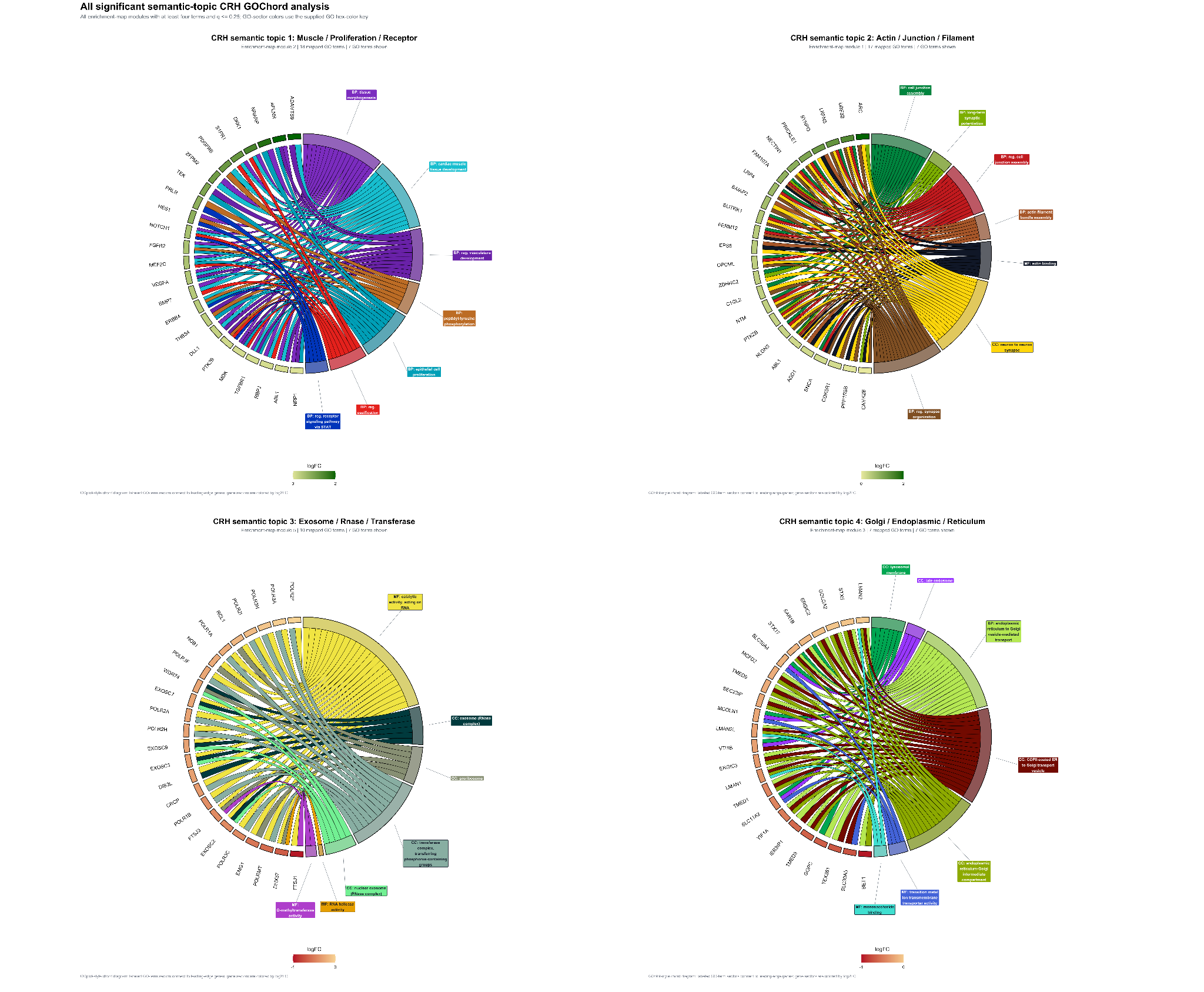

**Semantic-topic GOChord analysis of CRH neuron GO enrichment modules.**

GOChord diagrams show all significant semantic-topic modules identified from the CRH neuron GO enrichment-map network. The source enrichment map was generated from the top 20 FDR-ranked GO terms per ontology, with edges connecting terms by leading-edge/core gene overlap using shared >= 2 or Jaccard >= 0.04, with a maximum of 300 edges. Edge weights were defined as max(Jaccard, 0.001), modules were assigned by Louvain community detection, and node layout used Fruchterman-Reingold placement with seed 149 and niter = 4000. Semantic-topic modules were retained for GOChord plotting when term_count >= 4 and min_qvalue <= 0.25. Within each module, up to 7 GO terms were selected by qvalue and abs(NES), followed by ordering by qvalue, pvalue, and abs(NES). The combined figure includes 4 semantic topics, 28 displayed GO-term entries, and 24 leading-edge/core genes per panel. GOChord parameters were space = 0.02, gene.order = "logFC", gene.space = 0.34, gene.size = 3.55, nlfc = 1, lfc.min = -2, lfc.max = 2, and unique GO-sector colors assigned by GO ID.

Supplementary Figure 26:

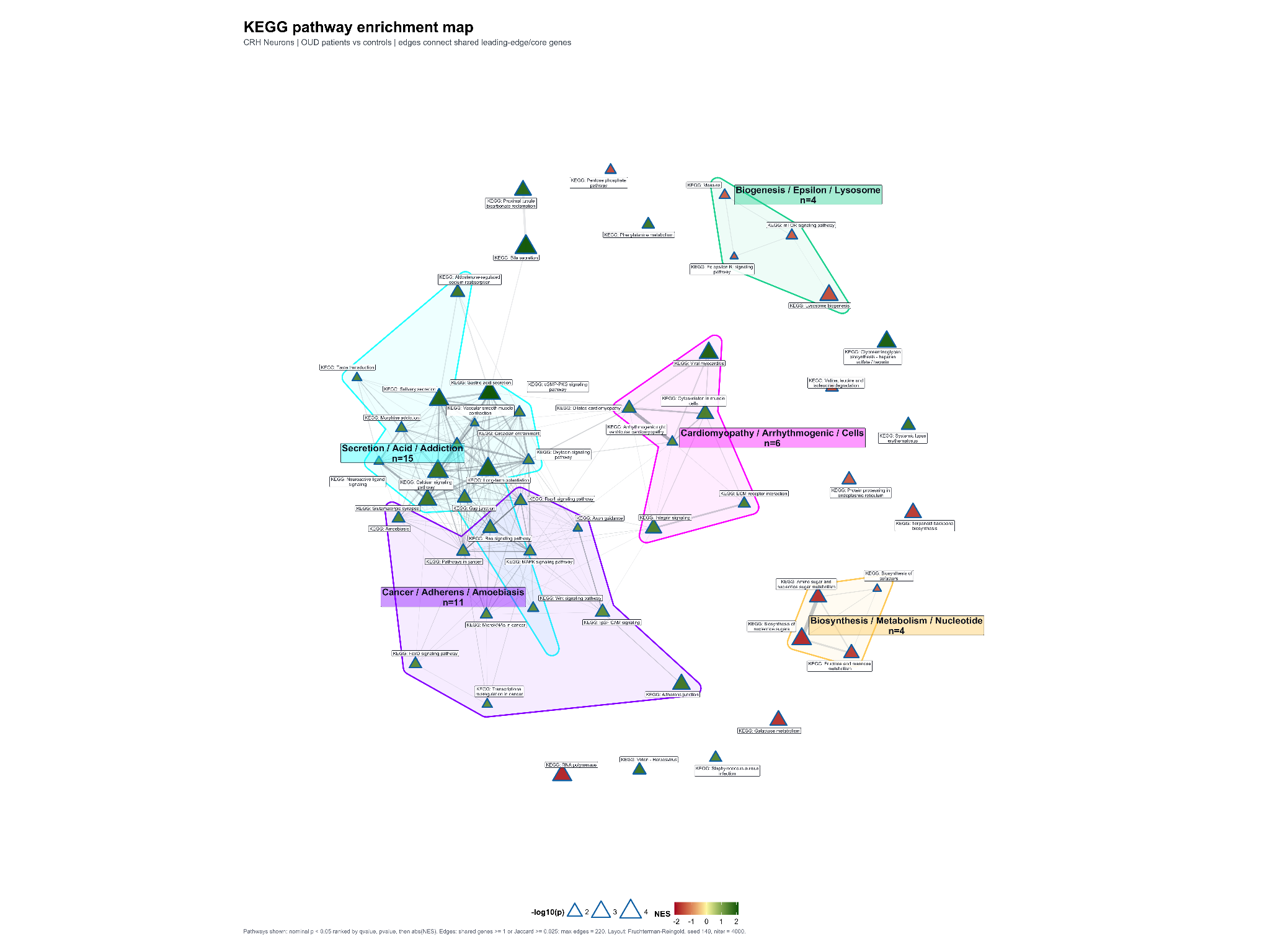

**KEGG pathway enrichment map in CRH neurons.**

Network map of nominally enriched KEGG pathways from the CRH neuron OUD patients versus controls GSEA. Nodes represent KEGG pathways, node fill indicates NES, node size indicates -log10(nominal p), and edges connect pathways sharing leading-edge/core genes. The map includes 53 KEGG pathways, 220 overlap edges, and 5 Louvain modules. Edge criteria were shared genes >= 1 or Jaccard overlap >= 0.025; layout used Fruchterman-Reingold, seed 149, niter = 4000. The module names were generated after clustering. For each Louvain module, the script collected the KEGG pathway descriptions in that module, split them into words, removed common stop words such as pathway, signaling, protein, regulation, and kegg, then selected the most frequent remaining words. Up to 3 frequent words were title-cased and joined with /. The labels are therefore keyword-derived summaries, not curated KEGG categories and not semantic ontology labels.

Supplementary Figure 27:

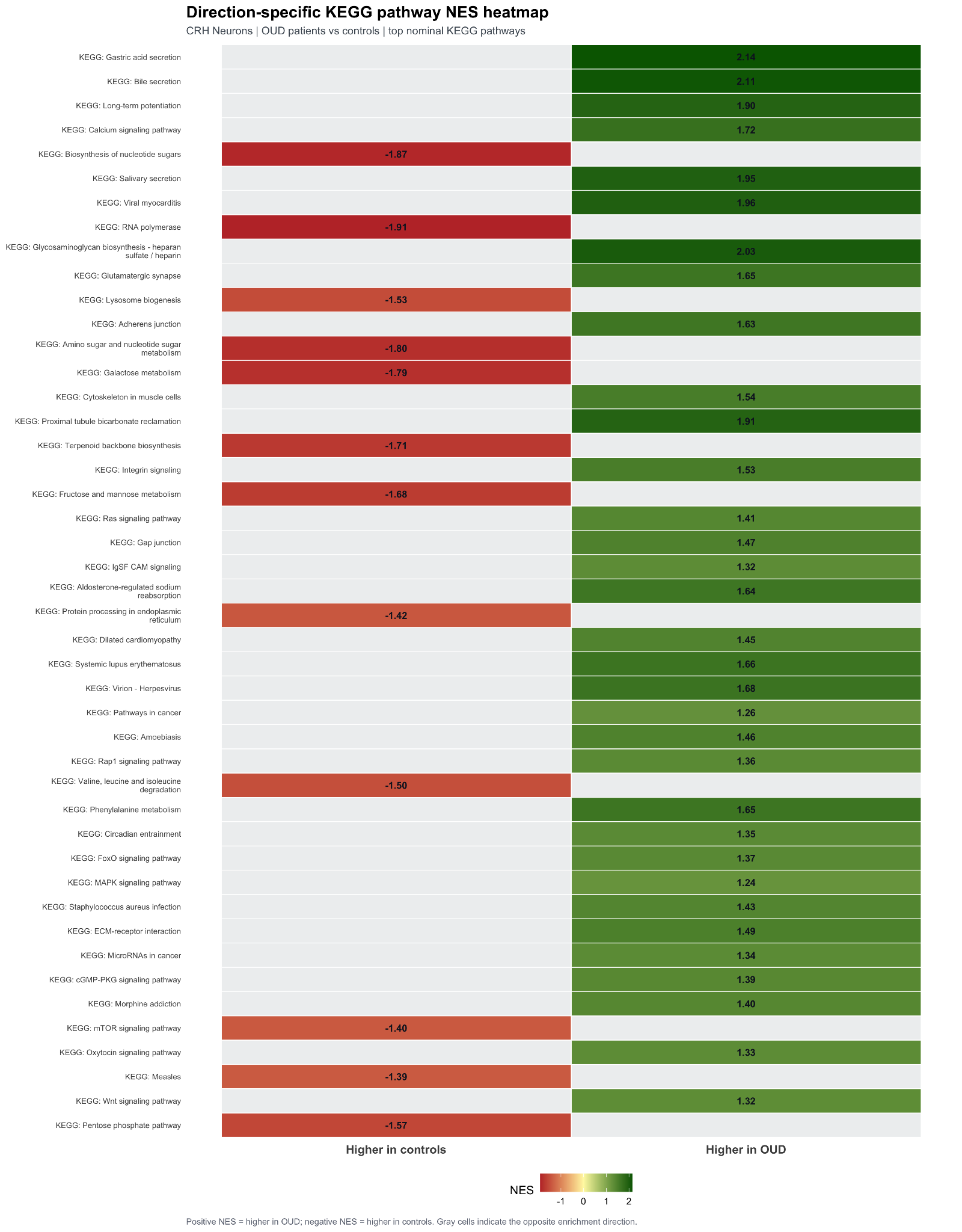

**Direction-specific KEGG pathway NES heatmap.**

Heatmap showing the top 45 KEGG pathways separated by enrichment direction. Green tiles indicate positive NES/higher enrichment in OUD, red tiles indicate negative NES/higher enrichment in controls, and tile labels report pathway NES rounded to two decimals. Pathways were ranked by qvalue, pvalue, then abs(NES).

Supplementary Figure 28:

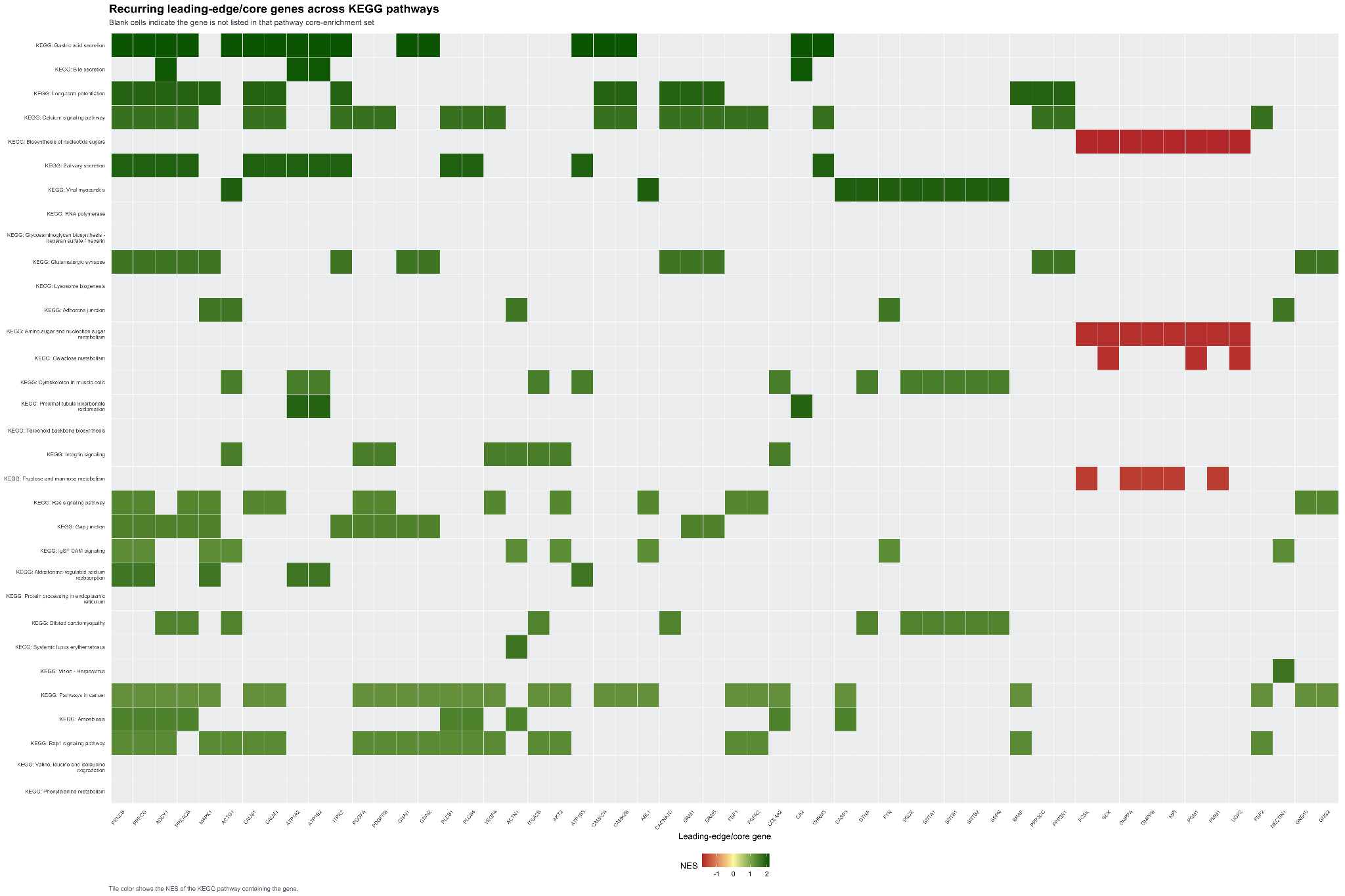

**Recurring KEGG leading-edge/core gene heatmap.**

Unfiltered gene-by-pathway heatmap showing recurring leading-edge/core genes across selected KEGG pathways. Rows show 32 KEGG pathways, columns show 56 recurrent leading-edge/core genes, and tile color reports the NES of the pathway containing that gene. Genes were prioritized by recurrence across pathways, maximum absolute pathway NES, pathway p value, and gene symbol.

Supplementary Figure 29:

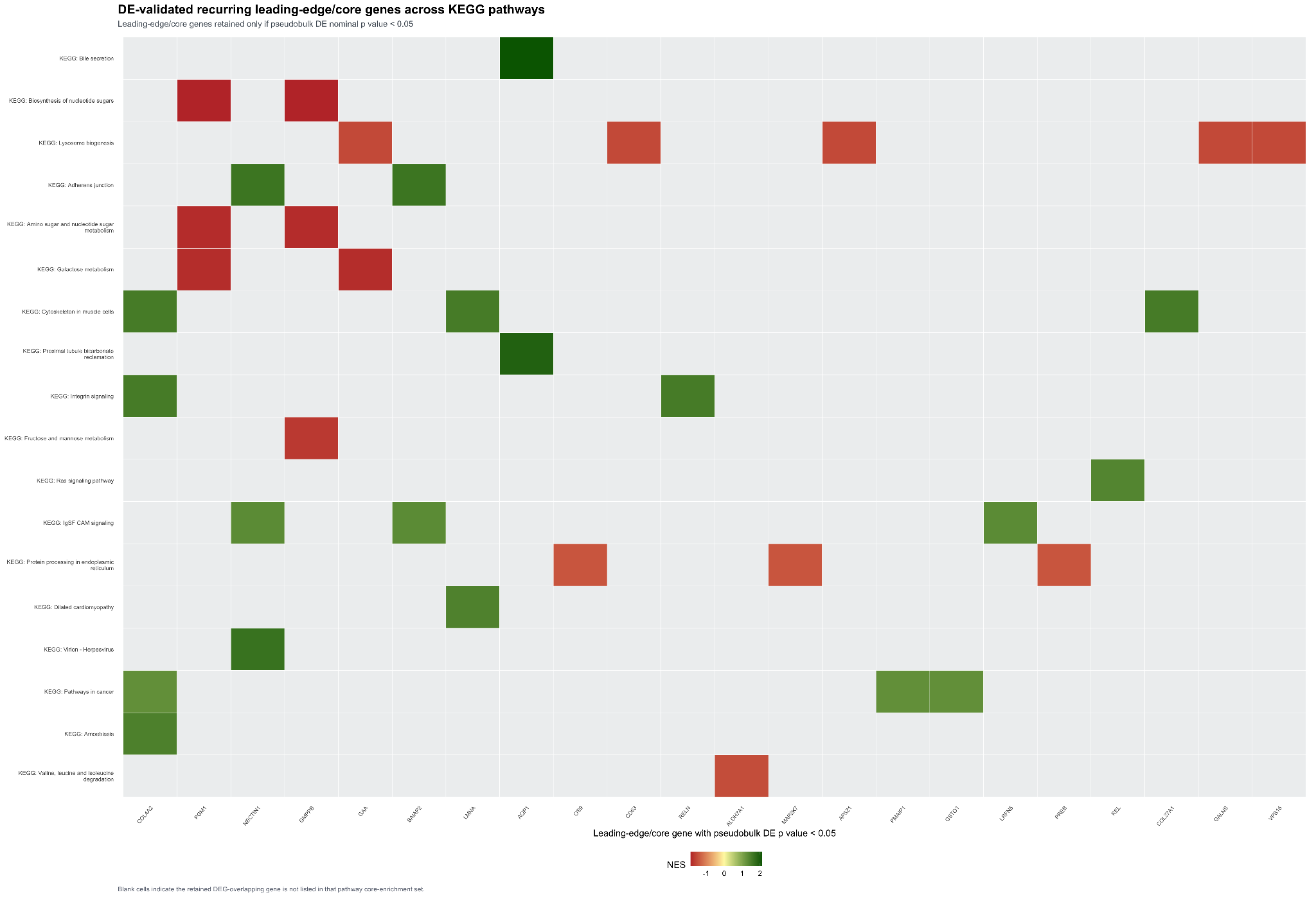

**DE-validated KEGG leading-edge/core gene heatmap.**

Companion heatmap retaining only KEGG leading-edge/core genes also present in the CRH pseudobulk differential expression table with nominal gene-level pvalue < 0.05. This filter retained 22 genes across 18 KEGG pathways.

Supplementary Figure 30:

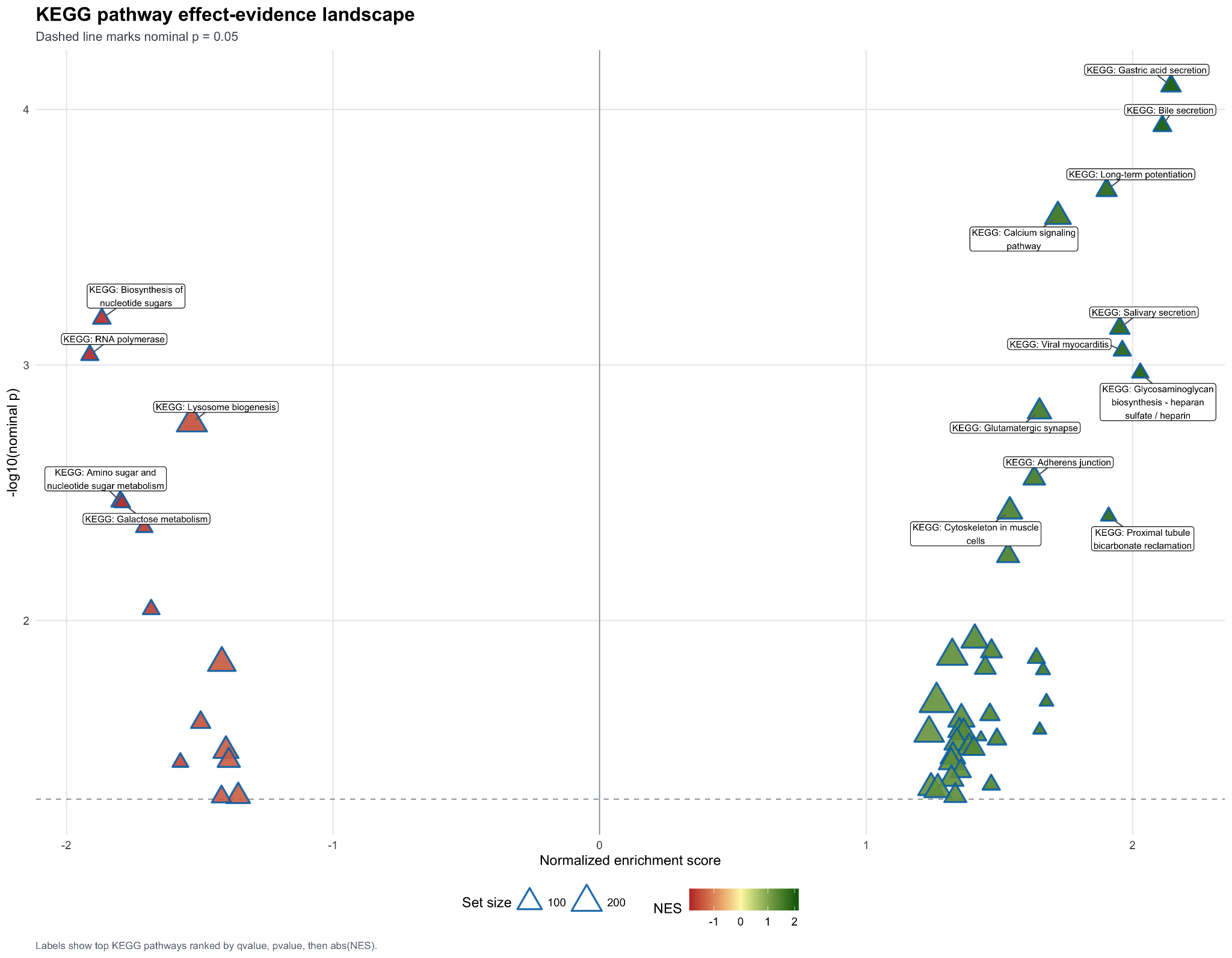

**KEGG pathway effect-evidence landscape.**

Scatterplot of KEGG pathway NES versus -log10(nominal p). Points represent 53 nominally enriched KEGG pathways, point size reflects KEGG set size, and labels mark the top 16 pathways ranked by qvalue, pvalue, and abs(NES). The dashed horizontal line marks nominal p = 0.05.

Supplementary Figure 31:

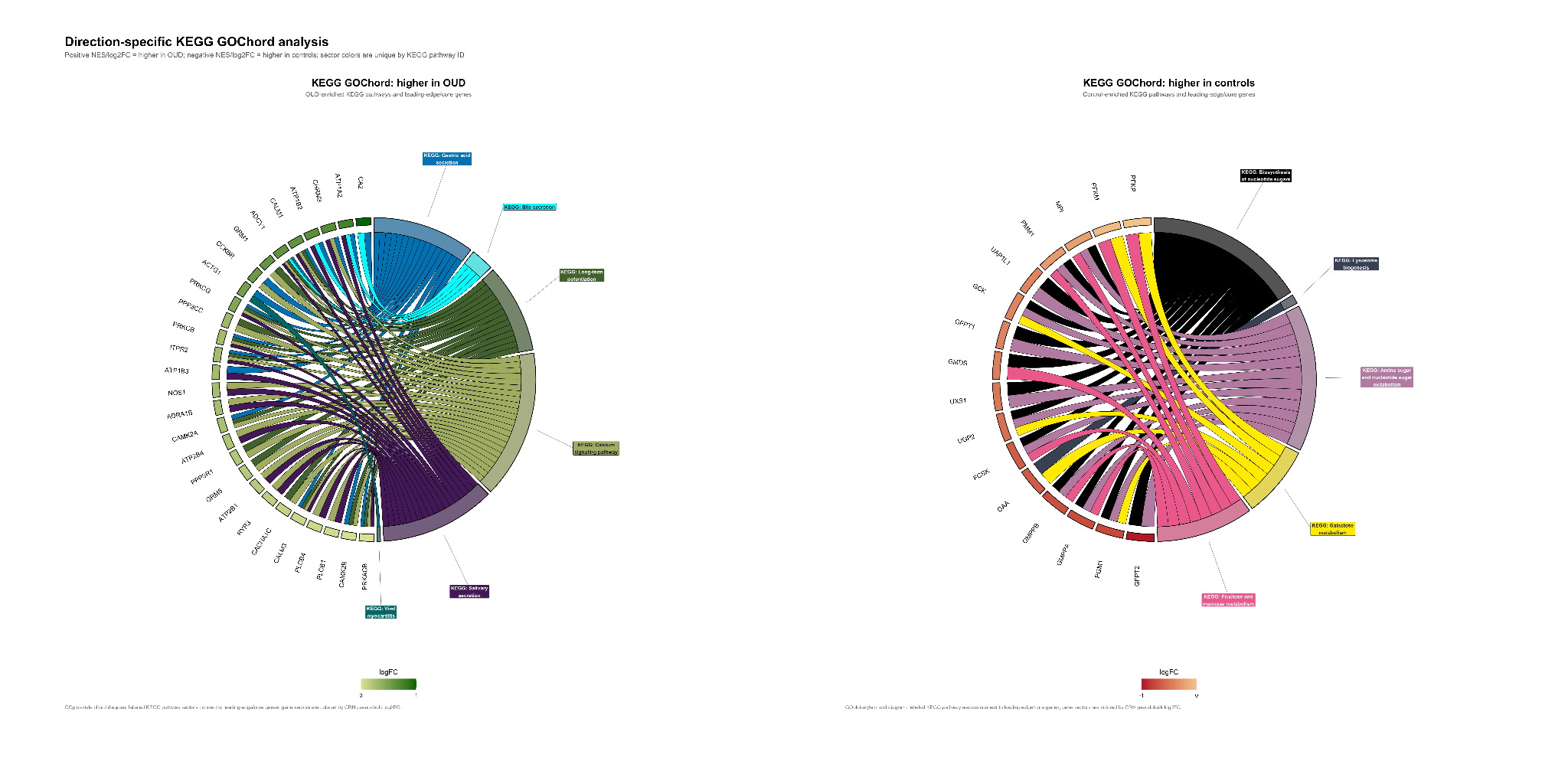

**Direction-specific KEGG GOChord analysis.**

GOChord-style diagrams connecting direction-specific KEGG pathways to their leading-edge/core genes. The OUD panel displays 6 OUD-enriched KEGG pathways and 28 genes; the control panel displays 5 control-enriched pathways and 16 genes after gene-overlap filtering. Pathway sectors use unique colors; gene sectors are ordered and colored by CRH pseudobulk log2FC.

Supplementary Figure 32:

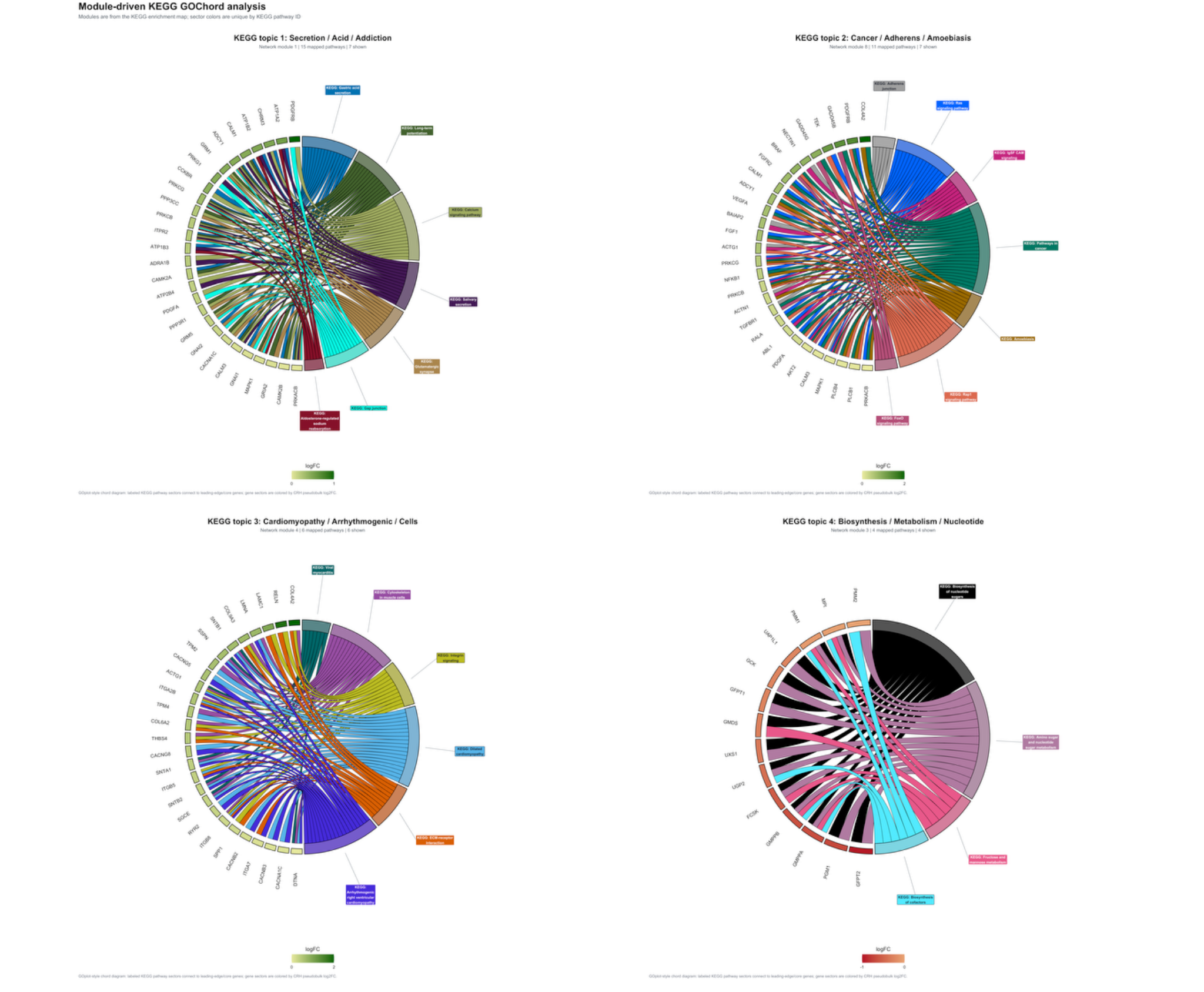

**Module-driven KEGG GOChord analysis.**

GOChord diagrams generated from KEGG enrichment-map Louvain modules. The combined figure shows the top 4 KEGG network modules; standalone files were also generated for all 5 modules. Displayed topics were Secretion/Acid/Addiction, Cancer/Adherens/Amoebiasis, Cardiomyopathy/Arrhythmogenic/Cells, and Biosynthesis/Metabolism/Nucleotide.

Supplementary Figure 33:

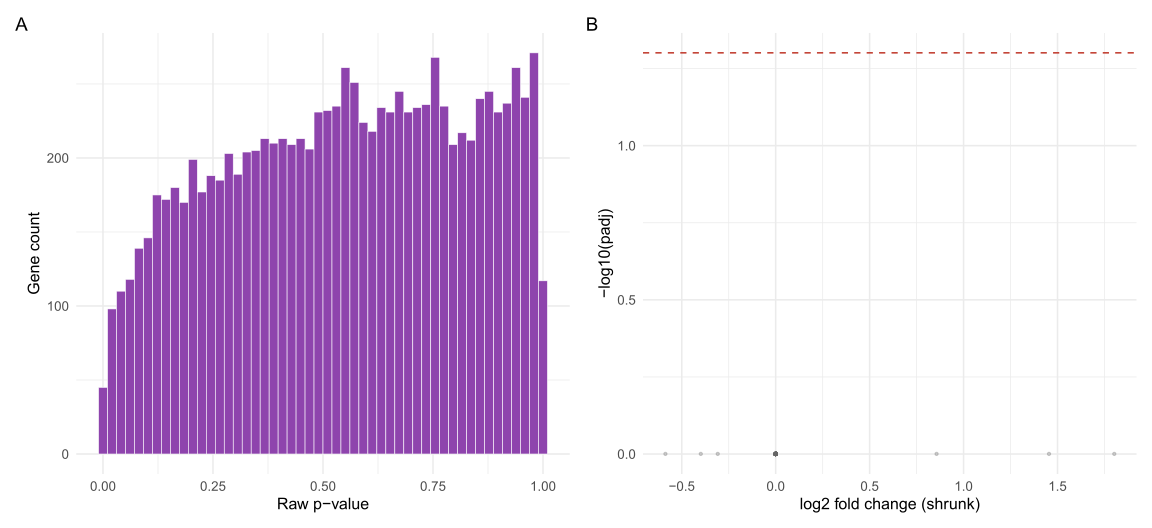

**Tier 3 (CRH+/PENK+ subcluster) differential expression: a clear null result in the targeted subpopulation.**

(A) Distribution of raw p-values across all 10,214 genes tested, comparing OUD to HC pseudobulk profiles within the CRH+/PENK+ subpopulation (Splat_410_841 and Splat_410_843; 25 donors, excluding donors with fewer than 10 contributing cells). The distribution appears uniform, indicating a true null rather than an underpowered non-finding. (B) Volcano plot of shrunken log3 fold change versus -log10(padj); the dashed line marks padj = 0.05. No genes reach significance (minimum padj = 0.999), despite tested effect sizes covering a genuine range (shrunk log2FC: -0.59 to 1.80)—a broader effect-size span than Tier 2, yet with a similarly flat significance profile, supporting that this null isn't due to limited effect-size variation. Combined with the Tier 2 result, no significant OUD-associated differential expression was observed at any of the three analytical scales examined in this study (general cell type, whole Splat_410, or the targeted CRH+/PENK+ subpopulation), despite this population’s independent, convergent validation as a genuine, biologically relevant, disease-associated group.

Supplementary Figure 34:

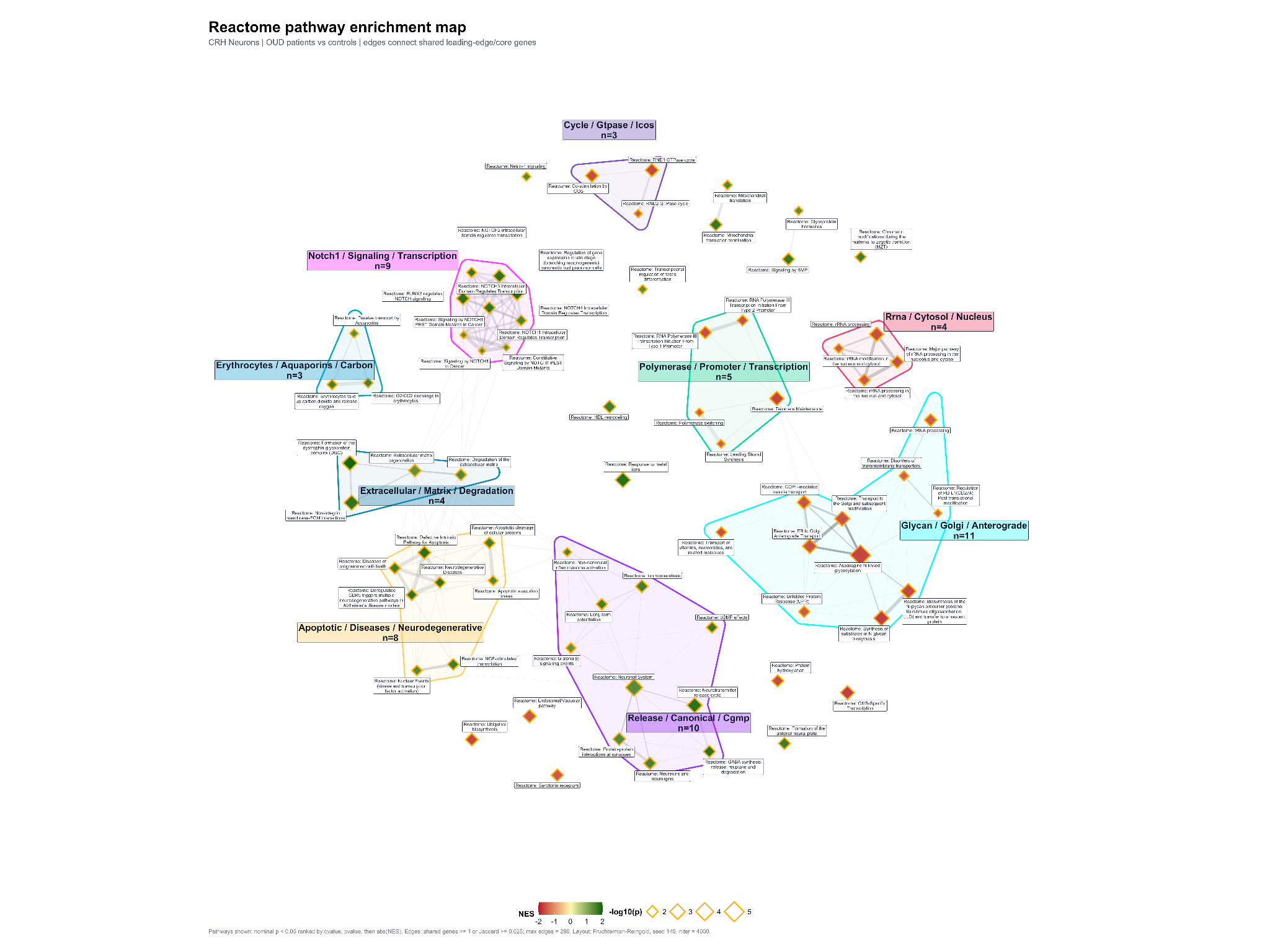

**Reactome Pathway Enrichment Map.**

Reactome pathway enrichment map showing overlap among nominally enriched Reactome pathways in CRH neurons from OUD patients versus controls. Nodes represent Reactome pathways, node fill represents NES, node size represents -log10(nominal pvalue), and orange diamond borders denote Reactome source identity. Edges connect pathways sharing leading-edge/core genes, with edge width scaled by Jaccard overlap and edge transparency scaled by shared-gene count. Colored hulls indicate Louvain modules detected from weighted pathway-overlap structure; bold colored labels summarize modules using keyword-derived labels from member pathway descriptions. Generation specifications: 72 pathways shown; 168 edges shown. Pathways selected using nominal pvalue < 0.05 and ranked by qvalue, pvalue, then absolute NES. Edges retained using shared genes >= 1 or Jaccard >= 0.025, max_edges = 280. Layout: Fruchterman-Reingold, seed = 149, niter = 4000. Clustering: Louvain community detection weighted by Jaccard overlap. Node fill: NES, red = higher in controls, yellow = near zero, green = higher in OUD.

Supplementary Figure 35:

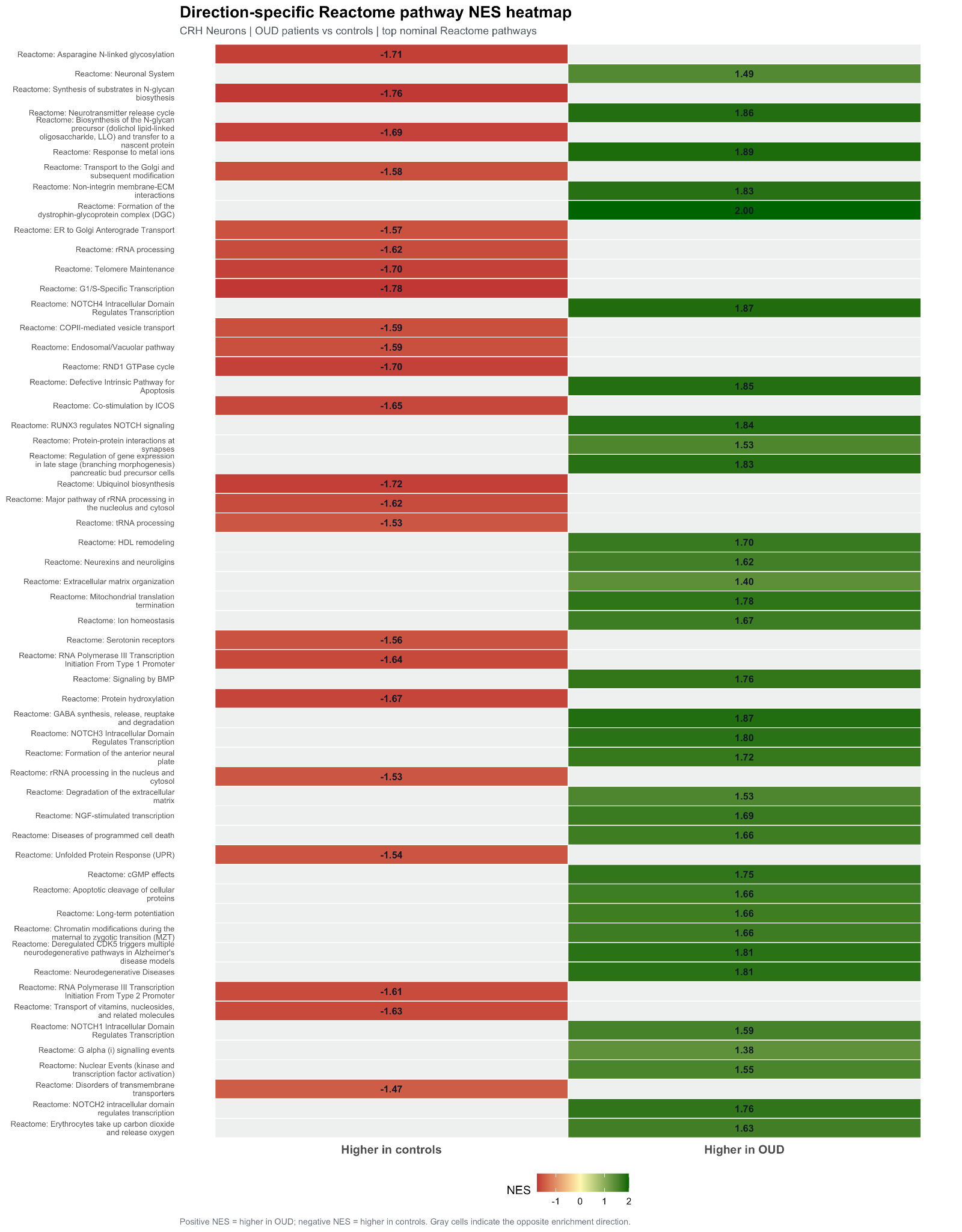

**Direction-Specific Reactome Pathway NES Heatmap.**

Direction-specific heatmap of top-ranked Reactome pathway normalized enrichment scores in CRH neurons. Each row represents a Reactome pathway, and columns separate pathways higher in controls from pathways higher in OUD. Colored tiles show the pathway NES only in the matching direction column; gray cells indicate the opposite direction. Positive NES indicates higher enrichment in OUD, and negative NES indicates higher enrichment in controls. Generation specifications: 56 pathways shown. Pathways selected from nominally enriched Reactome results using pvalue < 0.05 and ranked by qvalue, pvalue, then absolute NES. Tile fill uses a red/yellow/green NES gradient: red = higher in controls, yellow = near zero, green = higher in OUD.

Supplementary Figure 36:

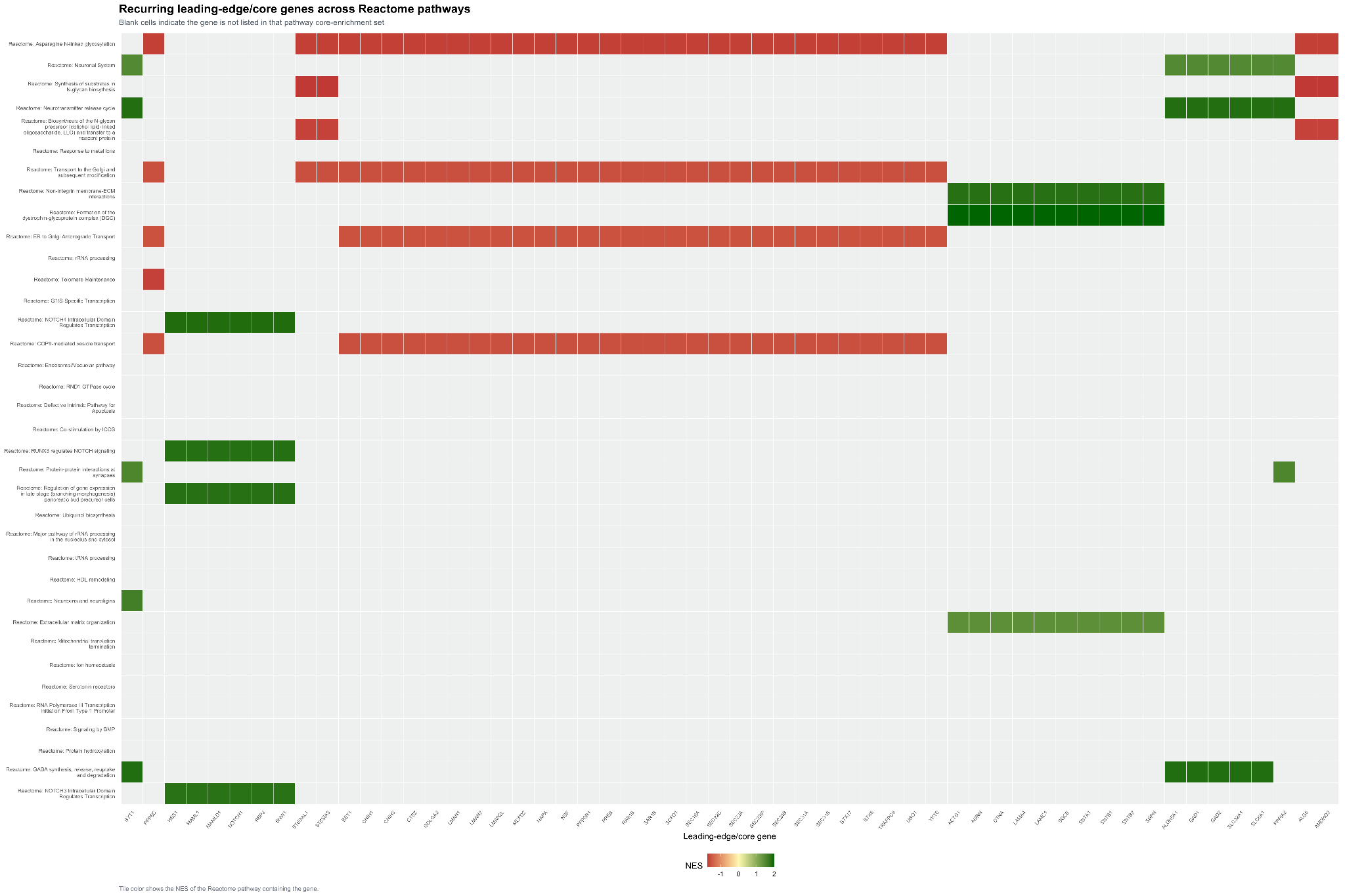

**Recurring Leading-Edge/Core Genes Across Reactome Pathways.**

Heatmap of recurring leading-edge/core genes across selected Reactome pathways in CRH neurons. Rows represent top-ranked Reactome pathways, columns represent recurrent leading-edge/core genes, and colored cells indicate that the gene was present in the pathway core-enrichment set. Tile color shows the NES of the Reactome pathway containing the gene. Blank cells indicate that the gene was not listed in that pathway core-enrichment set. Generation specifications: 36 Reactome pathways and 56 leading-edge/core genes shown. Pathways ranked by qvalue, pvalue, then absolute NES. Genes ranked by recurrence across selected pathways, maximum absolute pathway NES, best pathway pvalue, and gene symbol. Tile fill uses NES direction coloring.

Supplementary Figure 37:

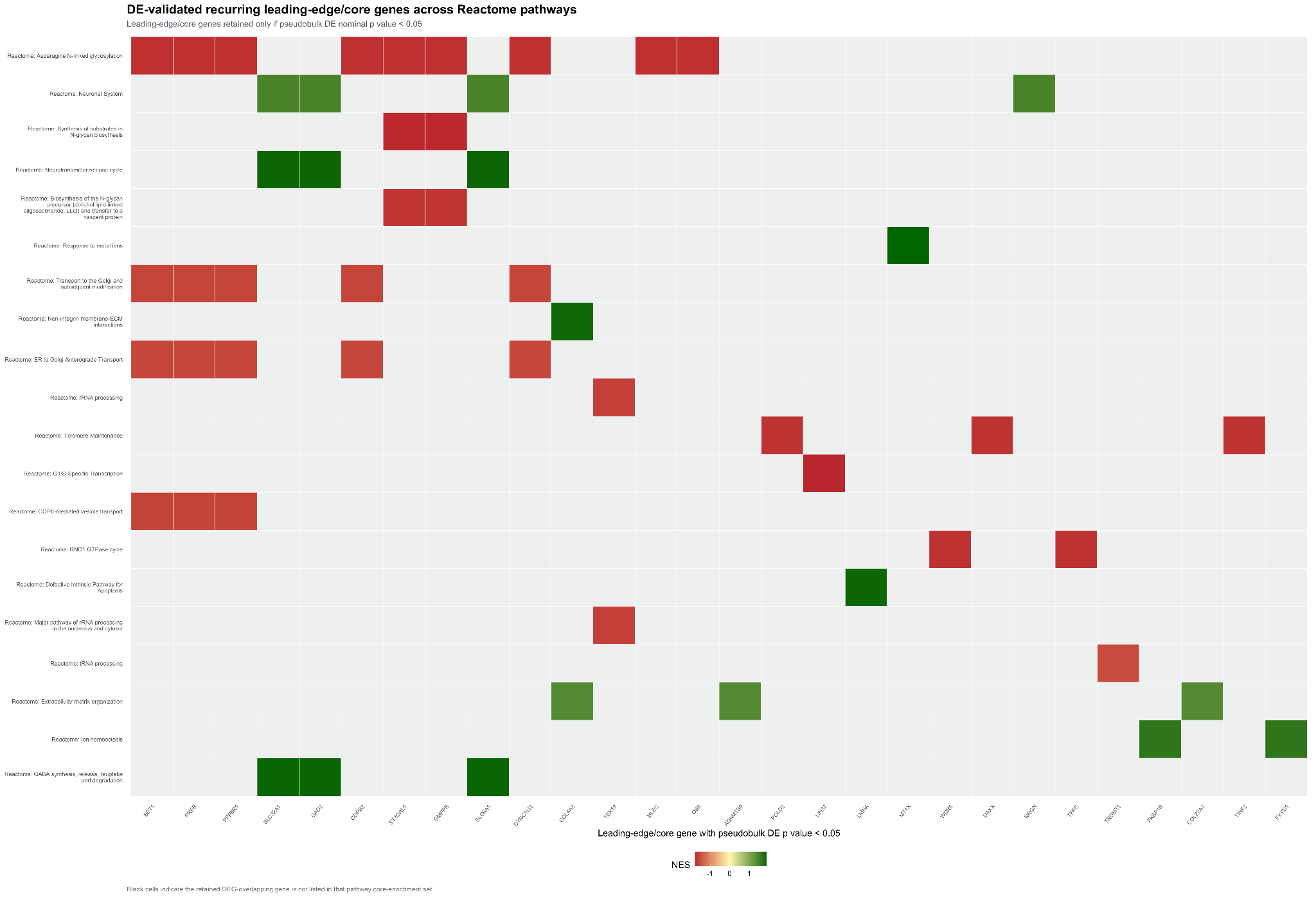

**DE-Validated Recurring Leading-Edge/Core Genes Across Reactome Pathways.**

Pseudobulk differential expression-validated heatmap of recurring Reactome leading-edge/core genes in CRH neurons. Genes were retained only if they were present in the CRH pseudobulk differential expression table with nominal pvalue < 0.05. Rows represent Reactome pathways retained after gene filtering, columns represent DE-overlapping leading-edge/core genes, and colored cells indicate gene membership in each pathway core-enrichment set. Generation specifications: 28 genes and 20 Reactome pathways shown after pseudobulk DE filtering. Pathways selected from the top 36 ranked Reactome pathways. Gene validation used pseudobulk DE pvalue < 0.05. Tile fill uses pathway NES direction coloring.

Supplementary Figure 38:

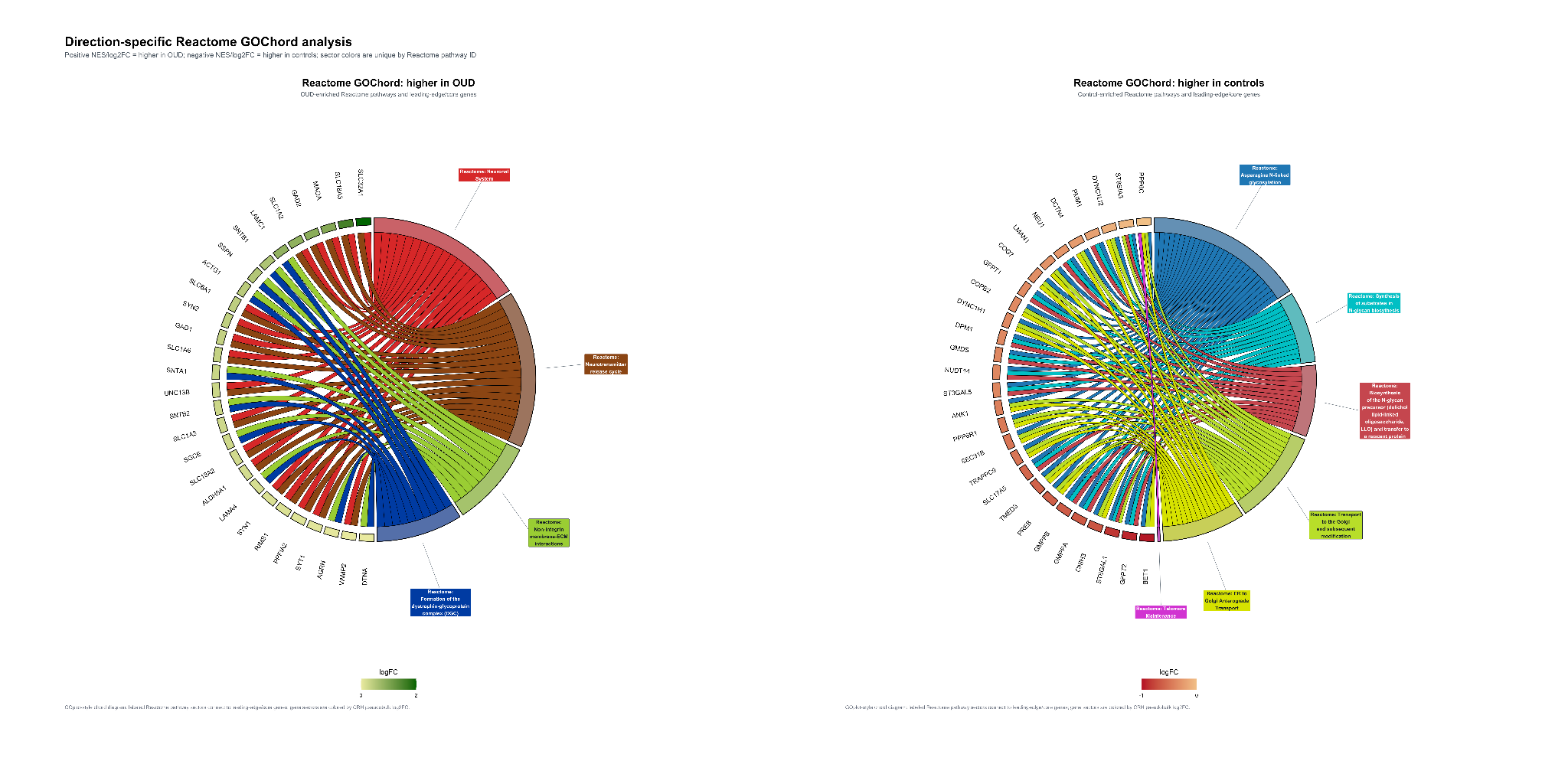

**Direction-Specific Reactome GOChord Analysis.**

Direction-specific GOChord visualization of Reactome pathways and leading-edge/core genes in CRH neurons. The left panel shows OUD-enriched Reactome pathways and their leading-edge/core genes; the right panel shows control-enriched Reactome pathways and their leading-edge/core genes. Reactome pathway sectors are assigned unique colors by pathway ID, ribbons connect genes to their corresponding Reactome pathways, and gene sectors are colored by CRH pseudobulk log2FC.

Supplementary Figure 39:

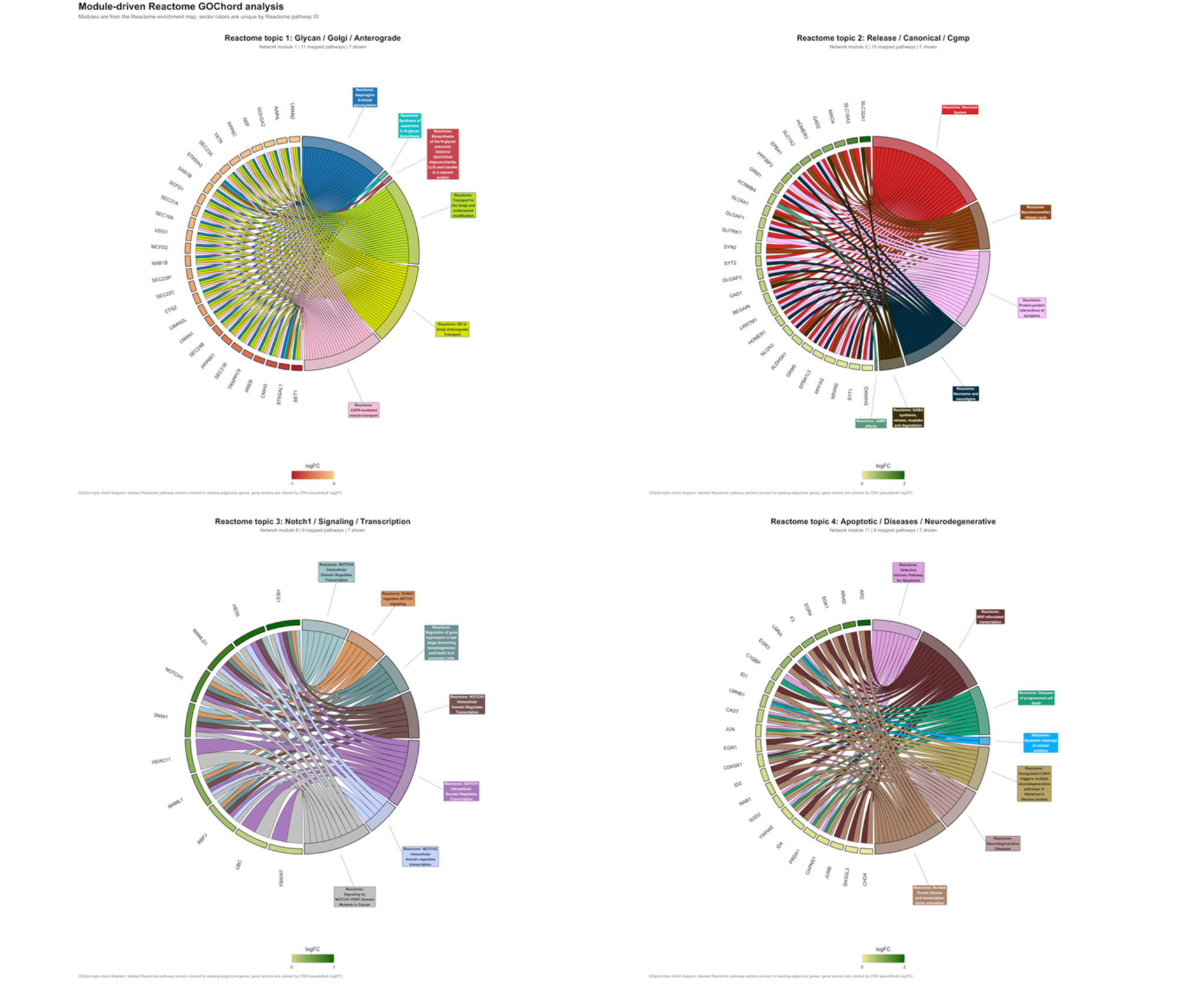

**Module-Driven Reactome GOChord Analysis.**

Module-driven GOChord analysis of Reactome pathway-overlap modules in CRH neurons. Modules were defined from the Reactome enrichment map using Louvain community detection weighted by Jaccard overlap of leading-edge/core genes. Each panel shows a selected Louvain module, with Reactome pathway sectors connected to leading-edge/core genes by colored ribbons. Pathway sectors are uniquely colored by pathway ID, and gene sectors are colored by CRH pseudobulk log2FC. Generation specifications: 9 Louvain modules were detected from the Reactome enrichment map; 47 pathways were selected across modules using up to 7 pathways per module. The combined figure displays the top 4 modules ranked by pathway count, minimum pvalue, and absolute mean NES: Glycan / Golgi / Anterograde, Release / Canonical / Cgmp, Notch1 / Signaling / Transcription, and Apoptotic / Diseases / Neurodegenerative. GOChord settings matched Figure 05, with module panels using up to 28 genes.

Supplementary Figure 40:

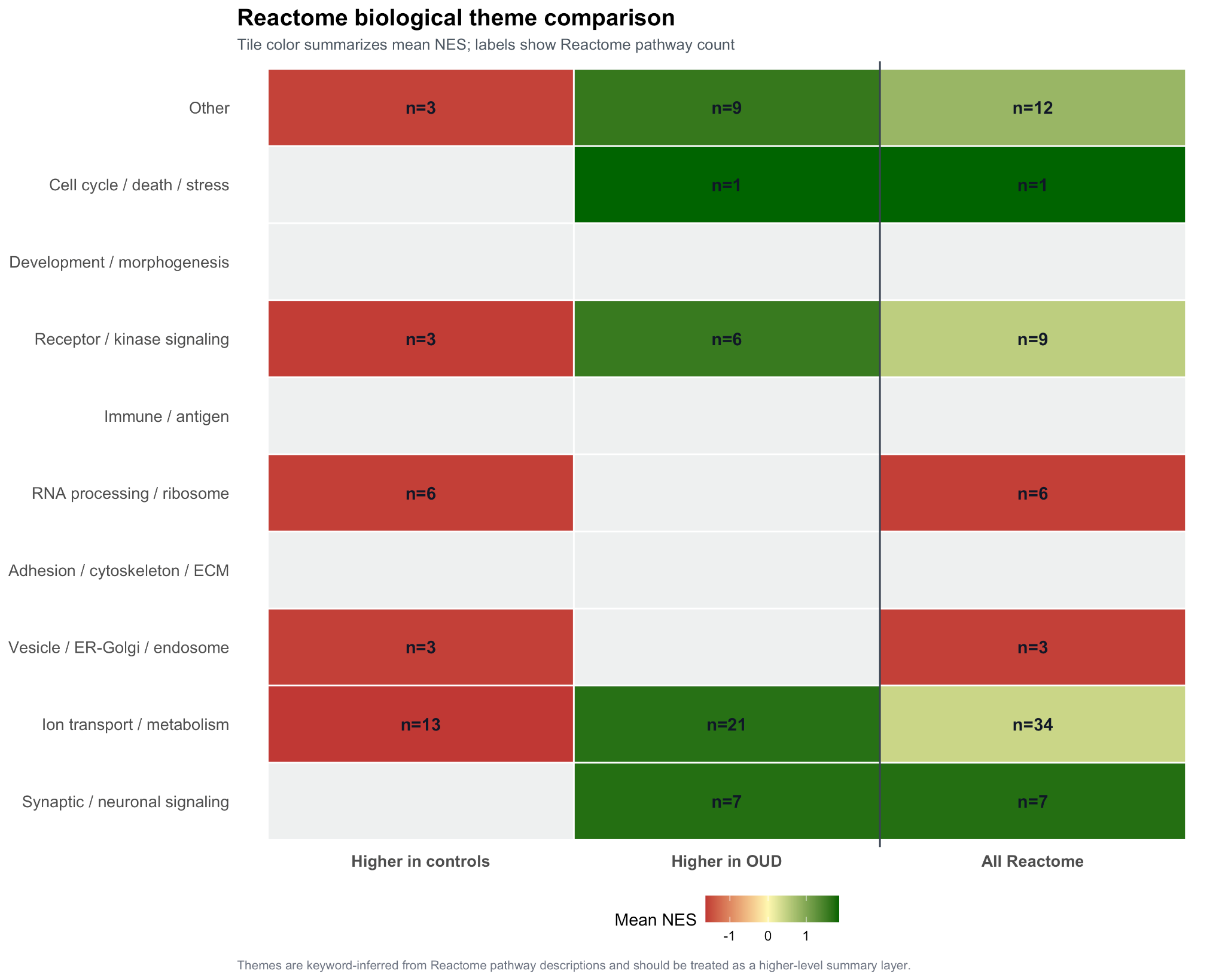

**Reactome Biological Theme Comparison.**

Biological-theme comparison of Reactome pathway enrichment in CRH neurons. Rows represent keyword-inferred biological themes, and columns summarize pathways higher in controls, pathways higher in OUD, and all selected Reactome pathways. Tile color indicates mean NES for each theme-by-direction group, and labels report Reactome pathway counts. Generation specifications: Themes were assigned by keyword matching to Reactome pathway descriptions, independent of Louvain network clustering. Directional groups were assigned from NES sign. Tile fill represents mean NES with red = higher in controls, yellow = near zero, and green = higher in OUD. Displayed theme groups include Synaptic / neuronal signaling, Ion transport / metabolism, Vesicle / ER-Golgi / endosome, Adhesion / cytoskeleton / ECM, RNA processing / ribosome, Immune / antigen, Receptor / kinase signaling, Development / morphogenesis, Cell cycle / death / stress, and Other.

Supplementary Figure 41:

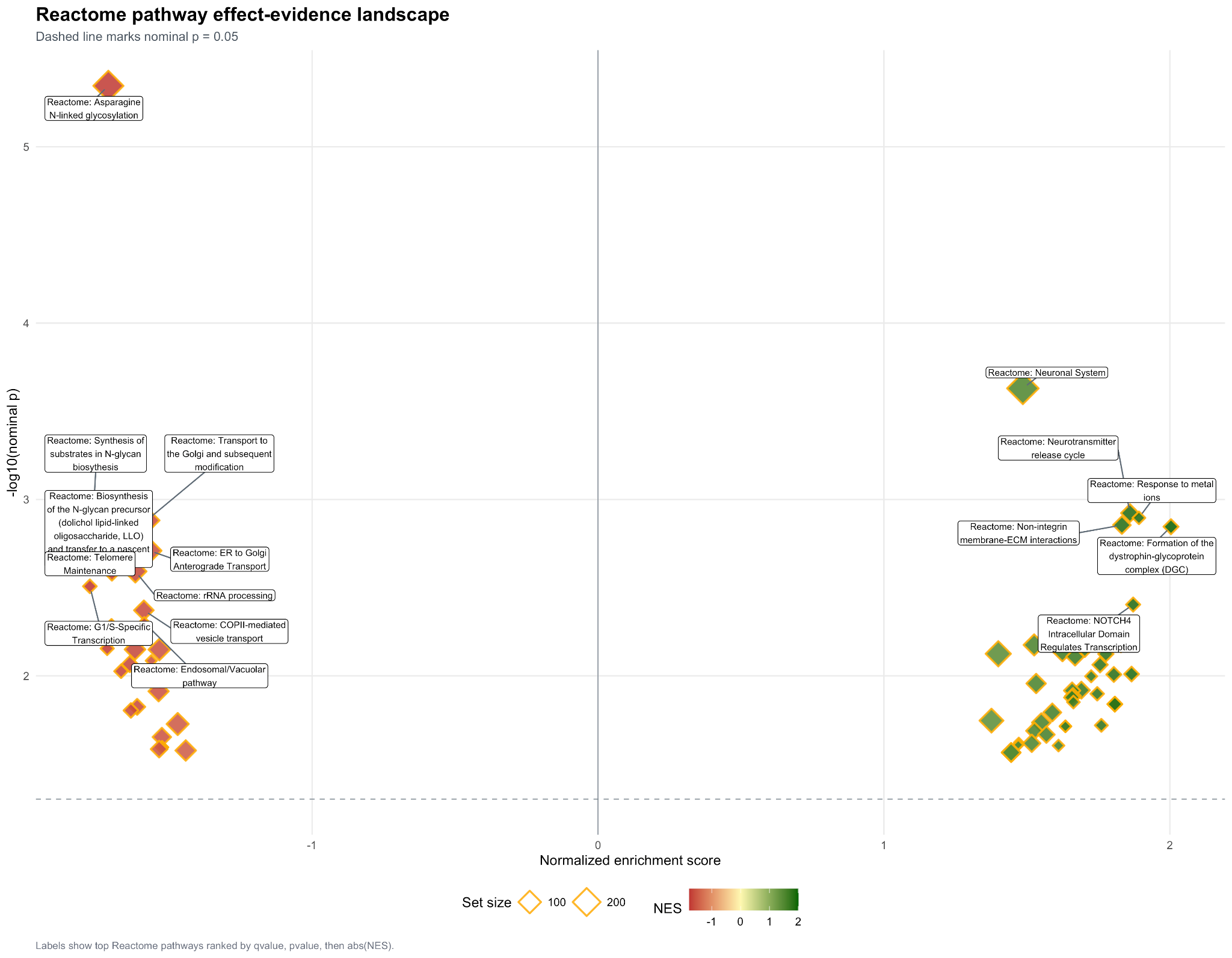

**Reactome Pathway Effect-Evidence Landscape.**

Reactome pathway effect-evidence landscape for CRH neurons comparing OUD patients with controls. Each point represents a Reactome pathway plotted by NES on the x-axis and -log10(nominal pvalue) on the y-axis. Point size represents Reactome pathway set size, and point fill represents NES direction and magnitude. The dashed horizontal line marks nominal pvalue = 0.05. Labeled pathways are the top-ranked terms by qvalue, pvalue, and absolute NES. Generation specifications: 72 selected nominal Reactome pathways shown. Labels applied to the top 16 pathways ranked by qvalue, pvalue, then absolute NES. Point shape: Reactome diamond, border `#FFB000`; fill: NES gradient.

Supplementary Figure 42:

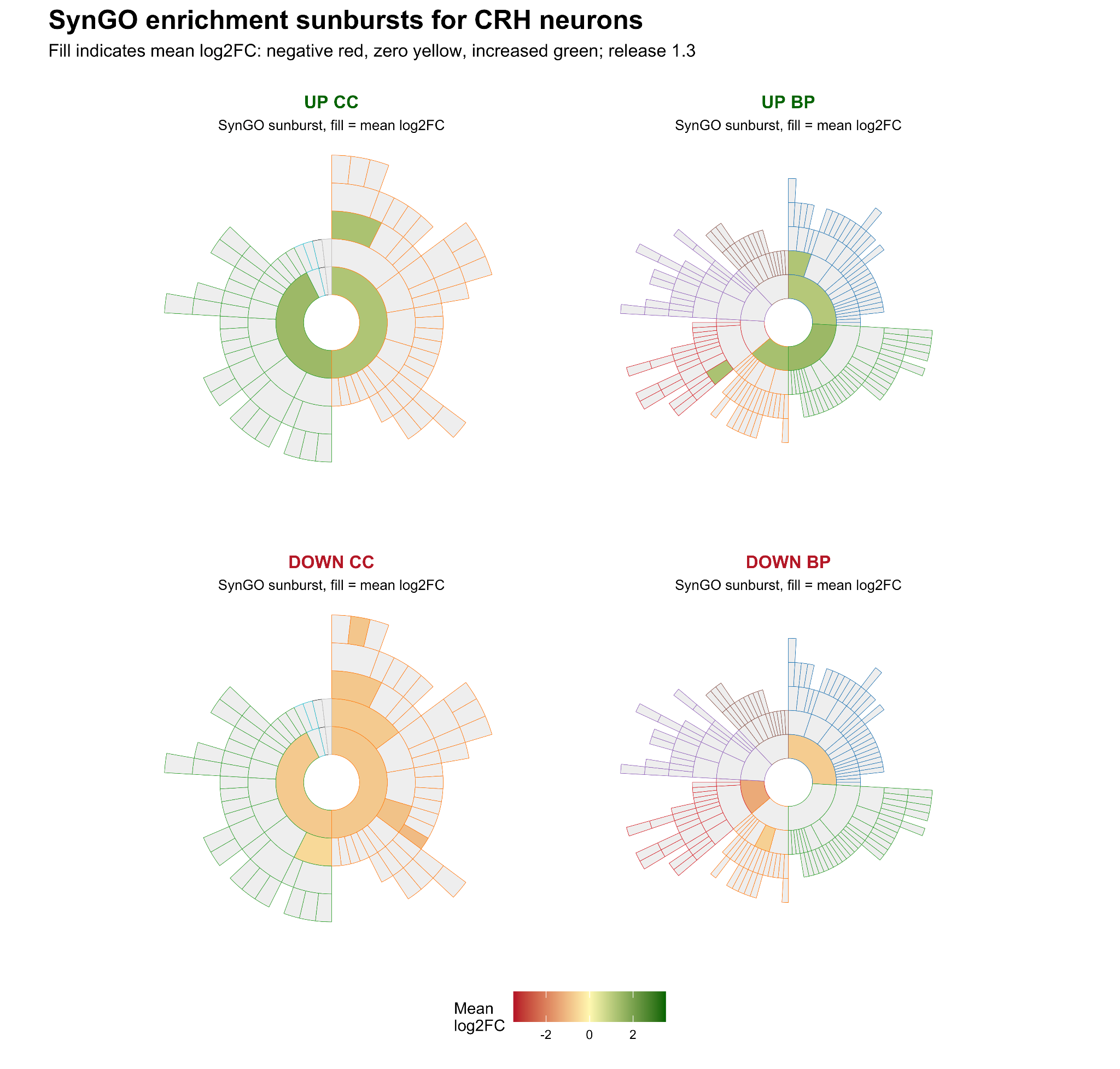

SynGO annotation of differentially expressed genes in CRH neurons.

SynGO sunburst plots summarize synaptic cellular component (CC) and biological process (BP) annotations for genes increased (UP) or decreased (DOWN) in CRH neurons identified by pseudobulk single-nucleus RNA-seq differential expression analysis. Foreground gene lists were defined using p < 0.05 and |log2FC| > 0.25; 77 UP genes and 171 DOWN genes were submitted. Of these, 13 UP genes and 16 DOWN genes mapped to the SynGO/HGNC lookup, against a background of 10,214 genes, including 1,408 SynGO-annotated genes. Filled sunburst sectors indicate SynGO terms containing at least three recursively annotated foreground genes; unfilled sectors represent ontology terms without sufficient foreground annotation. Sector fill denotes the mean foreground log2 fold-change for genes annotated to each term, clipped from -2 to 2, with decreased expression shown in red, zero centered at yellow, and increased expression shown in green. In UP genes, mapped SynGO annotations included postsynaptic density, postsynapse, presynapse, synapse assembly, synapse organization, presynaptic and postsynaptic processes, and chemical synaptic transmission. In DOWN genes, mapped annotations included postsynapse, presynapse, postsynaptic membrane/density terms, synaptic vesicle, synapse organization, synaptic signaling, and regulation of postsynaptic membrane neurotransmitter receptor levels. Overrepresentation testing used one-sided Fisher’s exact tests with Benjamini-Hochberg correction within each SynGO domain and direction; no CC or BP gene-cluster terms reached q <= 0.01. SynGO release 1.3 was used.

Supplementary Figure 43:

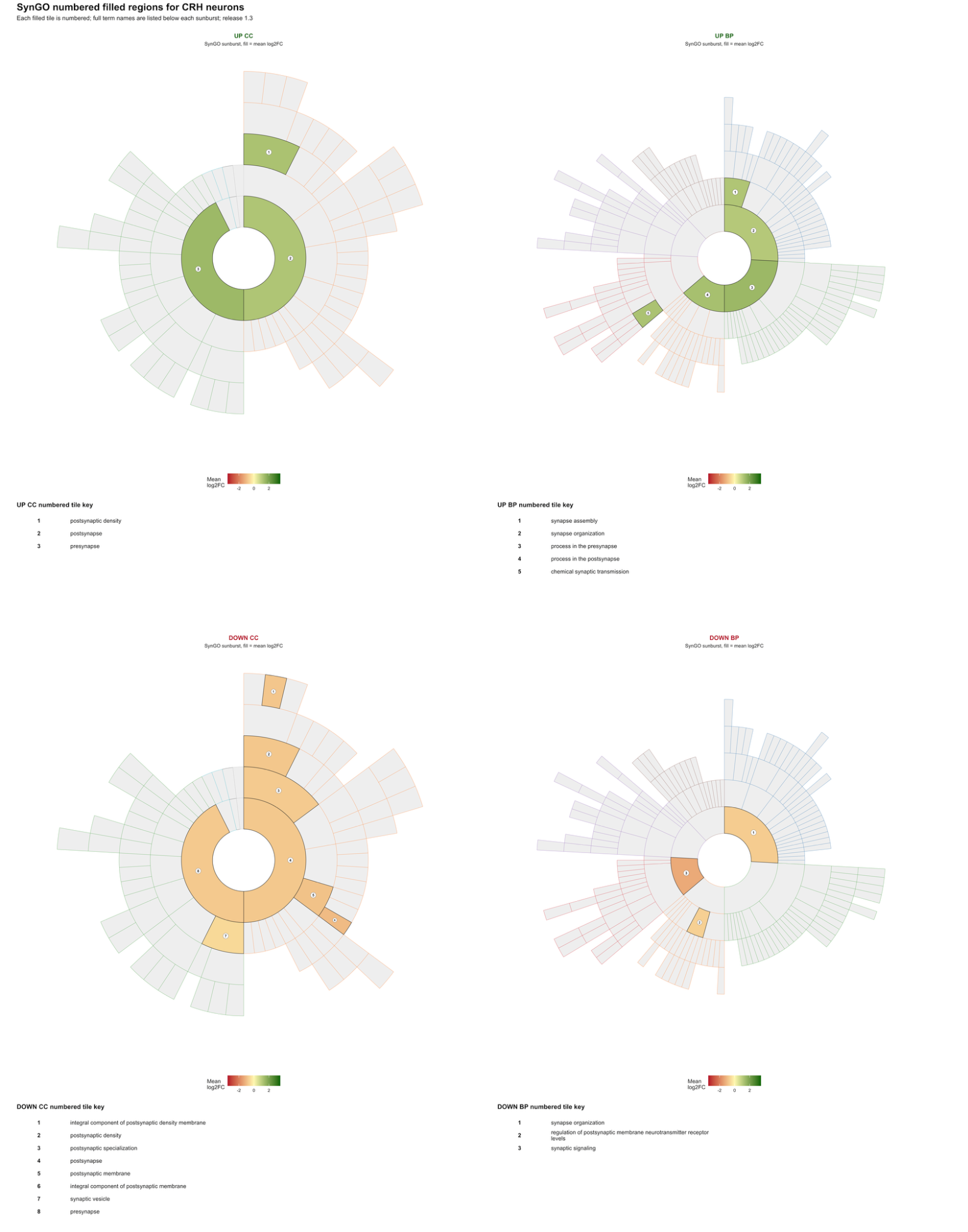

**Numbered SynGO sunburst annotations for CRH neuron differentially expressed genes.**

Numbered SynGO sunburst plots show cellular component (CC) and biological process (BP) annotations for genes increased (UP) or decreased (DOWN) in CRH neurons from pseudobulk single-nucleus RNA-seq differential expression analysis. Foreground genes were defined as p < 0.05 and |log2FC| > 0.25; 13 UP and 16 DOWN submitted genes mapped to SynGO. Filled sectors indicate SynGO ontology tiles with at least three recursively annotated foreground genes, and each numbered sector is defined in the corresponding tile key below the plot. Fill color represents mean foreground log2 fold-change for genes assigned to that tile, clipped from -2 to 2, with decreased expression in red, zero centered at yellow, and increased expression in green. UP-associated filled terms included postsynaptic density, postsynapse, presynapse, synapse assembly, synapse organization, presynaptic/postsynaptic processes, and chemical synaptic transmission. DOWN-associated filled terms included postsynaptic density and membrane components, postsynapse, presynapse, synaptic vesicle, synapse organization, synaptic signaling, and regulation of postsynaptic membrane neurotransmitter receptor levels. SynGO release 1.3 was used; no CC or BP gene-cluster terms reached q <= 0.01 after Benjamini-Hochberg correction.

Supplementary Figure 44:

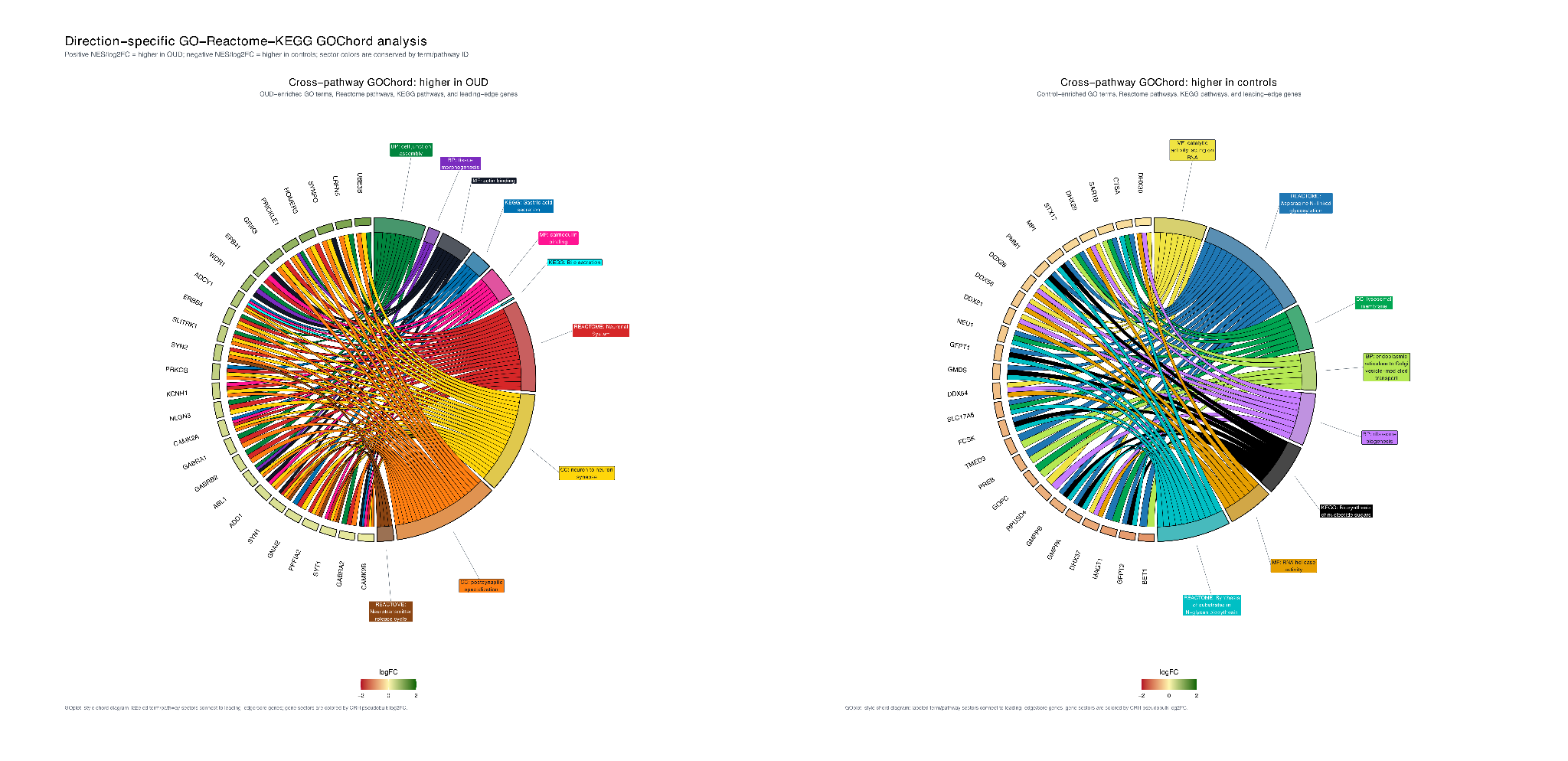

**Direction-Specific GO-Reactome-KEGG GOChord Analysis in CRH Neurons.**

Direction-specific GOChord plots showing relationships between enriched GO terms, Reactome pathways, KEGG pathways, and leading-edge/core genes in CRH neurons comparing OUD patients with controls. The left panel shows pathways/terms with positive NES values, corresponding to higher enrichment in OUD, and the right panel shows pathways/terms with negative NES values, corresponding to lower enrichment in OUD, or higher enrichment in controls. Outer sectors represent selected GO, Reactome, or KEGG terms/pathways, with unique colors assigned to each term/pathway ID. Ribbons connect each term/pathway sector to its displayed leading-edge/core genes. Gene sectors are colored by CRH pseudobulk log2FC.

Supplementary Figure 45:

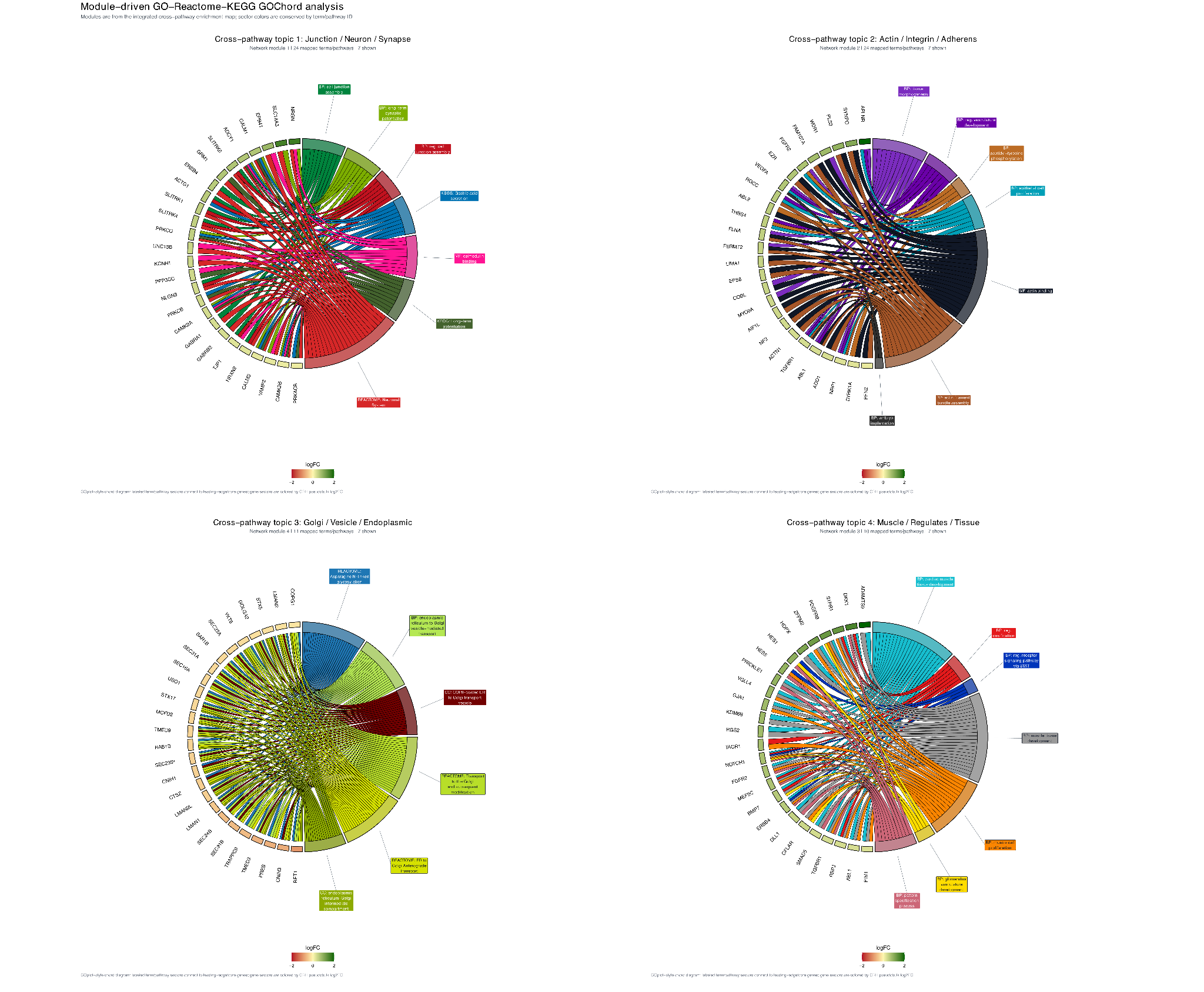

**Module-driven GO–Reactome–KEGG pathway analysis in CRH neurons.**

GOChord plots showing leading-edge/core-gene overlap within the top four modules identified from the integrated GO–Reactome–KEGG enrichment network comparing OUD and control CRH neurons. Each panel represents one Louvain-derived network module. Outer sectors denote selected GO, Reactome, or KEGG terms/pathways, with ribbons linking each term to associated leading-edge/core genes. Gene sectors are colored by CRH pseudobulk log2 fold change, and pathway sectors are uniquely colored by term identity.

Supplementary Figure 46:

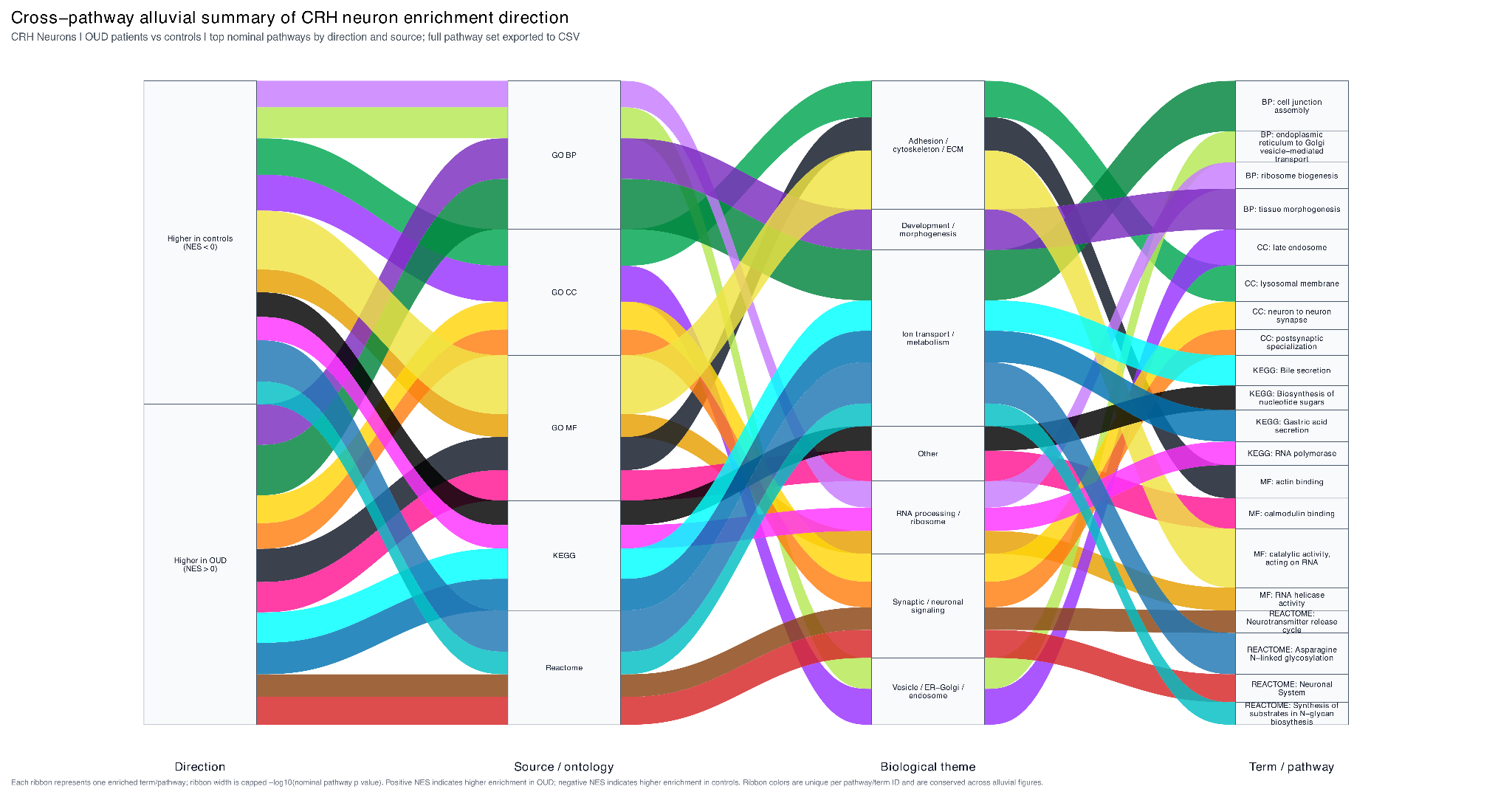

**Cross-pathway alluvial summary of CRH neuron enrichment direction**.

Alluvial plot summarizing nominally enriched GO biological process (GO BP), GO cellular component (GO CC), GO molecular function (GO MF), Reactome, and KEGG terms/pathways in CRH neurons from OUD patients versus controls. The plot displays 20 representative terms/pathways selected from the nominally significant integrated pathway set, with 10 showing positive normalized enrichment scores (NES; higher in OUD) and 10 showing negative NES (higher in controls). Flows connect enrichment direction, source/ontology, keyword-derived biological theme, and individual term/pathway label. Ribbon width represents capped -log10(nominal pathway p value), and ribbon color is assigned uniquely to each term/pathway ID. Higher-in-OUD terms/pathways shown: GO BP: cell junction assembly (NES=1.69, pathway p=2.89e-07, q=0.002), tissue morphogenesis (NES=1.79, pathway p=5.41e-06, q=0.016); GO CC: neuron to neuron synapse (NES=1.45, pathway p=2.42e-04, q=0.032); postsynaptic specialization (NES=1.39, pathway p=4.71e-04, q=0.052); GO MF: actin binding (NES=1.55, pathway p=5.03e-05, q=0.030); calmodulin binding (NES=1.70, pathway p=1.14e-04, q=0.045); Reactome: Neuronal System (NES=1.49, pathway p=2.34e-04, q=0.201), Neurotransmitter release cycle (NES=1.86, pathway p=0.001, q=0.272); KEGG: Bile secretion (NES=2.11, pathway p=1.16e-04, q=0.015); Gastric acid secretion (NES=2.14, pathway p=8.08e-05, q=0.015). Higher-in-controls terms/pathways shown: GO BP: endoplasmic reticulum to Golgi vesicle-mediated transport (NES=-1.78, pathway p=9.50e-05, q=0.029), ribosome biogenesis (NES=-1.58, pathway p=3.71e-04, q=0.056); GO CC: late endosome (NES=-1.72, pathway p=2.21e-05, q=0.005), lysosomal membrane (NES=-1.64, pathway p=1.91e-05, q=0.005); GO MF: catalytic activity, acting on RNA (NES=-1.87, pathway p=2.29e-08, q=2.70e-05), RNA helicase activity (NES=-1.75, pathway p=0.001, q=0.152), Asparagine N-linked glycosylation (NES=-1.71, pathway p=4.51e-06, q=0.008); Reactome: Synthesis of substrates in N-glycan biosynthesis (NES=-1.76, pathway p=0.001, q=0.272); KEGG: Biosynthesis of nucleotide sugars (NES=-1.87, pathway p=6.59e-04, q=0.030), RNA polymerase (NES=-1.91, pathway p=9.15e-04, q=0.030).

Supplementary Figure 47:

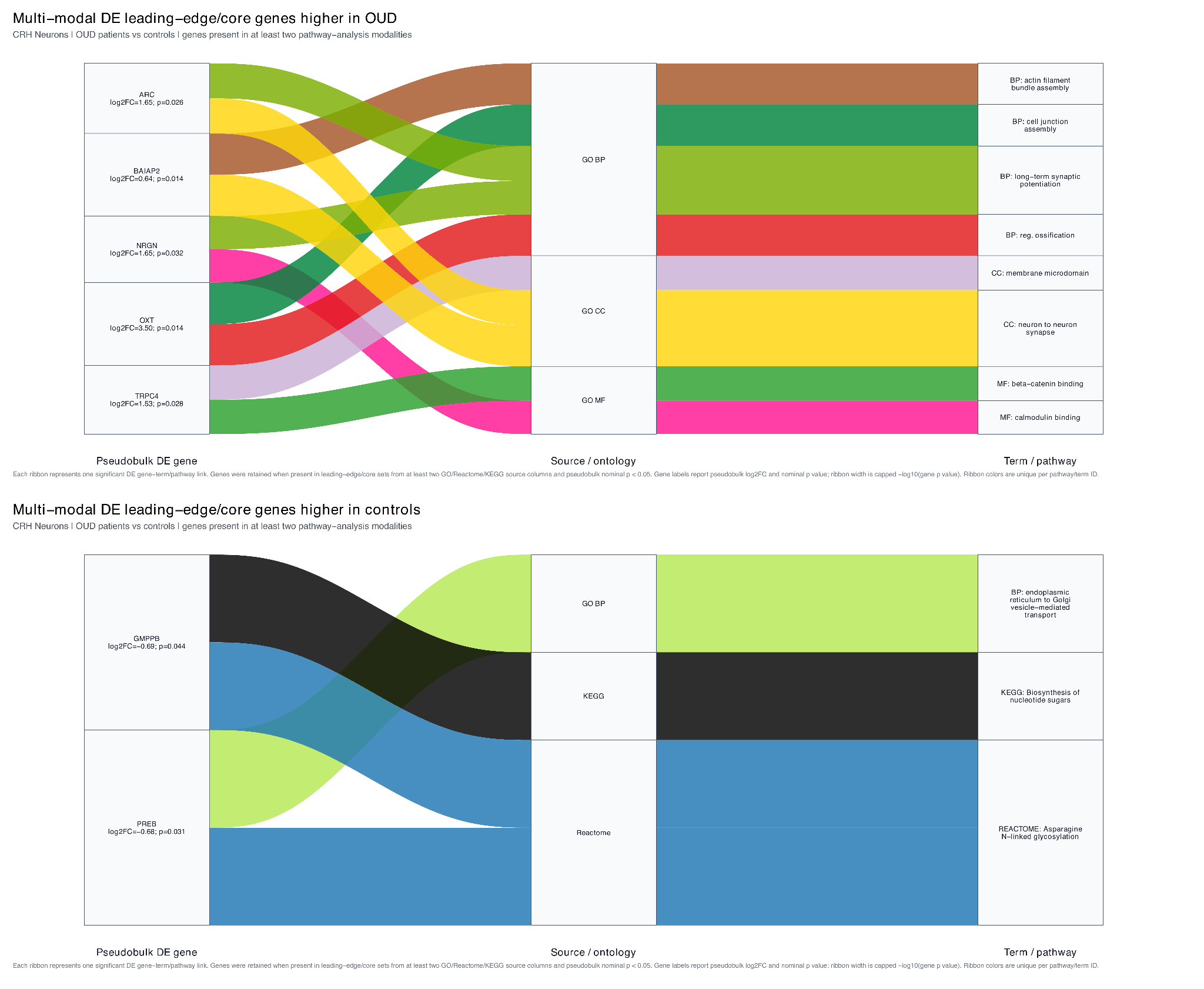

**Multi-modal DE leading-edge/core gene alluvial analysis.**

Combined version of Figures 13a and 13b restricted to pseudobulk-significant genes represented in leading-edge/core sets from at least two pathway-analysis modalities. Genes shown: ARC (log2FC=1.65, p=0.026); BAIAP2 (log2FC=0.64, p=0.014); GMPPB (log2FC=-0.69, p=0.044); NRGN (log2FC=1.65, p=0.032); OXT (log2FC=3.50, p=0.014); PREB (log2FC=-0.68, p=0.031); TRPC4 (log2FC=1.53, p=0.028). Terms/pathways shown: GO BP: actin filament bundle assembly (NES=1.82, pathway p=8.34e-05), cell junction assembly (NES=1.69, pathway p=2.89e-07), endoplasmic reticulum to Golgi vesicle-mediated transport (NES=-1.78, pathway p=9.50e-05), long-term synaptic potentiation (NES=2.04, pathway p=3.07e-05), regulation of ossification (NES=2.12, pathway p=1.93e-05); GO CC: membrane microdomain (NES=1.43, pathway p=0.003), neuron to neuron synapse (NES=1.45, pathway p=2.42e-04); GO MF: beta-catenin binding (NES=1.42, pathway p=0.010), calmodulin binding (NES=1.70, pathway p=1.14e-04); Reactome: Asparagine N-linked glycosylation (NES=-1.71, pathway p=4.51e-06); KEGG: Biosynthesis of nucleotide sugars (NES=-1.87, pathway p=6.59e-04).

Supplementary Figure 48:

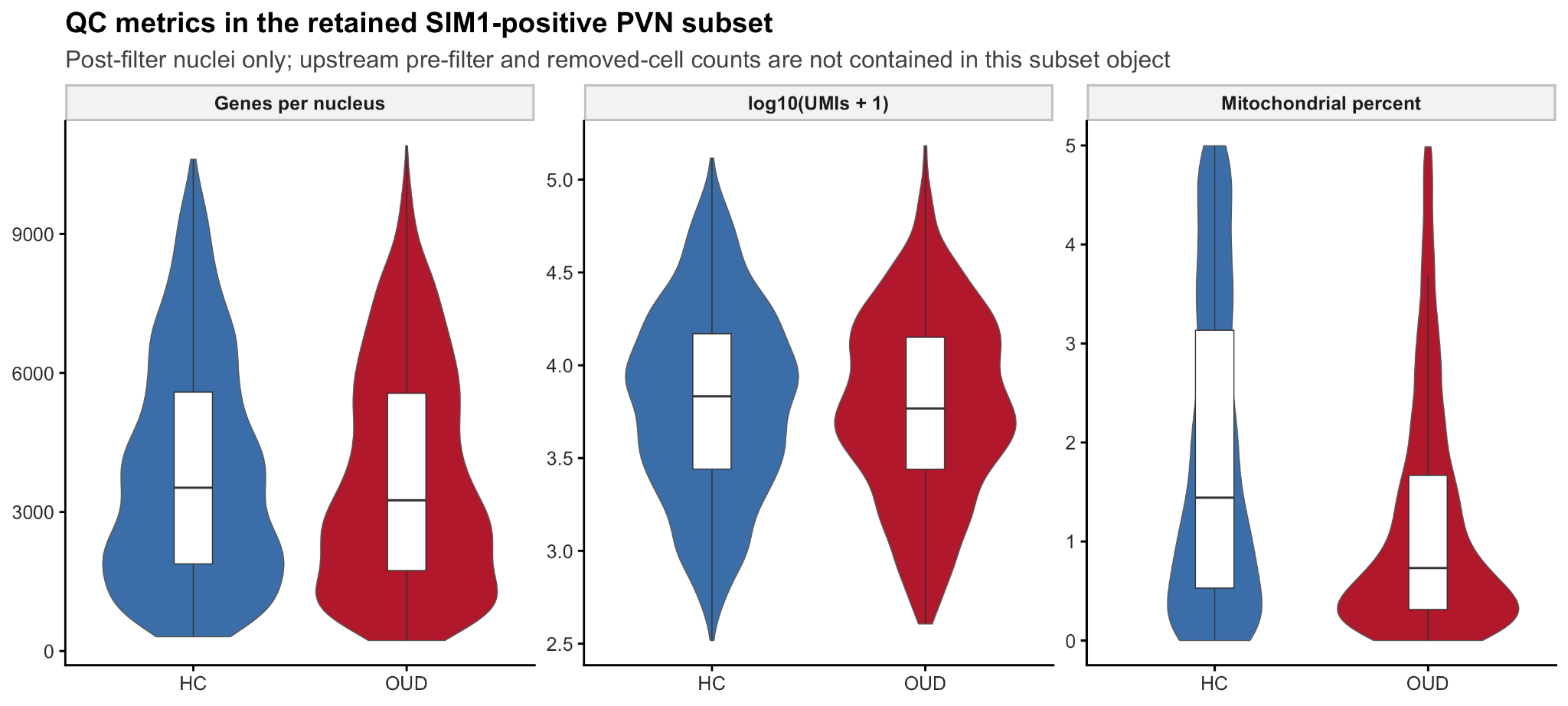

**Quality-control metrics in the available post-filter SIM1-positive PVN analytic subset.**

Violin and box plots show detected genes per nucleus, UMIs per nucleus, and mitochondrial read percentage stratified by the metadata-derived HC/OUD condition. The analysis condition was derived from the Opioid metadata column, with Non-Opioid coded as HC and Opioid coded as OUD. The available subset contained 2,847 nuclei from 25 donors, including 1,488 HC-associated nuclei from 15 donors and 1,359 OUD-associated nuclei from 10 donors. Across the full available subset, median detected genes per nucleus was 3,407, median UMIs per nucleus was 6,310, and median mitochondrial percentage was 1.05%. These metrics describe the downstream SIM1-positive PVN object only; full-cohort pre-filter and post-filter QC counts should be added from the original preprocessing workflow.

Supplementary Figure 49:

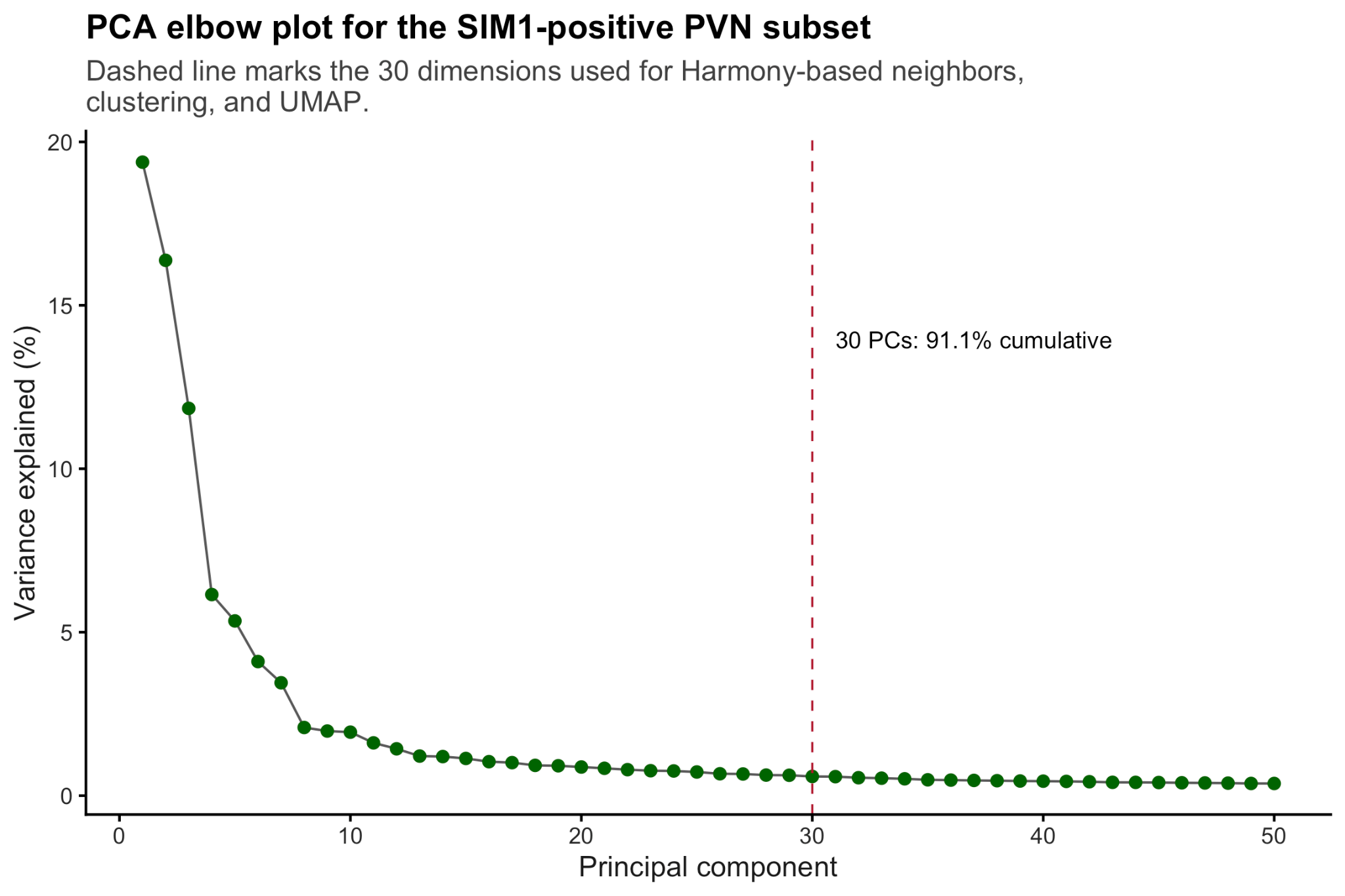

**PCA variance explained in the available SIM1-positive PVN subset.**

Bars show the percent variance explained by each principal component and the overlaid line shows cumulative variance explained. The first 30 PCs accounted for 91.1% of total PCA variance and were used as the dimensional input for Harmony-based clustering in the available annotation workflow.

Supplementary Figure 50:

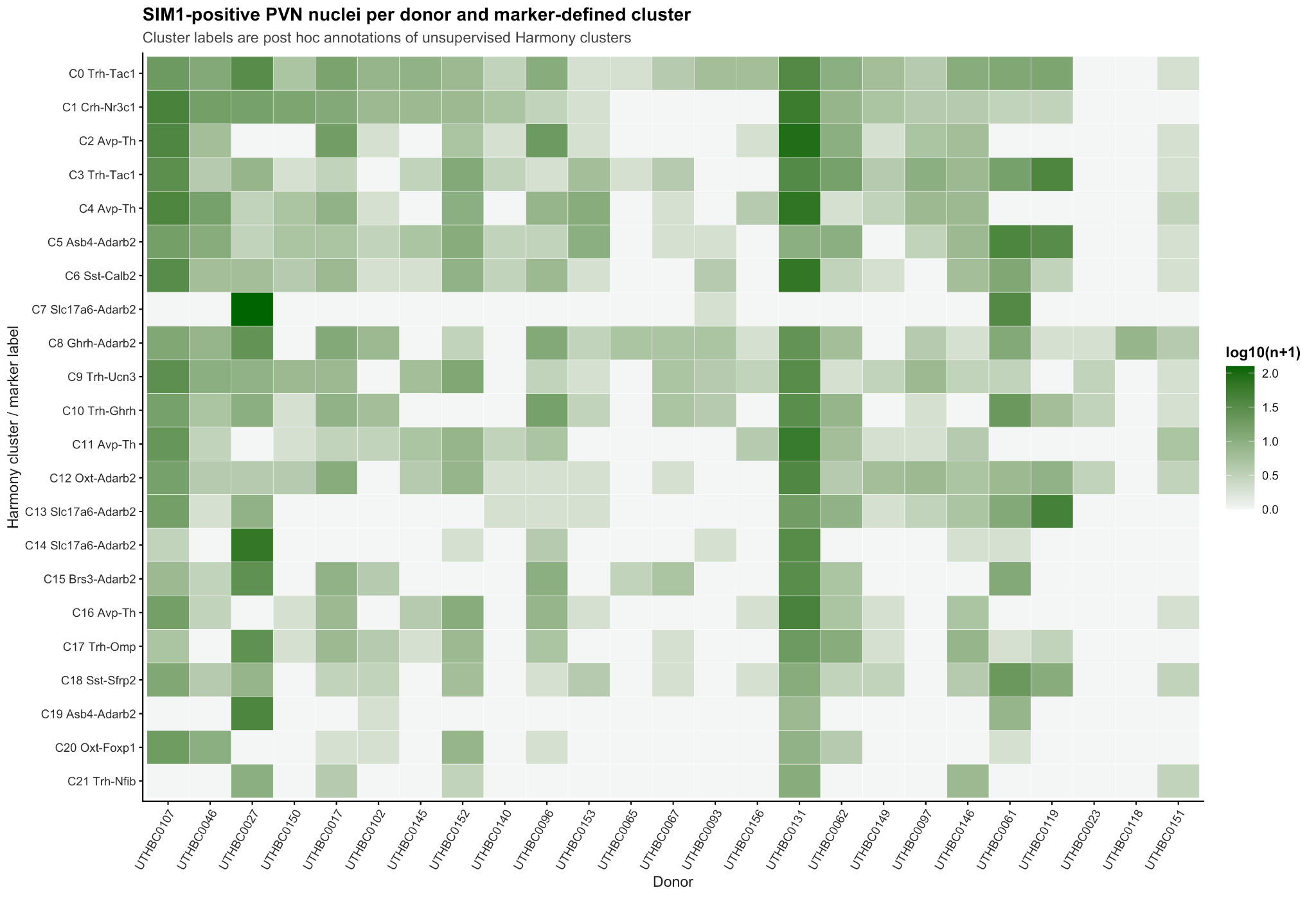

**Donor-level cell counts across marker-defined SIM1-positive PVN functional labels.**

Heatmap values represent the number of nuclei assigned to each functional label for each donor in the available post-filter SIM1-positive subset, with donors grouped by metadata-derived HC/OUD condition from the Opioid column. This visualization documents donor representation across annotated PVN neuronal populations and identifies labels with sparse donor-level coverage, an important consideration for downstream pseudobulk interpretation.

Supplementary Figure 51:

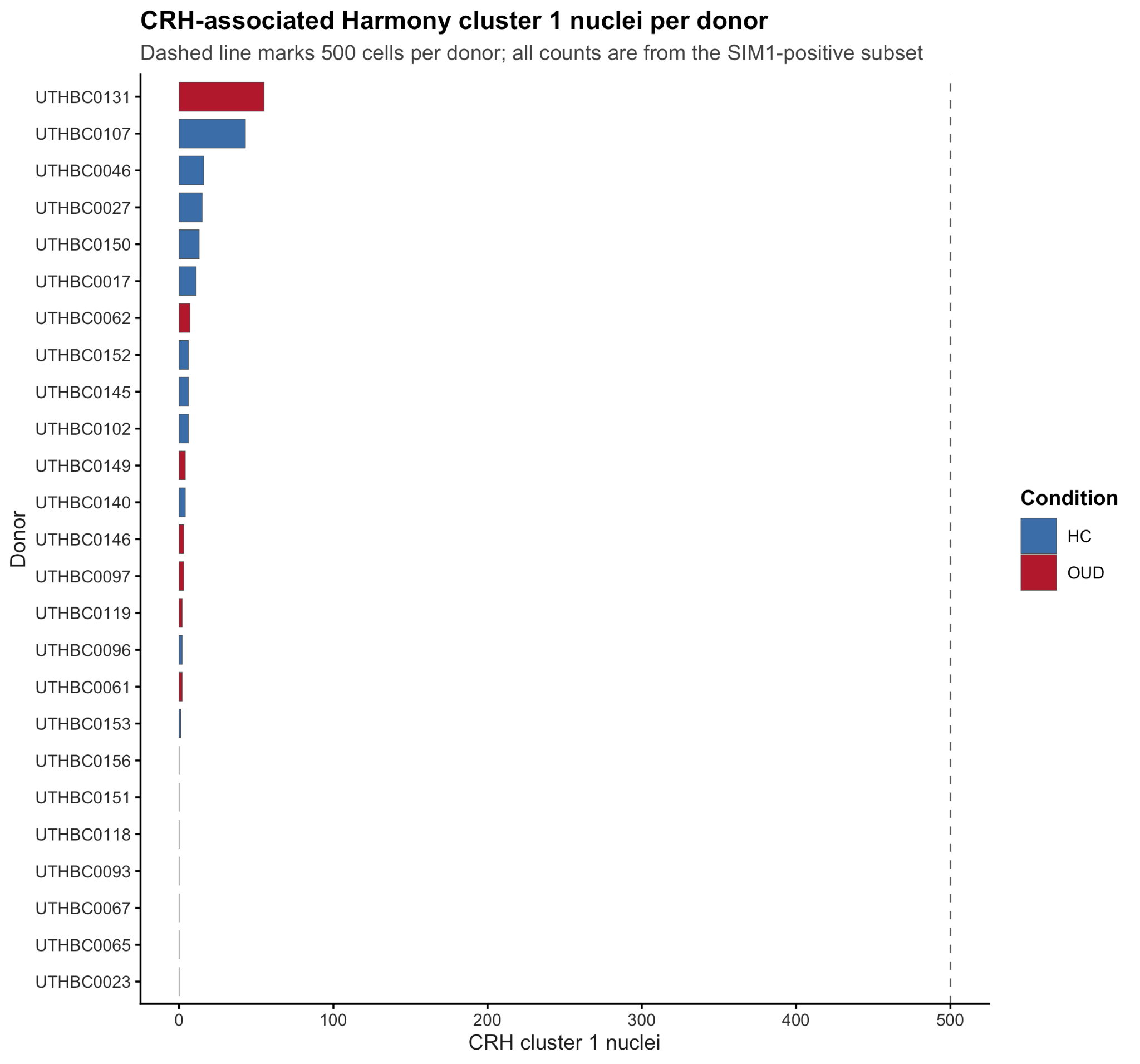

**Donor representation of the CRH-enriched Crh-Nr3c1 PVN population.**

Bars show the number of Harmony cluster 1 nuclei assigned to each donor in the available SIM1-positive subset, grouped by metadata-derived HC/OUD condition from the Opioid column. Harmony cluster 1 contained 199 nuclei overall, including 123 HC-associated nuclei and 76 OUD-associated nuclei. Among HC and OUD donors, the median number of CRH cluster 1 nuclei per donor was 6 and 2.5, respectively, and no donor reached 500 CRH cluster 1 nuclei.

Supplementary Figure 52:

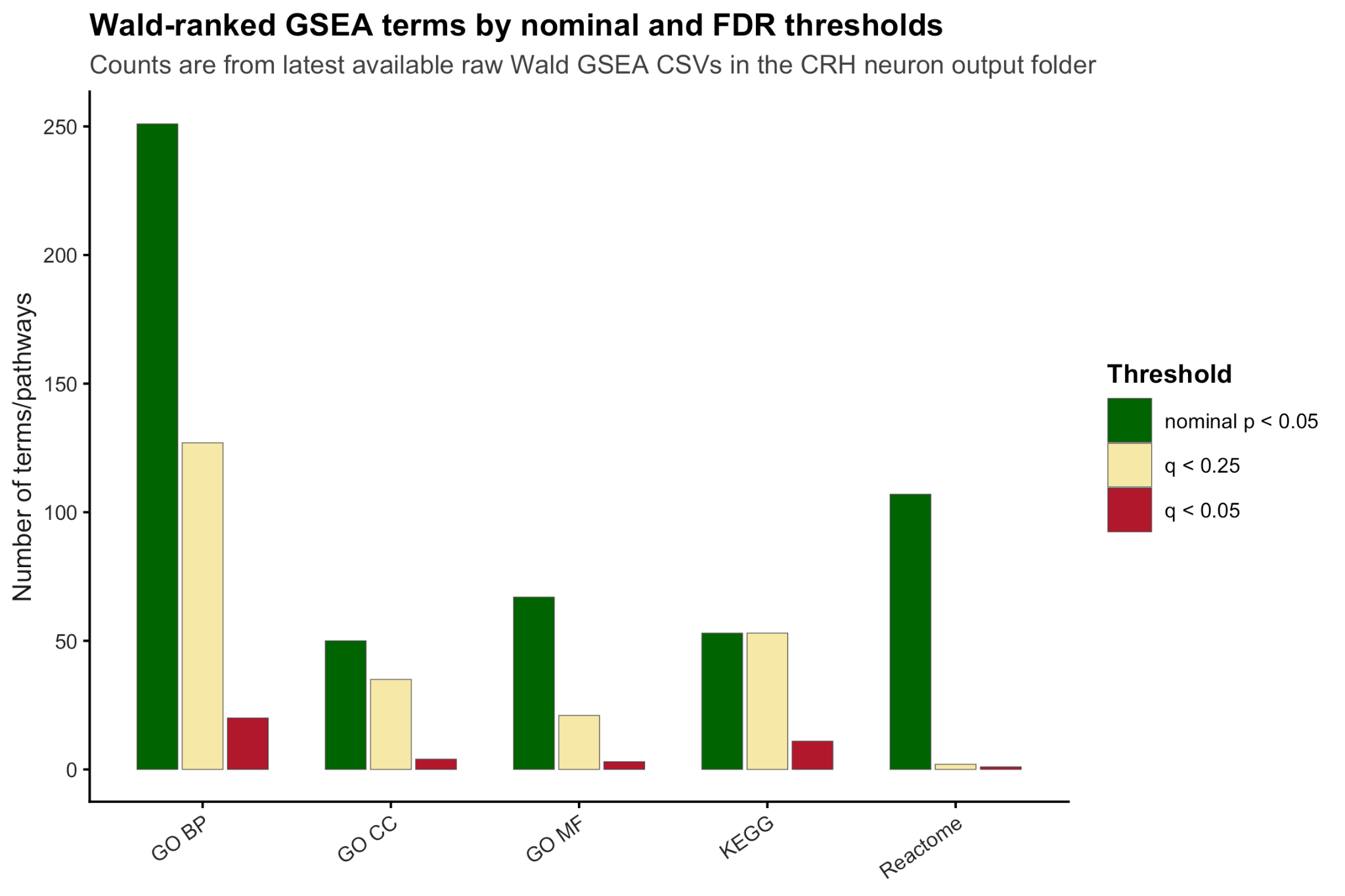

**Nominal and FDR-supported pathway counts for Wald-ranked GSEA.**

Bars compare the number of enriched terms meeting nominal P < 0.05, q < 0.25, q < 0.10, and q < 0.05 across GO biological process, GO cellular component, GO molecular function, KEGG, and Reactome analyses. Although many terms met nominal thresholds, FDR support varied by database, with q < 0.05 support for 20 GO BP terms, 4 GO CC terms, 3 GO MF terms, 11 KEGG terms, and 1 Reactome term. These counts should be reported alongside nominal P-value findings to clarify which pathway-level signals survive multiple-testing correction.

Supplementary Figure 53:

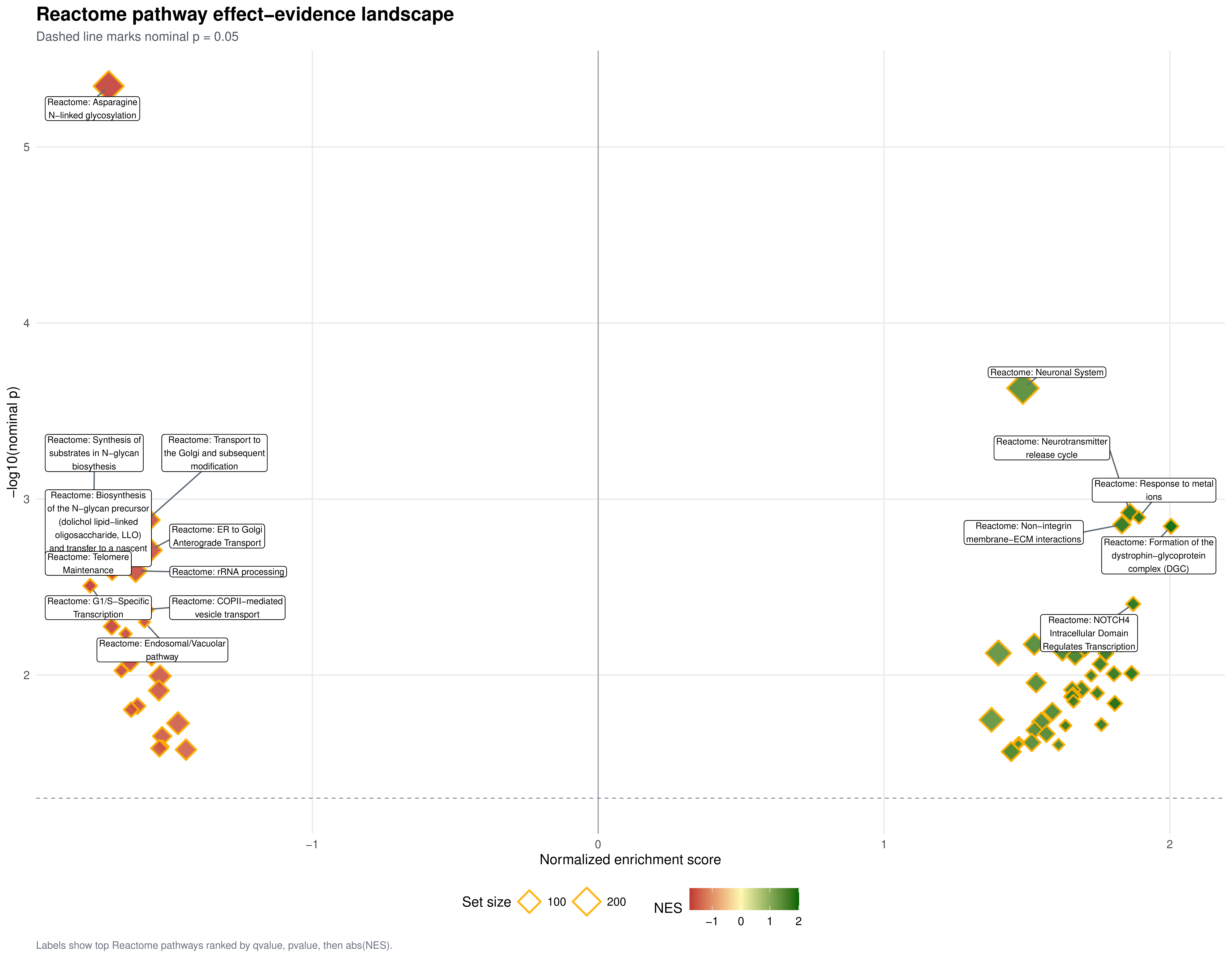

**Reactome Pathway Effect-Evidence Landscape.**

Reactome pathway effect-evidence landscape for CRH neurons comparing OUD patients with controls. Each point represents a Reactome pathway plotted by NES on the x-axis and -log10(nominal pvalue) on the y-axis. Point size represents Reactome pathway set size, and point fill represents NES direction and magnitude. The dashed horizontal line marks nominal pvalue = 0.05. Labeled pathways are the top-ranked terms by qvalue, pvalue, and absolute NES.

Supplementary Figure 54:

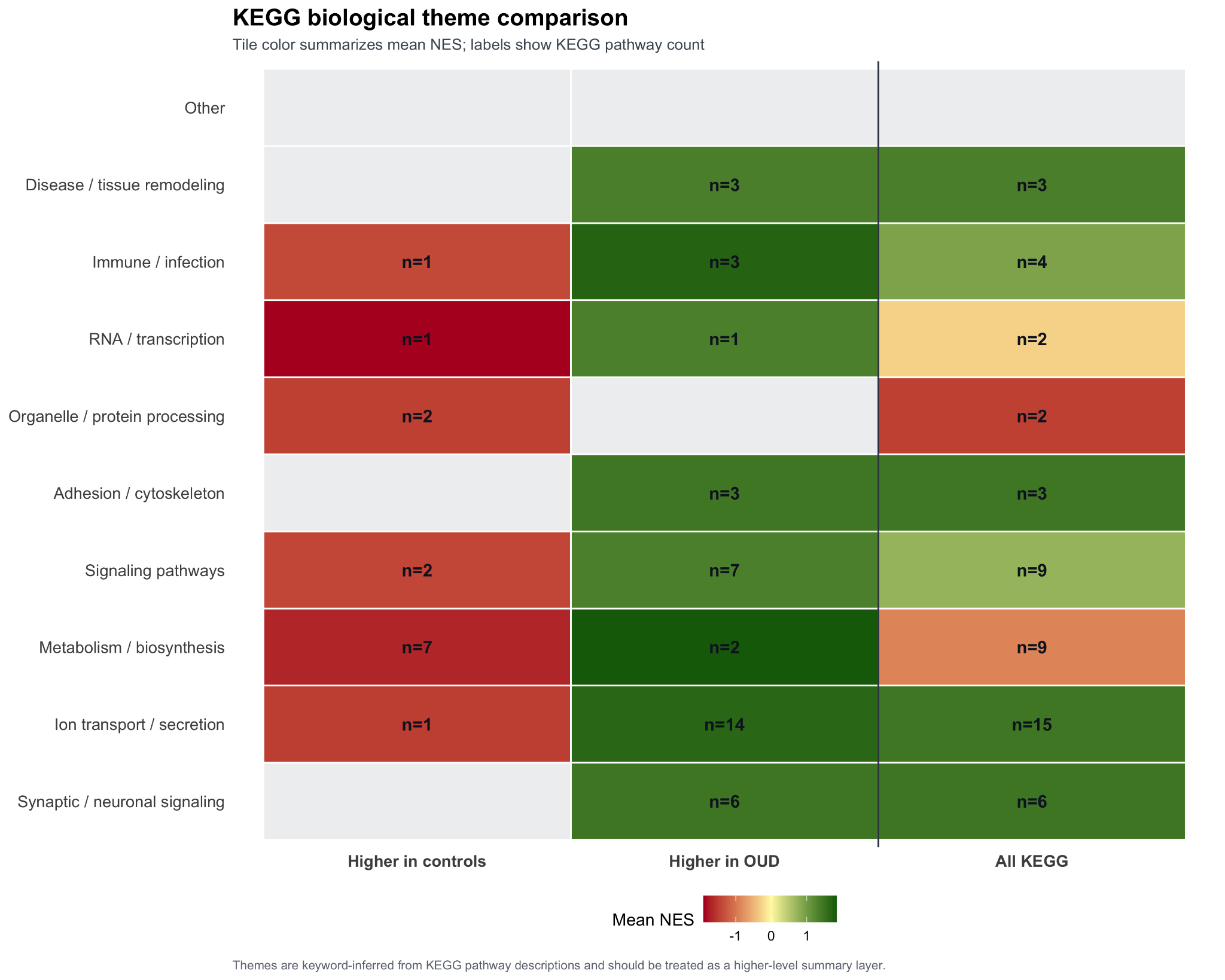

**KEGG biological theme comparison.**

Heatmap summarizing keyword-inferred KEGG biological themes by enrichment direction and overall KEGG mean NES. Labels show KEGG pathway counts per theme-direction bin, and tile color reports mean NES. Themes were assigned from KEGG pathway descriptions and should be treated as a higher-level summary layer.
